# Predicting future from past: The genomic basis of recurrent and rapid stickleback evolution

**DOI:** 10.1101/2020.11.13.382424

**Authors:** Garrett A Roberts Kingman, Deven N Vyas, Felicity C Jones, Shannon D Brady, Heidi I Chen, Kerry Reid, Mark Milhaven, Thomas S Bertino, Windsor E Aguirre, David C Heins, Frank A von Hippel, Peter J Park, Melanie Kirch, Devin M Absher, Richard M Myers, Federica Di Palma, Michael A Bell, David M Kingsley, Krishna R Veeramah

## Abstract

Similar forms often evolve repeatedly in nature, raising longstanding questions about the underlying mechanisms. Here we use repeated evolution in sticklebacks to identify a large set of genomic loci that change recurrently during colonization of new freshwater habitats by marine fish. The same loci used repeatedly in extant populations also show rapid allele frequency changes when new freshwater populations are experimentally established from marine ancestors. Dramatic genotypic and phenotypic changes arise within 5-7 years, facilitated by standing genetic variation and linkage between adaptive regions. Both the speed and location of changes can be predicted using empirical observations of recurrence in natural populations or fundamental genomic features like allelic age, recombination rates, density of divergent loci, and overlap with mapped traits. A composite model trained on these stickleback features can also predict the location of key evolutionary loci in Darwin’s finches, suggesting similar features are important for evolution across diverse taxa.

## Introduction

Can evolutionary outcomes be predicted? Biologists have long been fascinated with this question, including Darwin and Wallace’s anticipation of the existence of Morgan’s sphinx moth on the basis of orchid morphology (*1, 2*), Vavilov’s prediction of the types of morphological variants likely to occur in plants (*3*), and Gould’s gedankenexperiment about replaying the tape of life (*4*). Natural examples of recurrent evolution provide a particularly favorable opportunity to study the mechanisms that influence evolutionary predictability, including molecular patterns (*5, 6*).

While the predictability of evolution may appear in conflict with the unpredictability of historical contingency, understanding the past can yield important insights into future evolution. For example, vertebrate populations frequently harbor large reservoirs of standing genetic variation (SGV) (*7*) that give independent populations access to similar raw genetic material to respond to environmental challenges, as observed in diverse species including songbirds, cichlid fishes, and the threespine stickleback (*8–10*). SGV often originates in divergent species or populations where it is pre-tested by natural selection and then distributed by hybridization to related populations. Thus filtered and capable of leaping up fitness landscapes, SGV can also drive rapid evolution (*11*), helping address a very real practical challenge to testing evolutionary predictions: time.

Though systemic descriptions of evolution in real time have predominantly been restricted to invertebrates (*12*) and microbes (*13*), limited studies also exist in vertebrates, including guppies, finches, and stickleback (*14–16*). Rapid phenotypic evolution over decade-long time scales has enabled hypothesis testing against detailed observations at every step in the process, but this approach has yet to be extended to the genomic level.

At the end of the last Ice Age, threespine stickleback (*Gasterosteus aculeatus*) colonized and adapted to countless newly exposed freshwater environments created by the retreating glaciers around the northern hemisphere (*17*). This massively parallel adaptive radiation was facilitated by natural selection acting on extensive ancient SGV (*8*). Under the “transporter” hypothesis, these variants are maintained at low frequencies in the marine populations by low levels of gene flow from freshwater populations (*18*).

SGV enables new freshwater stickleback populations to evolve dramatically within decades (*16, 19, 20*), including conspicuous phenotypic changes in armor plates (*16*) and body shape (*21*). Little is known, however, about the underlying genomic dynamics early in adaptation, when evolution may be most rapid. We identify key molecular features that underlie repeated and rapid evolution of freshwater stickleback by comparing genomes from diverse extant populations with the earliest generation-by-generation changes in a detailed genomic time series from three newly founded populations. We identify several basic genomic and genetic features that can be used to predict evolutionary outcomes in stickleback, and show that they can predict genomic responses to selection in distantly related Darwin’s finches.

### Global resequencing and EcoPeak identification

Previous whole genome sequencing of threespine stickleback identified 174 loci covering 1.2Mb with alleles shared by common descent repeatedly selected in freshwater populations around the world (*8*). Just as human genetic diversity is greatest in Africa, where *Homo sapiens* arose (*22*), we hypothesized that the north Pacific region where stickleback originated (*17*) may contain a particularly rich pool of ancient adaptive alleles. To test this hypothesis, we generated whole genome sequence data with 76bp paired end Illumina reads for 38 new marine and 110 new freshwater stickleback, respectively (mean coverage of 8.1x). Combined with previous stickleback sequencing (*8, 23*), our dataset includes 227 individual genomes: 135 genomes from 70 northeast Pacific populations in Alaska, Haida Gwaii, British Columbia, and Washington and 92 genomes from 62 populations in California, Japan, and the Atlantic coasts of North America, Iceland, and northern Europe (Figure 1A).

**Figure 1:**
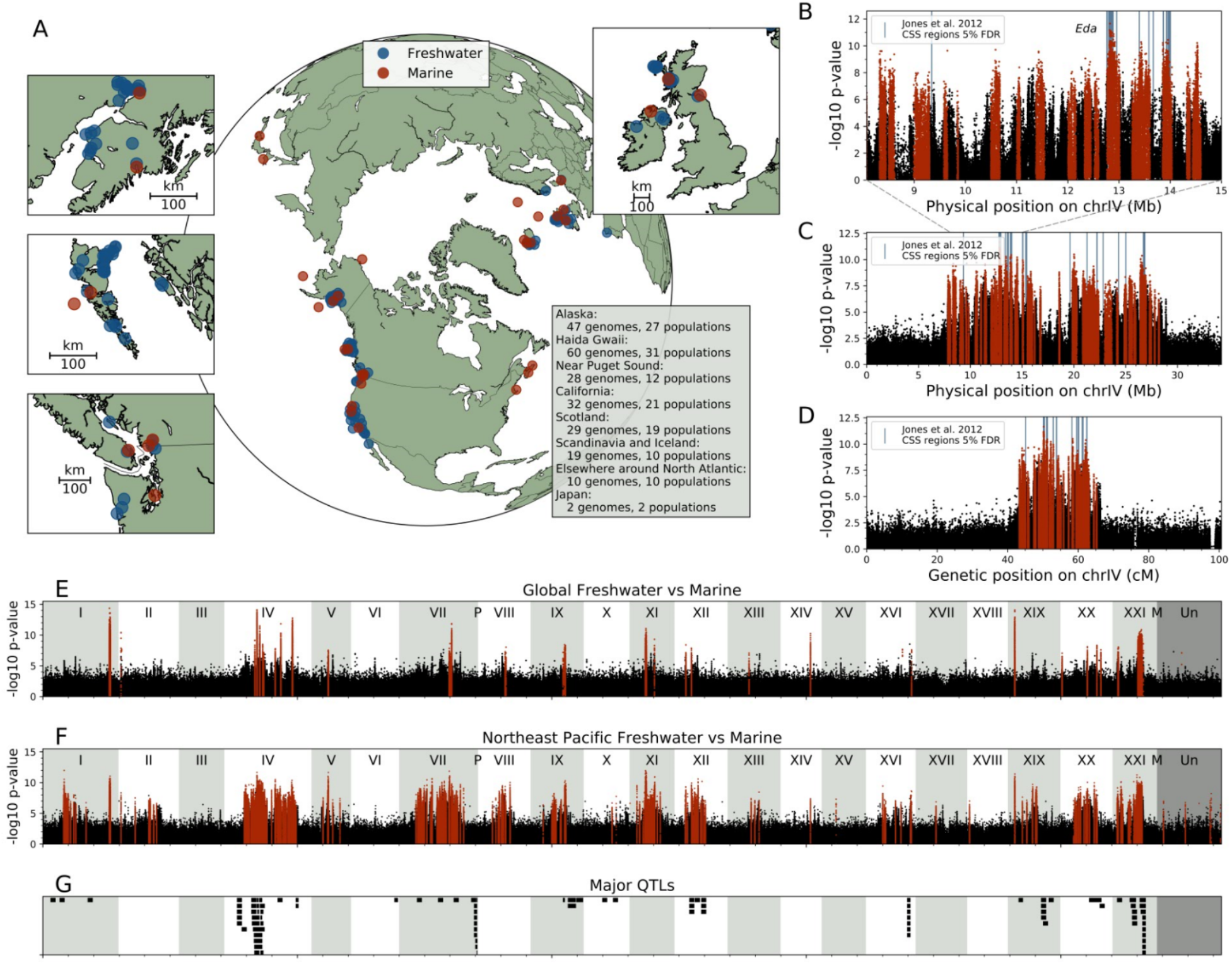
Recurrent peaks of ecological sequence differentiation between marine and freshwater stickleback from different regions of world. A) Marine (red) and freshwater (blue) stickleback from the locations shown were used for various analyses (Table S11.1). B) Detail of part of chrIV for SNP-based analysis of differential allele distribution between marine and freshwater ecotypes in the northeast Pacific basin. SNPs within specific-threshold EcoPeaks are red. A subset of regions overlap the globally shared peaks of marine-freshwater differentiation indicated by blue-colored bars (CSS 5% FDR identified by Jones et al. 2012). C) As in B, but for the whole chromosome (dotted lines from B to C). D) Same whole chromosome as in C, but with genetic (not physical) distance along the x-axis. E) & F) Genome-wide SNP divergence between marine and freshwater ecotypes globally and in the northeastern Pacific basin. G) Many differentiated regions overlap the location of major quantitative trait loci (QTL) controlling various morphological, physiological, and behavioral traits in previous genetic crosses (PVE > 20, interval < 5Mb from Peichel and Marques 2017).

We employed two methods to identify loci repeatedly differentiated in freshwater populations, both based on the expectation that unlike neutral loci, variants recurrently selected from standing variation will be more similar amongst geographically separated freshwater populations. First, we used a genetic distance-based approach within overlapping 2500bp windows tiled across the genome (as in Jones et al. 2012 (*8*)). While statistically powerful, this approach may miss younger loci with few differences between alleles and exhibits spatial resolution dependent on window size. Second, we analyzed the distribution of variants at individual bases across the genome, which has base-pair level resolution and less bias against younger loci, though at the cost of statistical power. After calling p-value-based peaks of ecotypic (freshwater- or marine-associated) differentiation using both methods, we accepted calls at two stringency levels, either requiring agreement between the two analyses at 1% FDR (specific) or support from either at 5% FDR (sensitive). We refer to these peaks of ecotypic differentiation as EcoPeaks. We called EcoPeaks for different geographic sets of samples to find alleles that were either shared globally, within the northeast Pacific, or within other geographic regions.

Although results of the global analysis largely matched previous reports (*8*), both the sensitive and specific call sets identified approximately five times as many Pacific EcoPeaks as global EcoPeaks, spanning seven-fold more of the genome (Figure 1E and 1F, Table 1). In addition, many novel EcoPeaks specific to the northeast Pacific exhibit even more consistent ecotypic differentiation (assessed by p-values) than others shared around the world (Figure 1B and 1C). Much smaller sets of novel EcoPeaks were identified in the North Atlantic, subglacial Pacific, and supraglacial geographic regions (Supplementary Table 1), consistent with other reports (*8, 24*).

**Table 1:** Overview of EcoPeaks and TempoPeaks. The Jones et al. (2012) comparisons are with the CSS 5% FDR set.

|  | **Global**<br>**EcoPeaks**<br>**(specific)** | **Pacific**<br>**EcoPeaks**<br>**(specific)** | **Pacific**<br>**EcoPeaks**<br>**(sensitive)** | **Cheney +**<br>**Scout** | **All young** |
| --- | --- | --- | --- | --- | --- |
| **# regions** | 39 | 209 | 212 | 86 | 344 |
| **Total bases**<br>**(%)** | 3.7 Mb<br>(0.78%) | 27.4 Mb<br>(5.82%) | 91.9 Mb<br>(19.53%) | 3.3 Mb<br>(6.95%) | 17.57 Mb<br>(3.73%) |
| **Median size** | 21.4 kb | 80.2 kb | 122.9 kb | 21.7 kb | 27.3 kb |
| **Recovery of**<br>**Jones et al.**<br>**2012 regions** | 86/174<br>(49.4%) | 112/174<br>(64.3%) | 158/174<br>(90.8%) | 47/174<br>(27.0%) | 98/174<br>(56.3%) |
| **Fraction in**<br>**Jones et al.**<br>**2012** | 18/39 (46.2%) | 29/209 (13.9%) | 33/212 (15.6%) | 10/86 (11.6%) | 24/344 (7.0%) |

As theoretical studies indicate that SGV is immediately available for evolution and may show an increased likelihood of large-effect alleles being advantageous compared to *de novo* mutations (*11, 25*), the rich genetic reservoir observed in the northeast Pacific provides a favorable system for studying the dynamics and predictability of rapid evolutionary change. Previous studies suggest that stickleback in the northeast Pacific can adapt to freshwater environments within decades (*19*). However, thus far studies have lacked temporal resolution of genome evolution in the critical early years of adaptation.

### Rapid contemporary evolution and TempoPeak identification

To characterize the earliest stages of evolution after the establishment of new freshwater populations, we analyzed annual samples from three lakes in Alaska that were recently founded by anadromous stickleback (Figure 2A). In 1982, stickleback in Loberg Lake (LB) were exterminated to improve recreational fishing (*16*). Sometime between 1983 and 1988, LB was invaded by completely plated (~33 plates/side) anadromous stickleback (most likely from neighboring Rabbit Slough, RS). The characteristic freshwater, armor-reduced phenotype increased rapidly from ~16% in 1991 to ~50% by 1995 and to ~95% by 2017 (*16*)(Figure 2B), with similarly rapid changes in overall body shape (*21*) and reproductive patterns (*26*). So as to more systematically examine even earlier generations of freshwater adaptation, Bell et al. (*27*) introduced ~3,000 anadromous RS fish into two other Cook Inlet lakes without outlets and similarly treated to exterminate fish: Cheney Lake (CH) in 2009 and Scout Lake (SC) in 2011. Low armor plated (~5-7 plates/side) stickleback began to appear in the 2nd and 3rd generation after founding in CH and SC respectively, and by 2017 they had increased to 20-30% (Figure 2B).

**Figure 2:**
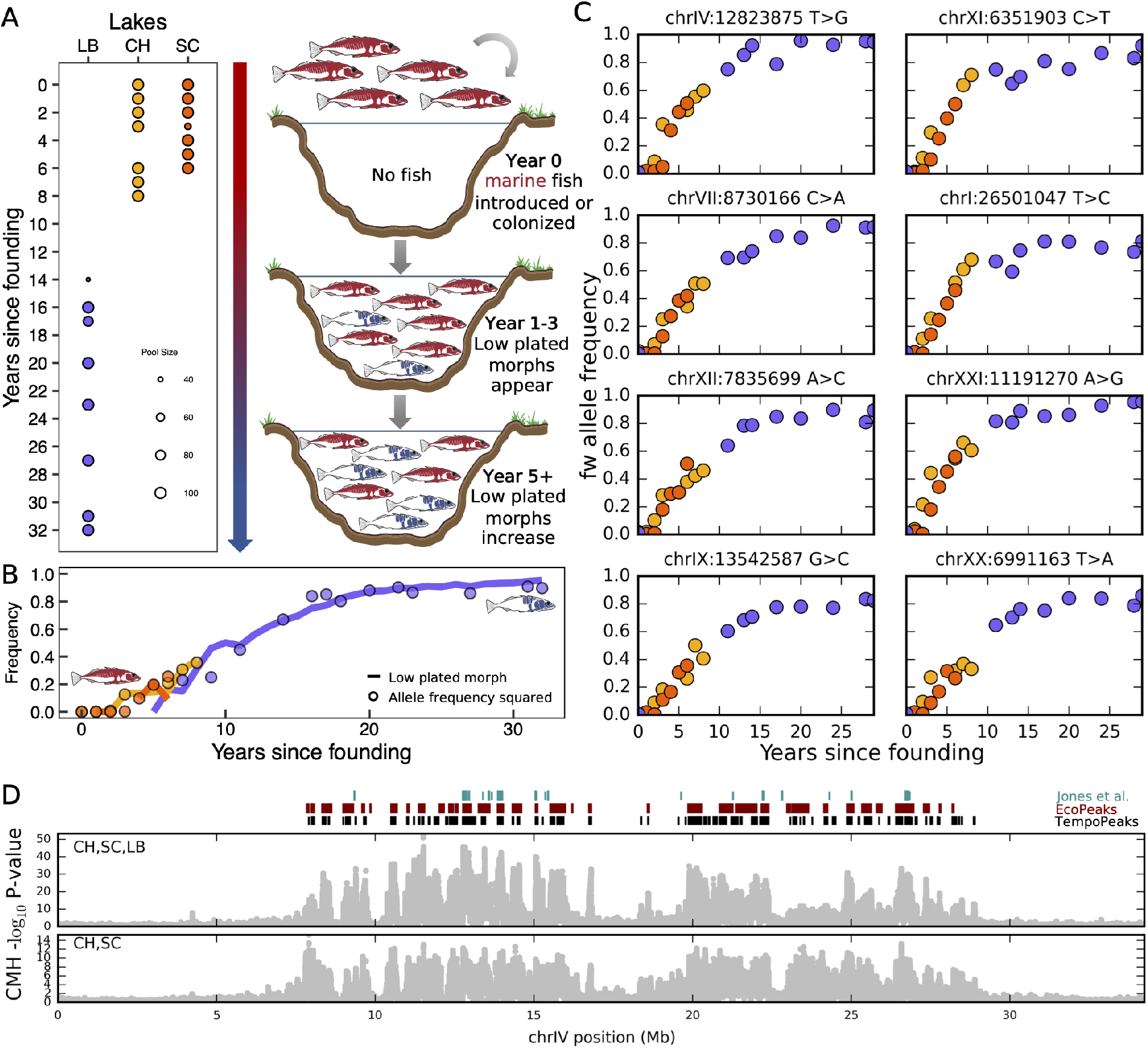
Contemporary evolution occurring in freshwater transplants in Cook Inlet, Alaska. A) The timing (years since founding) and approximate size of samples from lake populations (LB, CH and SC) founded recently by anadromous stickleback (left) and the scenario for divergence of anadromous populations after colonizing the lakes (right). Red and blue fish represent the complete armor plated and armor-reduced phenotypes, respectively. B) Frequency of armor-reduced phenotype across our CH, SC and LB time-series overlaid with the frequency squared for the freshwater *Eda* allele. LB data are based on a combination of individual genotypes and Pool-Seq frequencies, while CH and SC are based only on Pool-Seq frequencies. C) Allele frequency trajectories for 8 SNPs found within TempoPeaks on distinct chromosomes with the highest CMH scores (except for chrIV:12823875, the *Eda*-plate regulatory region SNP). D) Genome-wide distribution of window-based CMH scores across chrIV for different combinations of transplant lakes discussed in main text. Black, dark red and teal bars above figure represent CH+SC+LB TempoPeaks, northeast Pacific EcoPeaks and significant loci from Jones et al. (2012), respectively.

In order to obtain genome-wide allele frequencies across our time-series, we performed pooled whole genome sequencing (Pool-Seq) on all 7 available annual samples from CH and SC since founding and 8 from LB distributed between 1999 and 2017 (Figure 2A). Each freshwater Pool-Seq experiment consisted of 100 individuals (with three exceptions), with mean coverage of 223x per pool. In addition, we resequenced a pool of 200 RS individuals used to found the CH population in 2009 (RS2009) to 585x.

We identified SNPs with significant allele frequency change indicating directional selection using a modified Cochran–Mantel–Haenszel (CMH) test optimized for Pool-Seq data (*28*), followed by an approach analogous to our EcoPeak analysis to define both a sensitive and specific set of loci that we term TempoPeaks. Combining all three populations into a single CMH analysis (CH+SC+LB) and using RS2009 as a proxy for the founders of LB, we identified 524 sensitive and 344 specific TempoPeaks. Despite operating over very different time-spans, the visual correspondence between the EcoPeaks in long extant populations and the TempoPeaks in recently established populations is striking, particularly for the specific TempoPeaks, which have an 94% overlap with the sensitive EcoPeaks (Figure 2D). Even analyzing only CH+SC (thus focusing on <10 years of freshwater adaptation), we identified 271 sensitive and 86 specific TempoPeaks, 73% and 99% of which, respectively, overlap the sensitive EcoPeaks. This marked congruity strongly suggests that the ancient SGV represented by Pacific EcoPeaks is the primary genomic feature enabling extremely fast evolution of freshwater phenotypes in stickleback from the northeast Pacific basin.

The *Eda* SNP associated with armor plate variability (chrIV:12,823,875T>G (*29*)) is within the second most significant specific TempoPeak on chrIV. In both CH and SC the G allele increases rapidly from an initial frequency of <1 % to over 50% within 8 years, while approaching fixation in LB by 15 years. Notably, the square of G-allele frequencies (i.e. the expected number of GG homozygotes) tracks closely with frequencies of the low armor plate phenotype, consistent with almost complete recessiveness (*h*=0.0) for the G allele for this phenotype (Figure 2B). Nonetheless, in order to fit the allele frequency trajectory of this SNP, and in particular the extremely rapid increase in CH and SC, it was necessary to impose a dominance coefficient (*h*) of 1.0 along with a very large selection coefficient (*s*) of 0.55.

In fact, like *Eda*, most TempoPeaks display similarly sharp left-shifted sigmoidal allele frequency trajectories, indicative of very strong and dominant positive selection (Figure 2C). When modeling each peak SNP as independent we find an extremely high mean *s* of 0.30 (5th, 95th percentile 0.08-0.53) and *h* of 0.98 (5th, 95th percentile 0.95-1.0) for the 344 specific TempoPeaks found in CH+SC+LB. The estimated *s* values for chrIV, where there are 69 TempoPeaks, are particularly high (mean *s*=0.38), consistent with the accelerated evolution of this whole chromosome observed via an *F_ST_* analysis.

### Features associated with EcoPeak evolution

The remarkable speed at which northeast Pacific stickleback adapt to new freshwater environments suggests that analysis of EcoPeaks may provide unique insights into optimal genomic properties for evolution. Using *Gasterosteus nipponicus, Gasterosteus wheatlandi*, and *Pungitius pungitius* for calibration, we estimated molecular divergence time between a pair of freshwater (Little Campbell upstream) and marine (Little Campbell downstream) stickleback in windows tiled across the genome. We find that EcoPeaks as a whole are significantly older than the rest of the genome (1558 Ky vs 682 Ky, p <5e-324). Although peaks shared globally are typically older than those found just within the northeast Pacific, the imputed ages overlap considerably (Figure 3A). We estimate that the majority (161/209) are over a million years old and have passed through multiple freshwater colonization events during sequential ice ages.

**Figure 3:**
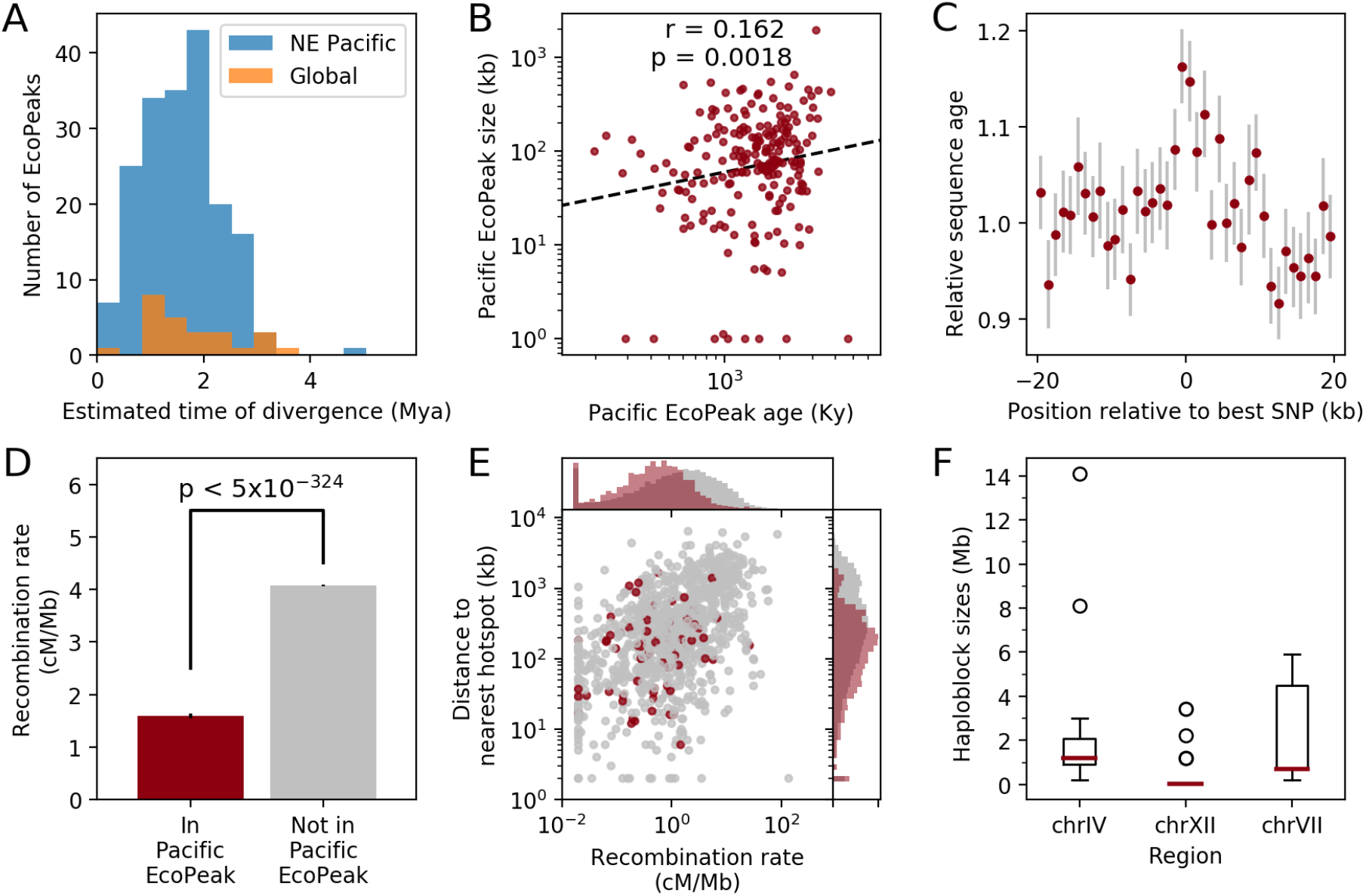
EcoPeak associations with age, region size, and recombination rate. A) Distribution of estimated molecular age for those EcoPeaks either shared worldwide (orange) or within the northeast Pacific (blue). B) EcoPeaks with older estimated molecular ages tend to be larger. C) Estimated ages decline with distance on either side of EcoPeaks. Each dot represents mean age in 1kb windows flanking the EcoPeak centers (grey bars, 1 SE). D) Recombination rates tend to be lower within EcoPeaks compared to the rest of the genome, +/1 SE. E) Recombination rates and distances to nearest 20x recombination hotspots, plotted for randomly subsampled 1kb windows tiled across the genome, with marginal histograms of all windows. Locations overlapping EcoPeaks (red) are shifted to both smaller hotspot distances and lower recombination rates compared to other genomic regions (grey). F) Observed haploblock size in marine fish carrying freshwater EcoPeaks on the indicated chromosomes across three marine populations. For all, specific northeast Pacific EcoPeaks are used.

Contrary to our expectations that recombination would disassemble regions over time, we found that older EcoPeaks are larger than younger ones (Figure 3B). This signature is strongest at the most significant markers within each EcoPeak, which are typically older than more distal sequences (Figure 3C). This suggests that individual regions may grow over time, with alleles originally based on an initial beneficial mutation accumulating additional linked mutations, snowballing over time to form a finely-tuned haplotype with multiple adaptive changes. This is consistent with work in other species identifying examples of evolution through multiple linked mutations that together modify function of a gene (*30–32*) and implies that progressive allelic improvement may be common.

We also observed that EcoPeaks frequently overlap major quantitative trait loci (QTLs) in stickleback (*33*) (Figure 1G, 73/209 overlaps observed vs 32/209 expected, p < 1e-15), suggesting that these variants underlie many mapped phenotypic traits. Just as the QTLs cluster in “supergene” complexes (*34*), so too do EcoPeaks (median observed interpeak distance 192kb vs 795kb expected, p = 4.88e-10). One particularly large complex (chrIV:8-17Mb) contains 22 EcoPeaks and the major QTLs controlling many different aspects of both defensive armor and trophic morphology (*e.g*., the length of dorsal and pelvic spines, the number of armor plates through *Eda*, gill rakers, and teeth). Thus, clustering may have important functional effects by allowing multiple traits and underlying EcoPeaks to be selected and inherited as a single unit, especially when in tight linkage. A fine-scale recombination map of RS stickleback (generated with LDHelmet (*35*)) shows that EcoPeaks are highly enriched in regions of low average recombination forming tightly linked haploblocks (Figure 3D, compare Figure. 1C, 1D). Interestingly, EcoPeaks are also enriched near local recombination hotspots within their neighborhood (Figure 3E), potentially facilitating reassembly of larger haplotypes upon freshwater colonization.

To further examine the frequency and size of haploblocks in individual fish, we surveyed 1643 stickleback from three Alaskan marine populations by SNP array genotyping. While most marine fish heterozygous for freshwater alleles carry a relatively small haploblock, some carry multi-megabase haploblocks containing multiple EcoPeaks (Figure 3F). Thus, a proper treatment of rapid stickleback evolution needs to account for the complex linkage of EcoPeaks rather than treating them independently.

### Modelling the genomic landscape of contemporary evolution

In order to estimate a more realistic distribution of fitness effects (DFE) that incorporates the genome’s recombination landscape, we developed a novel Deep Neural Network (DNN) approach that employs forward simulations. Our simulations, which are conceptually similar to those of Galloway et al. (*36*), attempted to replicate the dynamics of the “transporter model” (*18*), with one large (*N_e_*=10,000) anadromous population connected independently by gene flow to 10 smaller (*N_e_*=1,000) established freshwater populations. After 1,000 generations we founded three new freshwater populations, thus generating simulated allele frequency trajectories that reflect our annual LB, CH and SC samples (Figure 4A).

**Figure 4:**
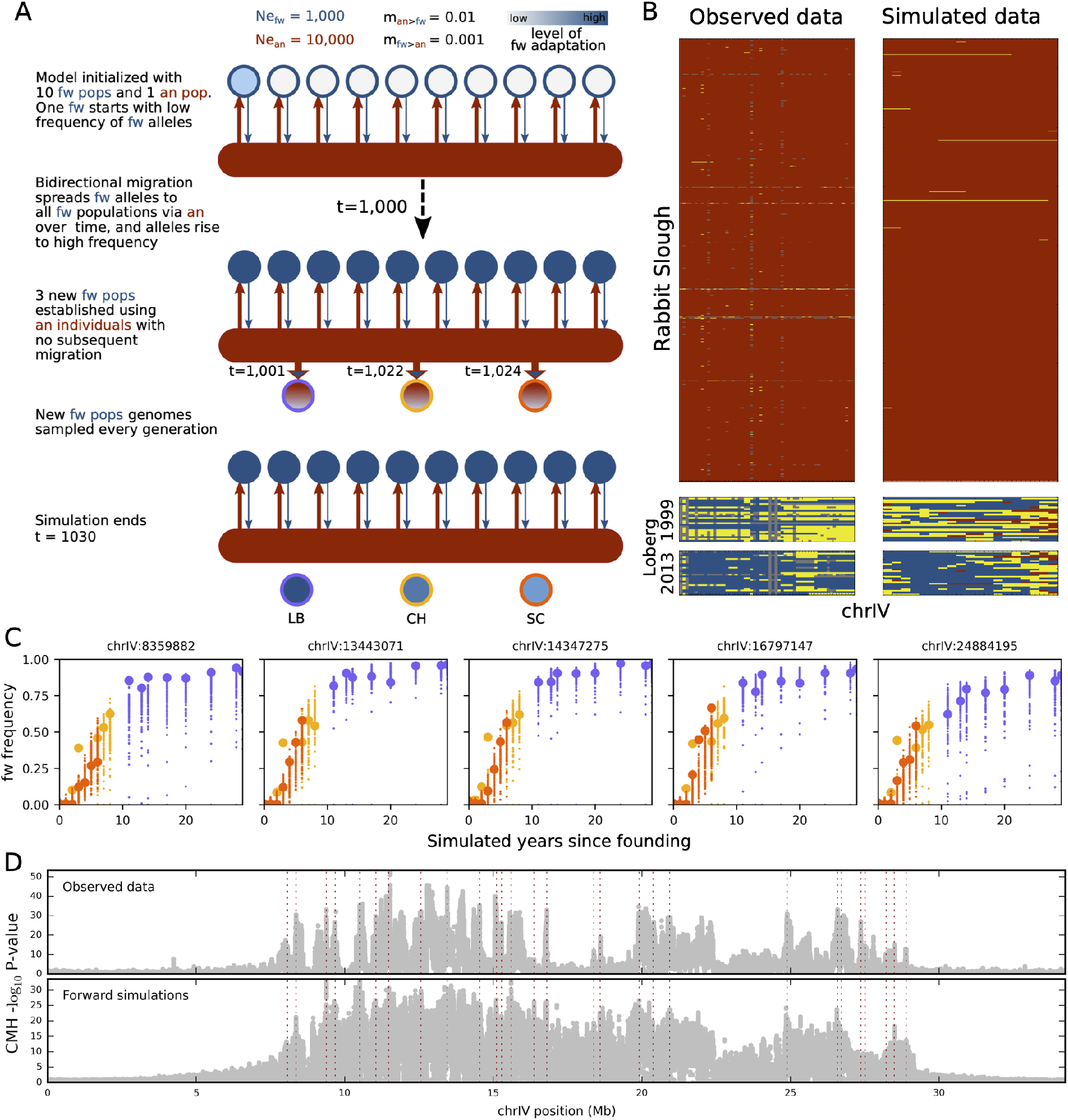
DNN simulation-based modeling of rapid and repeated stickleback evolution. A) Schematic showing evolutionary model of forward simulations under the transporter hypothesis. Red horizontal bars, anadromous (an) ancestor; blue circles, descendant freshwater (fw) isolates; and red to blue shaded circles, three adapting fw populations (i.e., LB, CH, SC) founded recently by an stickleback; and arrows, gene flow or founding events. B). Genotypes across chrIV for freshwater-associated SNPs in RS (n=750), LB in 1999 (n=25), and LB in 2013 for (left) observed and (right) simulated data under best fit DNN model. An homozygous, red; heterozygous, yellow; and fw homozygous genotypes, blue, respectively. C). Allele frequency trajectories for LB, CH and SC in 100 simulations under the best fit DNN model for 5 randomly selected SNPs.Observed data, larger points. D). Distribution of average CMH scores in windows of 2500bp across chrIV for (top) observed and (bottom) simulated data under best fit DNN model. Locations of SNPs under selection and used to fit DNN, red dotted lines.

Focusing our DNN analysis on a subset of 19 specific TempoPeak SNPs separated by ≥0.4cM (~100kb) along chrIV, we closely replicated observed allele trajectories of positively selected freshwater alleles across all SNPs simultaneously using a beta distribution-shaped DFE, for which the mean *s* across the 19 TempoPeaks was 0.063 and the standard deviation was 0.030, with reciprocal fitness costs implemented in the marine population (Figure 4C). The estimated *s* from our DNN was thus substantially smaller than the mean of 0.48 when each SNP was considered independently. In addition, 18/19 of the SNPs were predicted to be fully dominant and none fully recessive under the best model.

We validated our best fit DNN model by simulating the 19 selected TempoPeaks SNPs with the estimated DFE along with ~400k neutral SNPs distributed randomly along chrIV. Despite the neutral SNPs not being used in training the DNN, we were able to mimic the overall topology of the CMH scores across the entire genome, suggesting that our model was capturing the overall genomic architecture of freshwater adaptation (Figure 4D). Our best fit DNN model also appeared to recapitulate much of the haplotype structure of the array data from individuals from RS, LB1999 and LB2013 (Figure 4B). Notably, the transition to freshwater alleles appears to be somewhat slower on the right half of chrIV, where there are fewer EcoPeaks, TempoPeaks and QTLs.

Overall, our model suggests that extremely rapid and replicable allele frequency increases on chrIV in LB, CH and SC are mostly driven by multiple linked (primarily) dominant alleles, each with relatively smaller *s* values that act in concert, with recombination hotspots between them allowing rapid reassembly of optimum freshwater haplotypes, consistent with the transporter hypothesis. The lower individual *s* values may allow these dominant alleles to persist in the marine environment at low frequency after being disassembled by recombination, especially if some act in epistasis.

### Biological features with predictive power

Given the genome-wide dynamism of the earliest stages of freshwater adaptation, we attempted to identify genomic features that predict the speed of evolution at TempoPeaks and understand why some peaks are selected more rapidly than others. We used CMH scores as a proxy of evolutionary speed for each TempoPeak in CH+SC+LB and regressed these against a variety of sequence features.

Interestingly, the best predictor for the speed of evolution is the degree of ecotypic differentiation between marine and long-established freshwater populations (Pacific EcoPeak p-value), with variants more commonly differentiated in the northeast Pacific and around the world being selected more quickly (Figure 5A). Fisher’s geometric model indicates that alleles with large effects are usually disfavored; however, the “prefiltering” of ancient SGV that counters this tendency (*11*) largely benefits alleles that are broadly positively selected, possibly explaining this result.

**Figure 5:**
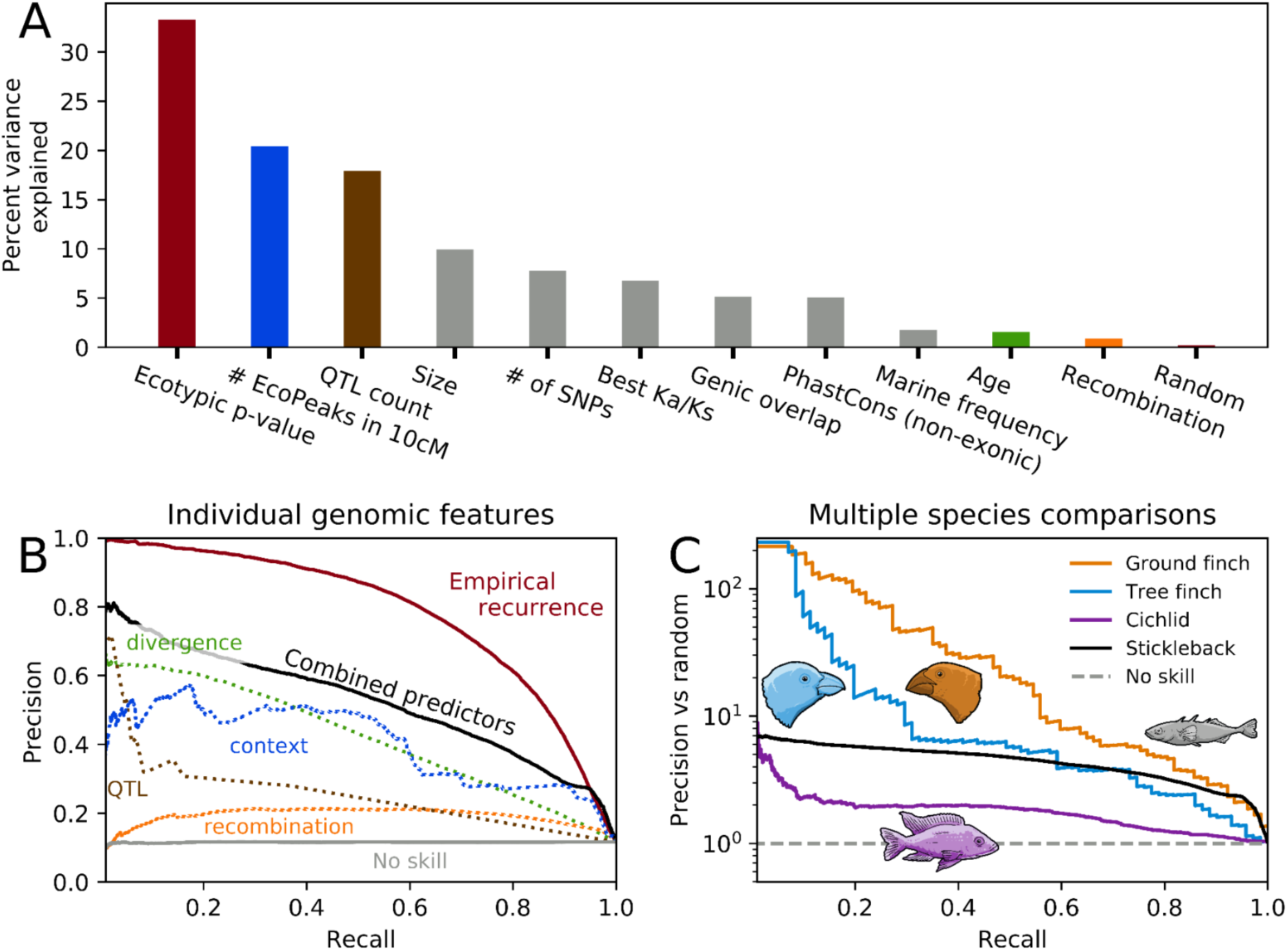
Properties underlying speed and locus of selection in stickleback, cichlids, and Darwin’s finches. A) Variance in the speed of TempoPeak selection explained by different underlying genomic features, including, colored bars: consistency of marine-freshwater differentiation (peak ecotypic p-value), number of additional EcoPeaks within 10cM, number of major QTLs overlapped, estimated allelic age, and recombination rate; and, grey bars: genomic size of EcoPeak, elevated Ka/Ks in coding regions, overlap with genic sequences, overlap with conserved noncoding sequence (PhastCons non-exonic), and carrier frequency of freshwater alleles in marine populations. B) Precision-recall curve for predicting the locations of selected loci in CH+SC+LB lakes by individual genomic features (dotted lines), a composite model trained with these basic predictors, or the empirical expectation of recurrence based on many extant populations. C) Performance above chance of the composite model applied to stickleback, cichlids, and two representative pairs of species of Darwin’s finches (ground finches: *G. magnirostris* vs *G. propinqua*; tree finches: *C. pauper* vs *C. psittacula*).

We also found that larger TempoPeaks are typically selected more rapidly. Similarly, greater TempoPeak density predicts more rapid divergence, suggesting that our simulation accurately reflects how nearby loci mutually reinforce their collective selection. Overlap with major QTLs also has a strong association with rapid evolution, while other variables such as age, recombination rate, genic overlap, sequence conservation, Ka/Ks, and ancestral marine frequency have smaller contributions to predictive power (Figure 5A).

We also tested whether underlying sequence characteristics could predict not just the speed of selection in CH+SC+LB, but also the location of the selected regions themselves. We find that recombination rate, QTL overlap, allelic age, and an integrated genomic context score (see Supplement) of the previous features are all useful predictors (Figure 5B). By combining these fundamental features into a logistic model trained on the survey of extant populations, the most confident predictions of selected regions in the rest of the genome achieve 85% precision. This model performs 67% as well as predictions based only on empirical repeatability in extant populations (Figure 5B). Thus, our understanding of underlying principles reflects an incomplete yet substantial proportion of evolutionary repeatability.

### Parallels in distant species

To test the generality of these predictive factors, we applied the stickleback-trained model to a dataset of 12 pairs of species of Darwin’s finches (*37*). Darwin’s finches have undergone adaptive radiation in the Galápagos Islands over the last several hundred thousand years, are ~435 million years divergent from stickleback, and face very different selective pressures. As in stickleback, however, the “islands of divergence” of all 12 analyzed pairs of species of Darwin’s finches (*sensu* Han et al. 2017) are enriched for ancient alleles overlapping mapped QTLs with low recombination rates. The top 100 windows predicted by the stickleback model recover a median of 28-fold more previously identified islands of divergence than expected by chance (Figure 5C, p< 1e-10), including the *Alx1* and *Hmga2* loci implicated in beak morphology in multiple species pairs (even without QTL input). The model also recovers a significant proportion of differentiated loci in a recent case of cichlid speciation (*38*). Thus, a handful of basic genomic properties allow strong quantitative predictions of the location of key evolutionary loci, even across widely separated branches of life.

## Conclusion

The importance of SGV for evolution is becoming increasingly apparent, especially in species with large genome sizes (*39*), including humans (*40*). At first glance the dependence of threespine stickleback on SGV for freshwater adaptation may appear to be a peculiarity in terms of repeatability and speed and their particular natural history. However, by more comprehensively understanding the dynamics of this highly optimized process, we have extracted general features of genome architecture and evolution that successfully translate to species on distant branches of the tree of life, thus demonstrating the tremendous power of the stickleback system to identify unifying principles that underlie evolutionary change.

## Acknowledgements

We thank field assistants, numerous researchers who provided samples, and many others who contributed to this work. (See online supplementary information.)

## Author contributions

MAB, experimental design, Alaskan sampling and population founding, tissue sampling, morphological data collection. PJP, Alaskan sampling and population founding. FAvH, Alaskan sampling, logistical support. WEA, Alaskan sampling, morphological data collection. DCH, Alaskan sampling. GARK, analysis of extant populations and genomic properties. DNV, MM, TB, DNA extraction, quantitation, selection, and pooling of Alaskan samples. FCJ, design of extant sampling, SNP-array. HIC, transfer and visualization of genome annotations. DMK, experimental design and conceptual guidance. KRV, KR, analysis and modeling of contemporary populations. GARK and KRV wrote the manuscript with input from all authors.

