## Supplementary Material for "Predicting future from past: The genomic basis of recurrent and rapid stickleback evolution"

Roberts Kingman et al.

#### TABLE OF CONTENTS

1. ACKNOWLEDGMENTS
2. IACUC APPROVALS
3. DATA ACCESSIBILITY
4. SAMPLE COLLECTION
5. DNA EXTRACTION AND QUANTIFICATION
6. POOLING CONTEMPORARY EVOLUTION DNA SAMPLES
7. GENOME SEQUENCING
8. ARRAY GENOTYPING
9. REFERENCE GENOME CONSTRUCTION
10. BIOINFORMATIC PROCESSING & VARIANT CALLING
11. ECOTYPE GROUPING OF GEOGRAPHIC POPULATIONS
12. ECOPEAK IDENTIFICATION
13. HYBRID ZONE ANALYSIS
14. ALLELIC AGE ESTIMATION
15. FREQUENCY AND HAPLOTYPE STRUCTURE FROM ARRAY GENOTYPES
16. COMPARING WGS, ARRAY AND POOL-SEQ ALLELE FREQUENCY ESTIMATES
17. RECOMBINATION MAP CONSTRUCTION
18. $F_{ST}$ ANALYSIS OF CONTEMPORARY EVOLUTION SAMPLES
19. NE INFERENCE FOR CONTEMPORARY EVOLUTION LAKES
20. CMH ANALYSIS OF CONTEMPORARY EVOLUTION SAMPLES
21. TEMPOPEAK IDENTIFICATION
22. COMPARING TEMPOPEAKS AND RECOMBINATION HOTSPOTS
23. ESTIMATING TEMPOPEAK SELECTION COEFFICIENTS ASSUMING SNP INDEPENDENCE
24. ESTIMATING TEMPOPEAK SELECTION COEFFICIENTS FOR CHRIV USING A DEEP NEURAL NETWORK APPROACH
25. PREDICTING SPEED OF STICKLEBACK ALLELE FREQUENCY CHANGES
26. PREDICTING LOCI OF SELECTION IN STICKLEBACK, FINCHES AND CICHLIDS
27. REFERENCES

### 1. ACKNOWLEDGMENTS

We thank the many individuals who contributed to this study:

*Fish samples used for geographic surveys:*
In addition to authors of this study, the following people contributed to the sample collections: Steve Arnott, Ben Blackman, Frank Chan, Pam Colosimo, Anne Dalziel, Bruce Deagle, Dánjal Petur Højgaard, Jan Arge Jacobsen, Bjarni Jonsson, Richard King, Dietmar Kueltz, Andrew Maccoll, Jeff McKinnon, Craig Miller, Seiichi Mori, Kristin O'Brien, Catherine Peichel, Mark Ravinet, Melissa Rhodes-Reese and NOAA, Tom Reimchen, Jonathon Richmond, Dolph Schluter, Mike Shapiro, Brian Summers, and Tim Vines.

*Undergraduate lab assistants.* James Ancona,  Hillary Babalola, Priya Chohan, Jonathan F. Gaige, Zain Khan, Jared Mallozi,

*Sampling access.* Mike Tauriainen and the staff of T & J Gravel Products permitted us to collect stickleback from their property on Scout and Loberg lakes, respectively.

*Field assistants.* Alaska: Stephen Abrams, Daniel Arciari, David B. Bell, Madeline R. Bell, Sam R. Bell, Bjorn Berland, Melanie Bobb, Glenn A. Bristow, Kaitlin T. Ellis, Anup K. Gangavelli, Megan A. Hahn, Alan C. Havens, Adam Hernandez, Jill Johnson, Erica Kalabacas, Anjali Karve, Freda Kreier, Ryan Lucas, Alice McGarry, Matthew McGee, Brian K. Lohman, Ryan Paitz, Anastasia Plaunova, Jennifer L. Rollins, Heinrich Schultz, David L. Soltz, Laura Stein, Abbey C. Thompson, Matthew P. Travis Heide Viitaniemi, Julia I. Wucherpfennig, Kathleen T. Xie

We thank Angie Hinrichs, Hiram Clawson, Kayla Smith, and Donna Karolchik for their contribution to the UCSC Stickleback Genome Browser annotations for *gasAcu1*, which were lifted to *gasAcu1-4* for this study.

#### 2. IACUC APPROVALS

Veeramah: 2019 - NF - 6.17.22 - FI, 2020 - R1 - 6.17.22 - FI, Stony Brook University

Bell: 237429-19, Stony Brook University

Kingsley: #13834, Stanford University

Jones: 35/9185.82-5 EB01/09 A, Baden-Württemberg Regierungspräsidium, Germany.
Friedrich Miescher Laboratory of Max Planck Society, Tübingen, Germany.

#### 3. DATA ACCESSIBILITY

All whole genome sequencing Illumina data of extant populations has been deposited in the Short Read Archive at accession PRJNA247503.

All whole genome sequencing Illumina data for contemporary Pool-Seq experiments has been deposited in the Short Read Archive at accession PRJNA671824.

All whole genome sequencing BGI data for Rabbit Slough 2009 genomes used to construct the recombination map has been deposited in the Short Read Archive at accession PRJNA671690.

SNP Genotyping Array data has been deposited on Data Dryad (DOI https://doi.org/10.5061/dryad.pvmcvdnjm)

EcoPeak, TempoPeak, and Rabbit Slough recombination rate data can be visualized and downloaded (via the Table Browser (*1*)) of the UCSC Genome Browser (http://genome.ucsc.edu/) (*2*) by copying the following track hub (*3*) URL into the “My Hubs” tab at https://genome.ucsc.edu/cgi-bin/hgHubConnect: https://sbwdev.stanford.edu/kingsleyAssemblyHub/hub.txt.

To convert between *gasAcu1-4* and the original *Broad S1* stickleback genome assembly, we provide the following liftOver chains:

***gasAcu1-4 to gasAcu1 liftOver chain***
https://drive.google.com/file/d/1k0cnix3iq8sgSbpLW_bxH5XsgY73QcDX/view?usp=sharing

***gasAcu1 to gasAcu1-4 liftOver chain***
https://drive.google.com/file/d/1w4PJezJYGC1I-VhFHarcJ20cZg7oSHeT/view?usp=sharing

***gasAcu1-4.2bit***
https://drive.google.com/file/d/1cHzWj2dQdi-5TUhrZnjVAu3En_iqa7MU/view?usp=sharing

These files will be available from Data Dryad upon publication at the following location: https://doi.org/10.5061/dryad.547d7wm6t

#### 4. SAMPLE COLLECTION

*General sampling methods for established populations*. Fish were generally trapped using unbaited minnow traps set near the shore, immediately euthanized with MS 222 (tricaine methanesulfonate), then preserved in 70% ethanol for transport and storage.

*Sample collections for SNP Genotyping array.* 751, 655 and 237 marine stickleback fish sampled from Rabbit Slough, Resurrection Bay, and Glacier Spit, Alaska respectively, were used for genotyping with a custom SNP genotyping array.

*General sampling methods and background for contemporary evolution lakes.* Annual samples for evolutionary time series to study rapid genomic evolution were made from populations in three Alaskan lakes, Cheney, Scout, and Loberg (https://www.adfg.alaska.gov/index.cfm?adfg=fishingSportLakeData.main&StockingAreaID=2 for lake information, **Fig S4.1**). These populations had been founded recently by anadromous (sea-run) threespine stickleback, and genetically determined phenotypic traits (*4*, *5*) were evolving rapidly in all of them (*6*–*10*). Stickleback were sampled under collecting permits from the Alaska Department of Fish and Game and sacrificed under protocols approved by the Institutional Animal Care and Use Committee of Stony Brook University to MAB. Each sample was trapped using unbaited Gees® Galvanized Minnow Traps (G-40), which were set for about 24 h near shore, on the bottom, at about 1 m depth. Fish were sacrificed using an overdose of MS 222 (Tricaine methanesulfonate) until they were unresponsive to a tap on the side of the bucket. They were rinsed in lake water and immediately immersed in 100 to 70% ethanol. The ethanol was replaced with fresh ethanol within 24 hours, and the sample was stored at room temperature. Sample details are in **Table S5.1**. The anadromous Threespine Stickleback population that breeds in Rabbit Slough (*11*), about 1.25 km from Loberg Lake, or a closely related population from an adjacent tributary in the same drainage probably founded the Loberg Lake population naturally (*6*). MAB introduced anadromous Rabbit Slough adults (*7*) into Cheney and Scout lakes. Thus, all three lake populations that we used to study contemporary genomic evolution were derived from the same or closely related anadromous threespine stickleback populations.

*Rabbit Slough samples.* Two thousand nine hundred sixty-four (2964) adult anadromous (sea-run) Threespine Stickleback were trapped in Rabbit Slough, Matanuska-Susitna Borough, Alaska as they migrated upstream to breed in 2009 (*7*). They were captured using nine one quarter-inch mesh traps, set entirely across the outlet of a culvert that discharges from under the west side of the Parks Highway. The first dorsal spine was clipped from each specimen before it was released into Cheney Lake, and the spines were immediately put into 95% ethanol.

*Loberg Lake samples.* Loberg Lake is a 4.5 ha lake near Palmer, in the Matanuska-Susitna Borough, Alaska at about 61.56° N latitude and 149.26° W longitude. The population was founded between 1983 and 1988, and we treat 1985 as the year it originated (*6*). It was apparently founded by a substantial number of anadromous threespine stickleback that entered the lake through a spring that discharges into Spring Creek, which is in the same drainage and close to Rabbit Slough (*6*). Annual samples from Loberg Lake were first made in 1990, and subsamples were preserved in ethanol for DNA sequencing starting in 1999 (*6*, *10*). Most of these samples came from a site about halfway between sites A and C of Bell et al. (*6*), but some came from the five other sampling sites around the lake. Samples were made with six to 20 one-eighth and one-quarter-inch mesh traps.

*Cheney Lake introduction and samples.* Cheney Lake is a 9.9 ha lake in Anchorage City-Borough, Alaska at 61.2° N. latitude and 149.76° W longitude (*7*). It was treated with rotenone to exterminate an exotic species in the fall of 2008 and between 29 May and 3 June 2009, 2964 adult anadromous stickleback were transported in their native water in aerated coolers (i.e., ice chests) from Rabbit Slough to the University of Alaska Anchorage (UAA), 55 km away. Their first dorsal spine was clipped for DNA extraction. They were held in aerated, aged tap water, and transported within 24 hours to Cheney Lake, about 5 km away, in aerated, aged tap water in a cooler, and released at the south end of the lake. The water in the cooler was exchanged with Chaney Lake water in a series of dilutions to reduce the temperature difference to <4 C. The stickleback were caught with large aquarium nets and released into the lake. They formed a linear school and swam rapidly into deeper water.

Annual sampling started in Cheney Lake in the late summer of 2009, when juvenile stickleback were abundant. Capture efficiency declined during the next two years, and virtually no stickleback were caught in 2013 and 2014. In 2015, they were caught in small numbers, which have increased progressively since then (*7*). Since 2015, traps have been set only at the tip of a peninsula on the north shore, where capture was most efficient. Samples were available for DNA extraction each year besides 2013 and 2014.

*Scout Lake introduction and samples.* Scout Lake is a 38.5 ha lake in Stirling, Kenai Peninsula Borough, Alaska at 60.54° N latitude and 150.83° W longitude (*7*). Between 4 June and 4 July 2011, 3047 adult stickleback from the Rabbit Slough anadromous population were released into Scout Lake using the same methods as in Cheney Lake. However, they were not abundant and had to be accumulated for several days at UAA before transport to Scout Lake, about 216 km away, and released at the east end of the lake. Annual samples were first made from this population in the late summer of 2011. Capture efficiency declined precipitously in 2013 and 2014, and their numbers have been recovering since then (*7*). Annual samples for DNA extraction have been made each year since.

*Tissue sampling.* Each sample was given code letters and a descriptive name, and each fish was individually numbered. Eight stickleback at a time were removed from the storage bottle and a pectoral or caudal fin (occasionally two fins from small fish) was grasped with needle-nose forceps and cut off with iridectomy scissors, which were washed in alcohol between specimens and in bleach between samples from different populations or dates. Fin clips were stored in 70% ethanol in 1.5 ml microcentrifuge tubes or 48-well plates. The fish from which fins were cut were stored in labeled 10 ml screw-cap tubes with two fish per tube (head up, head down). Fish from which a fin was not clipped were left in the storage bottle. The plates of fin clips were sealed in Saran Plastic Wrap® and taped shut for storage and shipment. Fin clips were made in DMK's laboratory and shipped in a cooler to KRV's laboratory by MAB. All ethanol-preserved stickleback specimens are stored in DMK's laboratory in a cold room.

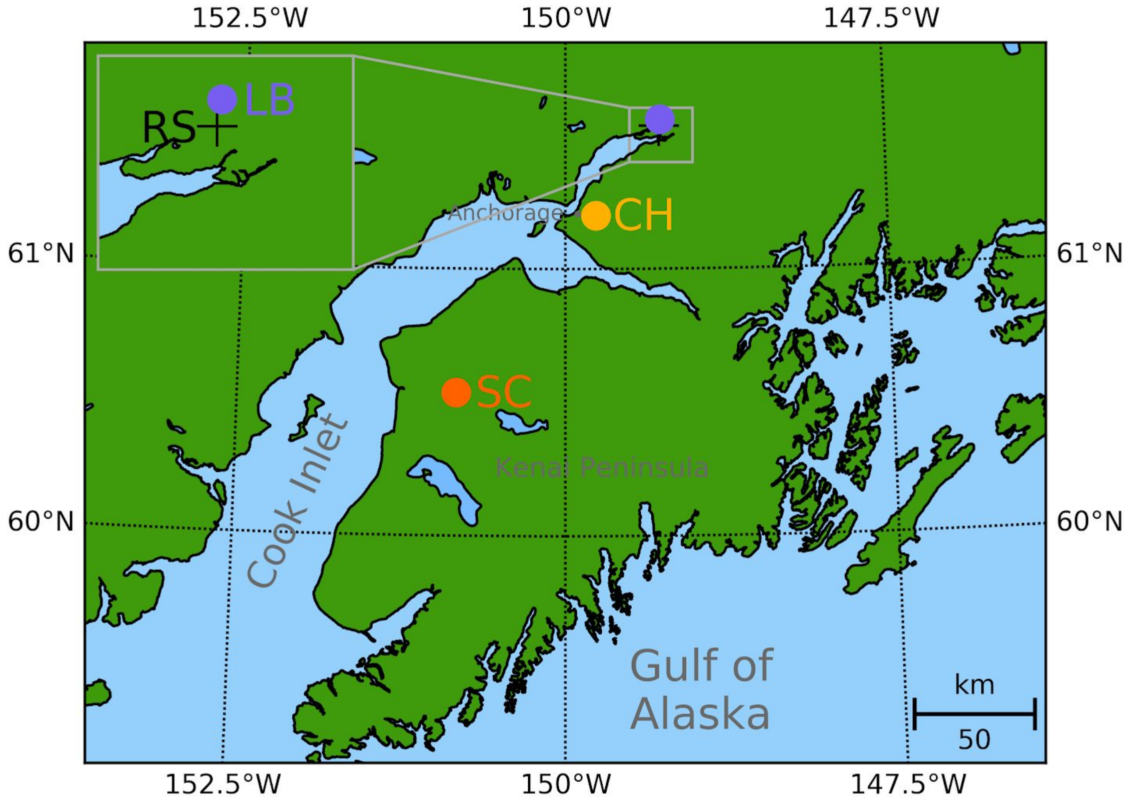


**Fig S4.1:** Map of Alaska showing location of three contemporary freshwater lakes as well as Rabbit Slough.

#### 5. DNA EXTRACTION AND QUANTIFICATION

*Geographic population survey samples.* DNA from caudal and/or pectoral fin tissues was extracted using overnight proteinase K digestion followed by phenol:chloroform isolation with PhaseLock tubes as previously described (*12*). DNA concentration was quantified using a NanoDrop spectrophotometer.

*SNP genotyping array samples.* DNA from SNP genotyping array samples were extracted from clipped spines or fins using overnight proteinase K digestion followed by phenol:chloroform isolation as previously described (*12*). DNA concentration was quantified using a NanoDrop spectrophotometer.

*Contemporary evolution samples.* DNA from contemporary samples was extracted from clipped fins or spines using the DNeasy 96 Blood & Tissue Kit following the standard “animal tissue” protocol. Tissue was digested with proteinase K for 24-48 hours, depending on the observed state of the digest. In general only one lake time point was used per 96-well plate to avoid cross-contamination between lake time-point. RNase A was either applied to individual samples during these extraction or to samples after pooling (see **Table S5.1**). DNA for each individual sample was quantified using the Qubit High-sensitivity DNA assay.

#### 6. POOLING CONTEMPORARY EVOLUTION DNA SAMPLES

DNA from annual Cheney, Scout, or Loberg lake time points were pooled at equimolar concentrations using an Opentrons OT-2 robot. Due to the limitations of the OT-2 P10 pipette's accuracy, we could only confidently pipette volumes greater than or equal to 3 ul. Given the spread of DNA concentrations across samples from a given lake time point, in most cases this made it impossible to make a single pool from all chosen samples. Thus, in most cases, two or three subpools were generated and merged into a single, final pool. The two exceptions were the Loberg 1999 and Loberg 2002 pools where we were able to directly pool all chosen samples into single pools.

For each lake time point, samples for pooling were primarily chosen from the set of successful extractions for said time point at random. Given that the final concentration of the pools for sequencing needed to be above or around 25 ng/ul, the least concentrated samples often needed to be excluded from random sample selection. Additionally, in certain cases, extractions with extraordinarily high concentrations (relative to other extractions from the same lake time point) were also excluded from sample selection due to the inability to accurately pipette the lower volumes necessary to pool these samples.

Chosen samples would be sorted in ascending order of DNA concentration (ng/ul). The least concentrated samples per subpool would have 10 ul of extract included in the subpool. Each subsequent (more concentrated) sample would have a smaller volume of extract (ul) containing an equivalent amount (ng) of DNA added to the subpool. This would continue until the pipetted volume reached our cut-off volume (which varied between 3 and 5 ul from pool to pool). At that point, a new subpool would begin and the process would begin again. Subpools would be labeled A and B (and when necessary C) with the A subpools being the least concentrated. The subpool DNA concentrations were assessed using the Qubit dsDNA HS Assay Kit to confirm that concentrations were consistent with the calculated expected values (based on the concentrations of the constituent samples). Subpools would then be equimolarly merged based on their concentration and the number of samples each subpool contained. Aliquots of the B and C subpools were diluted down to the same concentration as the A subpool. Volumes proportional to number of samples in the subpool would then be taken from each aliquot to constitute the final pool. Pools with extensive RNA contamination would then also be treated with RNAse A and cleaned up using a standard ethanol precipitation. Additionally, due to two Qiagen plates being switched, we believe that one freshwater individual sample (from Arc Lake, a lake not analyzed in this study) may have been accidentally included in the Rabbit Slough 2009 pool; however, given that there were 199 actual Rabbit Slough samples in this pool, any effects would be minimal.

#### 7. GENOME SEQUENCING

*Geographic population survey whole genome sequencing.* Genomes of fish sampled throughout the species range were sequenced using 2x76bp Illumina paired-end sequencing libraries. Individually barcoded genomic libraries were prepared following standard Illumina TruSeq protocols using DNA extracted from caudal and/or pectoral fin tissue. Libraries were sequenced on an Illumina HiSeq2000 at the Broad Institute to a mean coverage depth of 5.5X.

*Contemporary evolution population Pool-Seq.* Genomic libraries were prepared from pooled DNA from each individual lake time-point by the New York Genome Center (NYGC) and underwent paired-end re-sequenced with read lengths of 150bp either on a single Illumina NovaSeq 6000 S4 flowcell (n=16) or on individual lanes of an Illumina HiSeq 2500 (n=8), dependent on the level of DNA fragmentation and library complexity observed from pre-screening (see **Table S5.1**).

*Contemporary evolution population whole genome resequencing.* Candidate individual samples from the Rabbit Slough 2009 population for whole genome re-sequencing were chosen based on high estimated DNA concentrations. These were then examined on an electrophoresis gel and the 20 samples with the least evidence of DNA fragmentation were sent for high-coverage whole genome resequencing at Beijing Genomics Institute (BGI) using their proprietary DNBseq technology. Samples were sequenced to a target coverage of 30x based on 100bp paired end sequencing.

#### 8. ARRAY GENOTYPING

*SNP discovery and genotyping array design.* An Illumina GoldenGate custom SNP genotyping array was designed to genotype large numbers of individuals at SNPs tagging previously identified adaptive loci that have diverged in parallel among marine and freshwater fish throughout the stickleback species range (*13*). SNPs were selected to tag 72 "adaptive" genomic regions (235 SNPs) and 50 "random" and putatively neutral genomic regions (128 SNPs) defined by cyclic-rotation of the bed intervals of the tagged adaptive regions by a random integer to ensure roughly equal linkage-disequilibrium among adaptive loci and random loci.

Tagged SNPs were ascertained for the array using variant call data from Illumina short read sequencing of 10 marine and 11 freshwater fish (*13*). For targeted adaptive loci, between 3 to 5 SNPs with an allele frequency difference >=0.9 and with alternate alleles present in 4 or more individuals of each ecotype were selected for each target region. For "random" loci between 2-5 SNPs with a minor allele frequency between 0.35 to 0.65 in the combined sample of 21 fish were selected. Where possible SNPs with nearby variants in the flanking 60 base pairs were excluded, and the SNPs with the highest 'design score' from Illumina's design verification process were selected for the genotyping array.

*SNP genotyping.* SNP genotyping was performed on 250ng of genomic DNA using Illumina GoldenGate 384 custom SNP arrays (Illumina) according to manufacturer's protocols. Genotype calls were made using GenomeStudio 2.0 software (Illumina), with results from each SNP individually inspected to verify genotype clusters.

#### 9. REFERENCE GENOME CONSTRUCTION

*gasAcu1-4 creation.* We use as our reference genome a slight modification of the recent Hi-C guided improvement of the stickleback genome (*14*), which we term *gasAcu1-4*. The Peichel et al. 2017 Hi-C genome was not modified in any way except to address two long-standing issues that have persisted throughout many versions of the stickleback genome.

First, the subtelomeric region of chrVII is of great biological interest due to its well-documented role in controlling pelvic spine development (*15*). However, due to its highly repetitive nature, it is extremely difficult to assemble and many important sequences, including the key gene *Pitx1*, are missing entirely from existing genome assemblies, while other sequences in the region are scattered in small unassembled scaffolds. We address this issue by including the sequence from Salmon River BAC clones (Genbank GU130435) (*16*) as chrP and removing overlapping fragmented sequences from chrUn and the end of chrVII. We note that chrP is derived from a marine population, while the rest of the genome is from a freshwater population (Bear Paw Lake), so all analyses concerning this chromosome must be interpreted as such.

Second, the mitochondrial genome was previously split into two fragments buried within chrUn, while a separate mitochondrial genome sequence from Northern Japan was added as chrM. We corrected these issues by removing the duplicated Bear Paw Lake mitochondrial genome from chrUn and using it to replace the exogenous chrM sequence, resulting in a single copy of the mitochondrial genome derived entirely from Bear Paw Lake.

*Liftover chains.* LiftOver (*1*) chains (*17*) between gasAcu1 and gasAcu1-4 were generated from BLAT (*18*) alignments using the script https://github.com/ENCODE-DCC/kentUtils/blob/master/src/hg/utils/automation/doSameSpeciesLiftOver.pl as well as Kent Utilities (https://github.com/ucscGenomeBrowser/kent) following the procedure described here: http://genomewiki.ucsc.edu/index.php/DoSameSpeciesLiftOver.pl

*Annotation.* Ensembl (*19*) release 94 gene annotations for *gasAcu1* were downloaded via the UCSC Genome Browser and lifted to *gasAcu1-4* using all default liftOver parameters.

*Major Quantitative Trait Loci.* In order to annotate our new genome build with quantitative trait loci (QTL), we began with all QTLs from the comprehensive 2017 review of all reported stickleback QTLs by Marques and Peichel (*20*). We first filtered out results from stickleback species other than *Gasterosteus aculeatus,* and then for each reported interval lifted the coordinates from *gasAcu1* to *gasAcu1-4*. Finally, to create a set of well-bounded major QTL, we required each QTL to have PVE > 20 and have a reported confidence interval < 5Mb. All markers on chrP were assigned to the end of chrVII, where chrP physically resides. No filtering was performed on account of trait similarity within a cross or multiple crosses analyzing the same trait. This yielded 108 major QTLs. Some loci contain more overlapping QTL reports than visualized in Fig. 1F.

#### 10. BIOINFORMATIC PROCESSING & VARIANT CALLING

*Geographic population survey data.* Reads were mapped to *gasAcu1-4* with `bwamem` (0.7.17-r1188). `Picard` (v2.18) was used to add read groups and mark duplicate reads. Indel realignment, base quality recalibration was performed using `GATK` (v4.1.2.0), following their Best Practices guidelines. A subset of 50 genomes was used to bootstrap confident variant calls that were fed back into GATK as the truth set for base and variant recalibration. `GATK` (v4.1.2.0) (*21*) `HaplotypeCaller` was used to construct individual gVCFs and `GenotypeGVCFs` was used to perform multisample calling across the 227 samples. Variant recalibration was performed with `GATK` (v4.1.2.0) `VariantRecalibrator` and `ApplyVQSR`, and SNP and indel variant calls were accepted at -ts-filter-level 99.9. This yielded calls at 11,964,532 SNPs and 2,925,455 indels.

*Contemporary evolution population data.* Raw reads were trimmed and merged using `AdapterRemoval` (v2.2.2) (*22*). `bwa mem` (v0.7.15-r1140) (*23*) was used to map both paired and collapsed reads against *gasAcu1-4*. `Picard` (v1.93(1476)) was used to add read groups and mark duplicate reads. Indel realignment, base quality recalibration was performed using `GATK` (v3.7) (*21*). SNPs and Indels identified from the established population whole genome sequencing above were masked during BQSR and used as a reference set for indel realignment respectively.

*Contemporary evolution genome variant calling.* `GATK` (v3.7) (*21*) `HaplotypeCaller` was used to construct individual gvcfs and `GenotypeGVCFs` was used to perform multisample calling across the 20 Rabbit Slough whole genomes samples. Variant Quality Score Recalibration (VQSR) was also performed using `GATK` (v3.7) (*21*), with SNPs and Indels identified from the established population whole genome sequencing used as the truth set.

*Contemporary evolution Pool-Seq variant calling*. Allele frequencies in Pool-Seq populations at SNPs identified from the established population genomes was performed using the maximum-likelihood method of Lynch et al. (*24*) using custom scripts (https://github.com/kveeramah/Lynch-PoolSeq-estimator). We also calculated allele frequencies from a set of SNPs ascertained only in the Pool-Seq samples We first used `popoolation2` (v1201) (*25*) to generate a sync file describing allele-specific and indel read depths at each genomic position for each contemporary Pool-Seq population. We then used the method of Lynch et al. (*24*) to estimate allele frequencies for each position and extract any position with there was evidence in any population of a minor allele frequency greater than 5% and where no more than 1% of reads showed evidence of an overlapping indel.

#### 11. ECOTYPE GROUPING OF GEOGRAPHIC POPULATIONS

The sequenced established population genomes were divided into five subsets reflecting different geographic regions: those from the northeast Pacific (the primary analysis group discussed in this paper, “c150”, 57 freshwater and 12 marine), Northern Europe (“c151”, 18 freshwater and 9 marine), California freshwater with all Pacific marine (“c153”, 18 freshwater and 15 marine), a balanced global selection (used in Fig. 1D, “c155”, 56 freshwater and 28 marine), and a subset thereof containing only populations from regions glaciated by the last Ice Age (“c154”, 38 freshwater and 25 marine). While multiple individuals were sequenced from some populations, a maximum of one randomly selected fish per population is used in these analysis groups to prevent undue overweighting. The global analysis group contains roughly equal numbers of populations from the northeast Pacific, California, and Europe, as well as a few others around the world. For full details on samples included in each analysis group, see **Table S11.1**.

#### 12. ECOPEAK IDENTIFICATION

Two complementary methods were used to identify EcoPeaks in the older, established populations: a genetic distance-based approach over small windows, and a test of random allelic distribution at each individual variant base.

First, we utilized the same genetic distance-based approach as Jones et al. (*13*). This involves building N×N pairwise nucleotide divergence ($\pi$) matrices for small overlapping windows tiled across the genome (2500 bp windows with 500bp step size), where N is the total number of samples in each analysis group. The distance matrix was then used to calculate a marine-freshwater cluster separation score (CSS), quantifying the average marine-freshwater genetic distance after subtracting the genetic distance found within each ecotype. All details are the same as in Jones et al. (*13*) except for the following increases in stringency: at least 4 variable sites were required in the window; at least 2/3 of samples had to have a call at any particular base for the base to be used; and the reference distribution to estimate p-value was generated through Monte Carlo sampling. Due to the larger number of samples in this study, an exhaustive examination of all possible combinations was infeasible, and so we continued Monte Carlo sampling until either 10 samplings as extreme as the actual data were observed, or 1,000,000 combinations were generated (compared to 352,716 combinations in Jones et al. (*13*)). If after one million samplings none were as extreme as the observed data, the window was conservatively treated as 0.5 successes/1,000,000 draws and given a p-value of 5e-7 (**Fig S12.1**). As many windows were this extreme, we also computed a separate z-score for each window by fitting the observed CSS value to a normal distribution (**Fig S12.2**).

Second, we analyzed the distribution of allele counts between marine and freshwater populations at every base in the genome with two alleles present at >10% frequency in the combined analysis metapopulation. At each such base, after conditioning on the observed number of homozygous reference, heterozygous, and homozygous non-reference calls, we used a multivariate hypergeometric generalization of Fisher's Exact Test to compute a two-sided p-value for the probability of an imbalance in allele counts at least as extreme as observed occurring by chance (**Fig S12.3**)

For both the SNP-based and 2,500 bp window-based analysis, nearby significant values were grouped into peaks that appear to behave as a single unit. On each chromosome, starting from the most significant unassigned p-value, points were greedily added to the peak window if above significance s and within distance d of another point within the peak. For the sensitive call set, s was set to two LOD below the significance threshold, while for the specific call set s was set to halfway between two LOD below threshold and the maximum peak value (and a minimum of 2 LOD below maximum peak value). Changes in d result in only minor differences in peak boundaries; for all call sets d was set to 50,000 bp. Peak extension into a more significant peak results in disqualification of the weaker peak, as it is not clearly independent of the primary peak.

Peaks were filtered at either a 1% false discovery rate for the specific calls or at 5% for the sensitive calls. The single base and window peaks were then intersected for the final specific calls, or unioned for the final sensitive calls. These final sets of peaks were subsequently termed EcoPeaks.

In addition to the northeastern Pacific basin and global analyses described in the main text, this analysis was also performed on additional geographic subsets of the sequencing data, as described above in Section 11, to generate additional EcoPeaks call sets for populations from Northern Europe, a subset of global regions covered by glaciers in the last ice age, and freshwater California in addition to the northeastern Pacific basin and global analyses described in the main text (**Fig S12.4**).

We also assessed the relative contribution of coding and non-coding mutations. Following Jones et al. 2012 (*13*), EcoPeaks and TempoPeaks were classified as "regulatory" if they did not overlap any known genes, "coding" if they contain at least one mutation consistently differentiated between ecotypes ($p < 10^{-6}$) that alters an amino acid, and otherwise as "probably regulatory". This approach is conservative for calling regulatory regions and overcounts coding regions, especially for larger EcoPeaks and TempoPeaks, as even single amino acid changes that may have inconsequential functional effects will result in classifying the entire region as "coding". Consistent with the previous study, we classified a large majority of regions as either "regulatory" or "probably regulatory" in all call sets (**Fig S12.5**), and attribute the slight differences in proportion of "coding" regions between call sets primarily to region size.

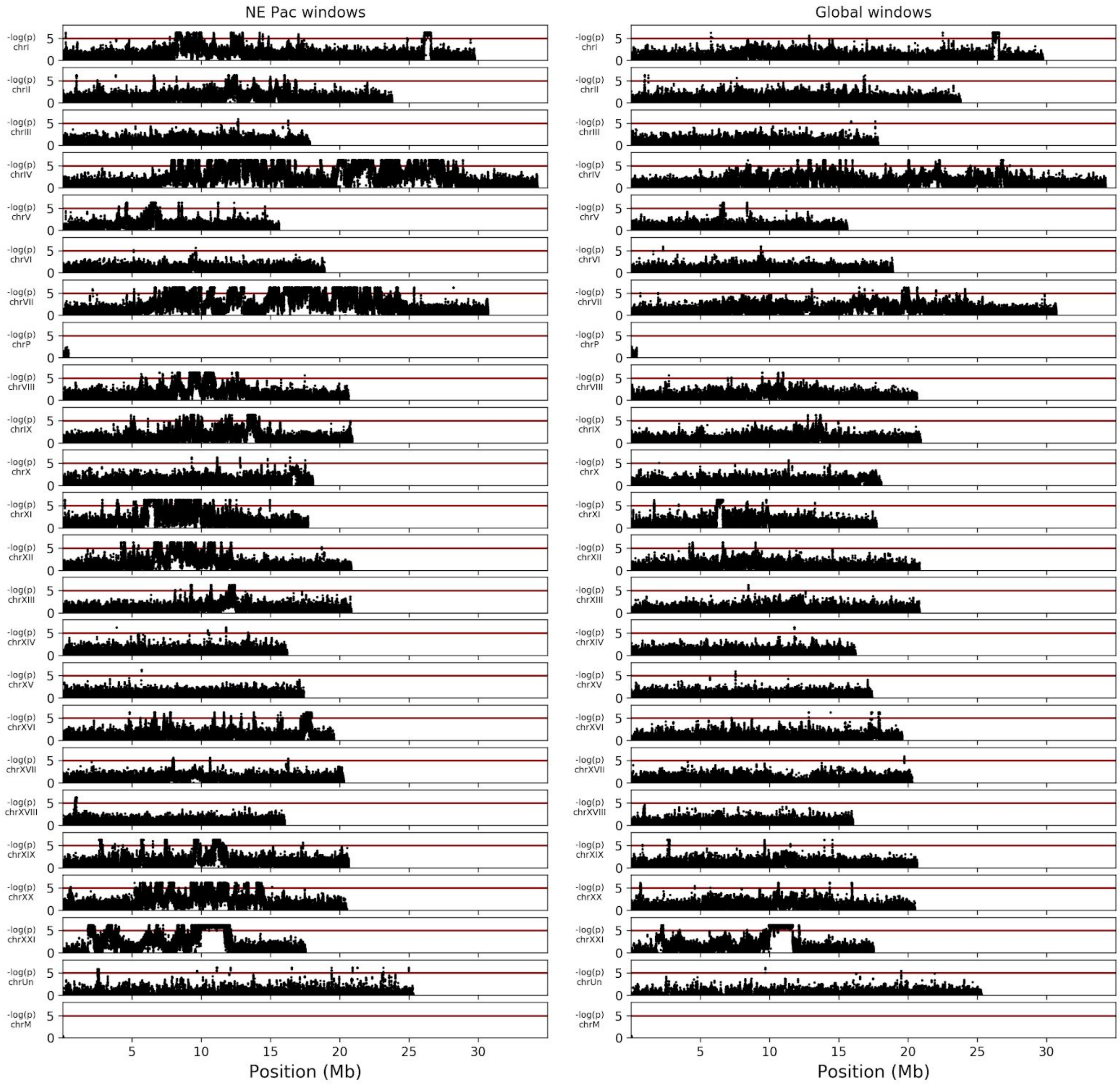


**Fig_S12.1:** Detailed view of CSS window-based p-values in either the northeast Pacific or globally. The red line represents an unadjusted p-value of 1e-5 (<10 windows expected beyond this threshold by chance genome-wide).

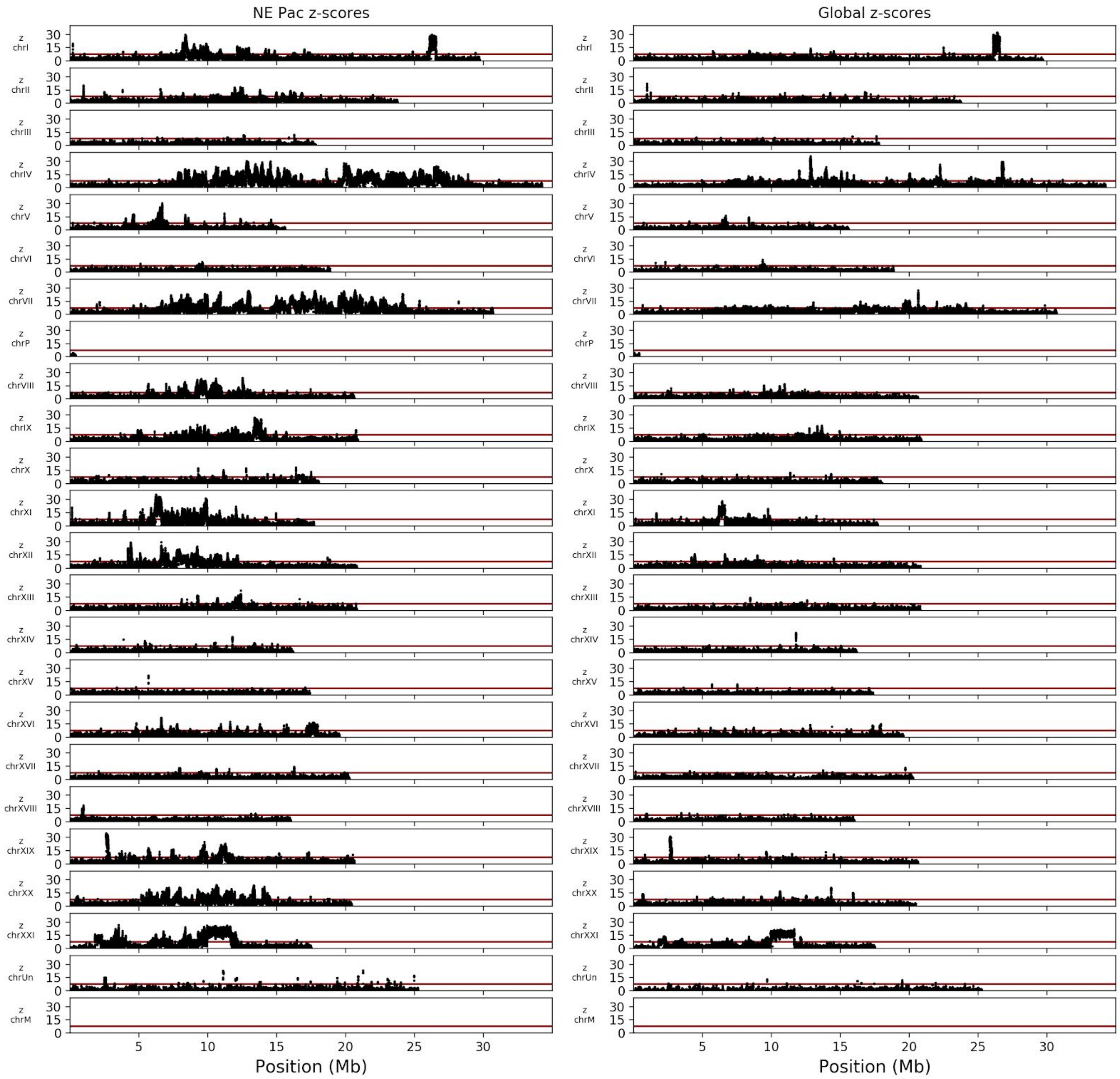


**Fig_S12.2:** Detailed view of CSS window-based z-scores in either the northeast Pacific or globally. The red line represents a z-score of 7.5.

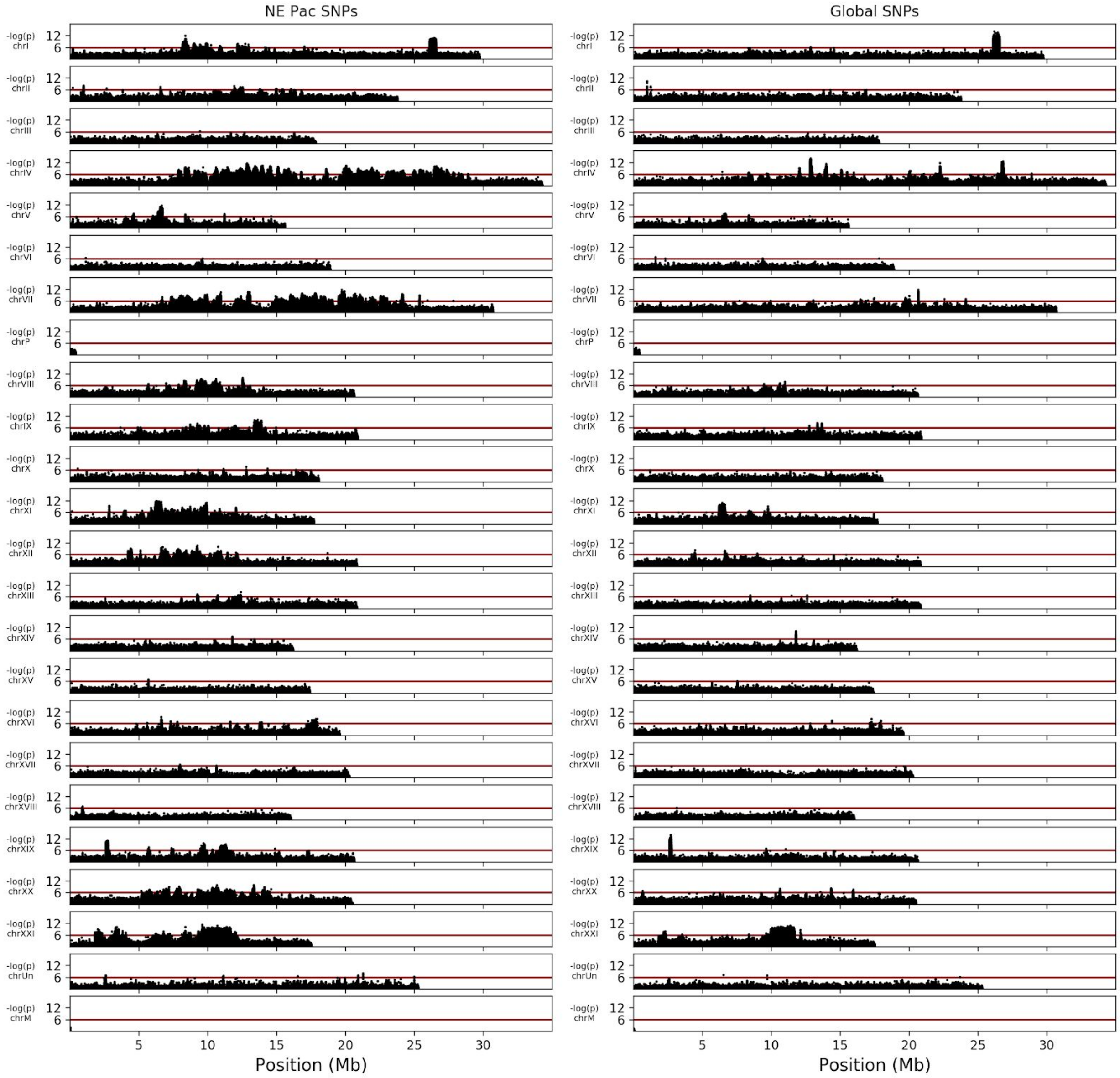


**Fig_S12.3:** Detailed view of SNP-based analysis in either the northeast Pacific or globally. The red line represents an unadjusted p-value of 1e-6 (5 SNPs expected beyond this threshold by chance genome-wide).

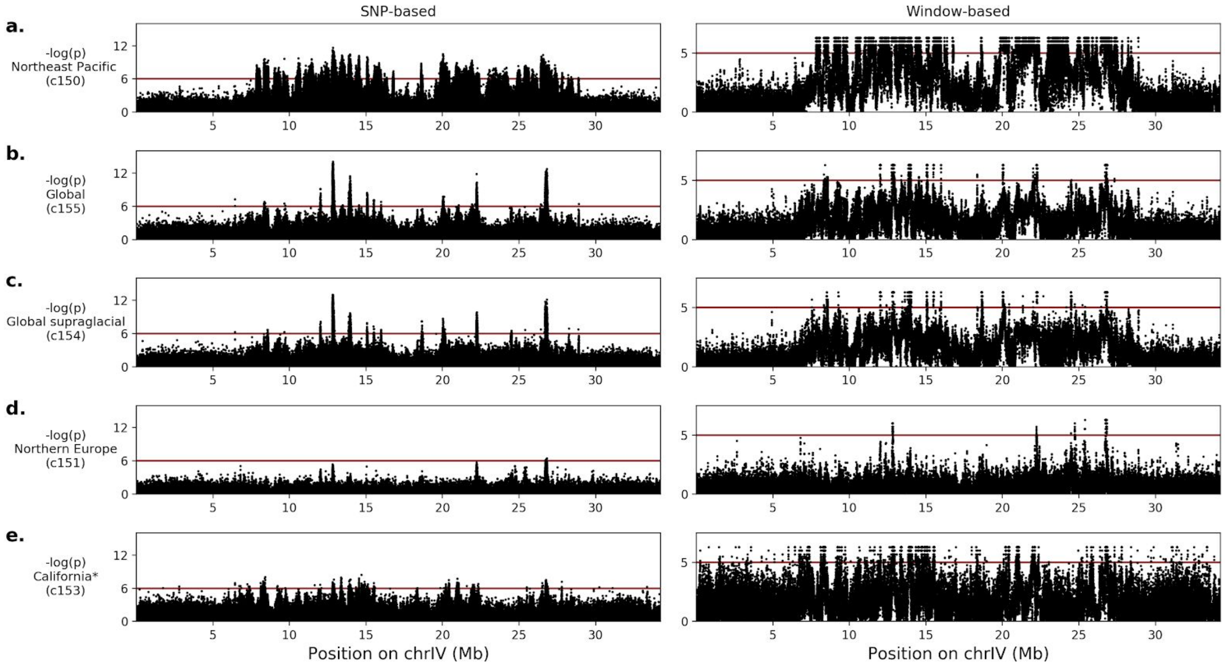


**Fig_S12.4:** Detailed comparison of results for chrIV in five geographic analysis groups. The California freshwater fish are compared to all Pacific marine fish. Sample numbers and statistical power vary between groups. Genome-wide, we expect <10 results in any analysis group beyond the significance indicated by the red line by chance.

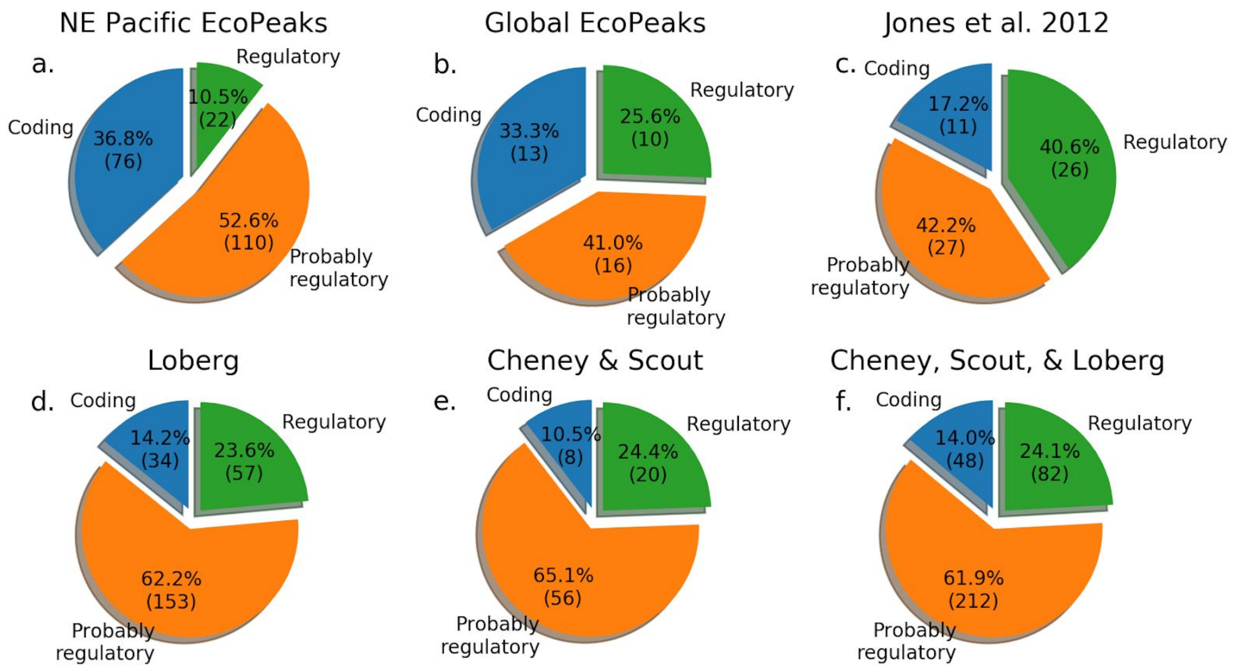


**Figure S12.5: Classification of EcoPeaks and TempoPeaks into coding, regulatory, and probably regulatory.** Peaks are classified as regulatory if they do not overlap a coding sequence, coding if they contain a consistently differentiated amino acid change, and probably regulatory if they overlap a coding sequence but do not contain a consistently differentiated amino acid change. This thus represents an upper bound on the proportion of coding changes. The results in c. are as reported in Jones et al. 2012 for the strictest set of region calls.

#### 13. HYBRID ZONE ANALYSIS

To confirm that increased EcoPeak differentiation in the Pacific was not due to increased freshwater genetic homogeneity within the northeast Pacific, we also analyzed hybrid river systems from the Pacific and Atlantic basins with gene flow between the upstream freshwater and downstream marine populations for two signatures of selection, high Fst and low heterozygosity. We see far fewer genomic windows containing either high Fst or low heterozygosity within each hybrid zone in the Atlantic than in the Pacific, suggesting there actually are far more genomic regions under selection in the Pacific (**Fig S13.1**). This is consistent with other reports and supports the northeast Pacific as ideal for studying the optimal properties of evolution (*13*, *26*).

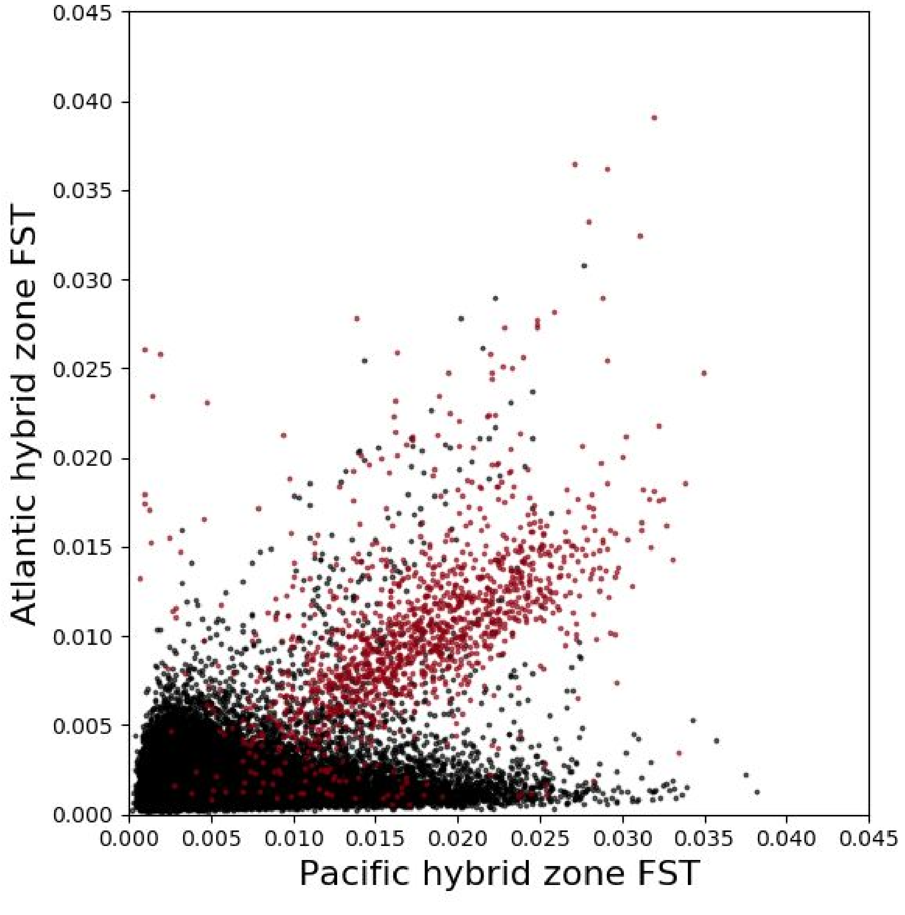


**Fig_S13.1:** Comparison of genomic window FST values in Atlantic and northeast Pacific river systems. Each dot represents a 2500 bp genomic window, with windows overlapping specific global EcoPeaks in red, with FST calculated between upstream freshwater and downstream marine stickleback. There are substantially fewer windows of elevated FST in the Atlantic population (Midfjardara River, Iceland) than in the Pacific population (Little Campbell River, British Columbia).

### 14. ALLELIC AGE ESTIMATION

Allelic divergence and age was computed from five upstream (freshwater) and five downstream (marine) fish from Little Campbell River. In non-overlapping 1kb windows were tiled across the genome, variants homozygous for different alleles were counted and used to compute marine-freshwater sequence divergence *d*, with a minimum set to 0.5 variants/window. Allelic age was then estimated from sequence divergence by three methods: a uniform rate of 0.00571 changes/base/My, the mean synonymous substitution rate to *Gasterosteus nipponicus* and *Gasterosteus wheatlandi (27)*; a variable rate based on the number of substitutions in the variants in *Gasterosteus nipponicus* data aligned to the *Gasterosteus aculeatus* genome *(28)*, calibrated to species divergence 2 Mya (*29*); and a variable rate based on the sequence identity with the *Pungitius pungitius* reference genome calibrated to species divergence 26 Mya (*30*). Windows without clear alignment were excluded. The median of the remaining estimators was used for all analysis.

The fold-change between typical genomic regions and the oldest regions is comparable to a recent report (*31*), but our estimates of absolute age are closer to carefully validated Sanger sequencing-based analysis of third base pair substitution rates (*27*). We also examined the relationship between estimated allele age and recombination rate (**Fig S14.1**). We observed a non-significant trend for older alleles to have lower recombination rates. If the trend is real, it may reflect older alleles having more time to suppress recombination and form "supergenes" (*32*, *33*). Older EcoPeaks are larger whether measured in physical distance or genetic distance.

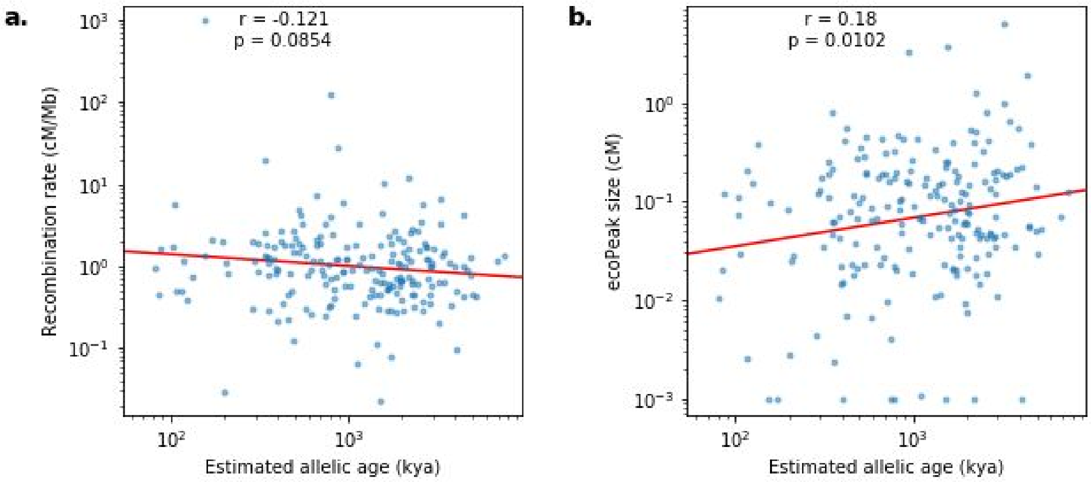


**Fig_S14.1:** Recombination rates and ecoPeak sizes compared with estimated allelic ages. a. Visualization of northeast Pacific EcoPeak age vs. recombination rate, with weak association. b. Visualization of northeast Pacific EcoPeak age vs genetic size, with the same trend of older regions being larger by genetic size seen with physical size.

### 15. FREQUENCY AND HAPLOTYPE STRUCTURE FROM ARRAY GENOTYPES

To characterise the frequency, extent of haplotype tracts, and linkage disequilibrium of freshwater-adaptive standing genetic variation among marine sticklebacks, we genotyped 751 Rabbit Slough (RABS), 655 Resurrection Bay (RSBY) and 237 Glacier Spit (GLSP) wild caught marine sticklebacks using a custom Illumina GoldenGate array.

*Data filtering.* The genotyping data was filtered to remove individuals with excessive missing genotype calls (mind 0.3), and SNPs with excessive failure rate or heterozygosity using Plink (*34*) (filter parameters `-mind 0.33`; `-geno 0.33`; `-het 0.8`) to retain a dataset comprising 302 snps and 742, 628, 199 individuals for RABS, RSBY and GLSP respectively.

*Estimating Frequency of Freshwater-adaptive alleles.* Diploid genotypes were recoded to represent the number of derived freshwater alleles [0,1 or 2] where the derived allele was defined to be the minor allele across all marine fish. To estimate the frequency of freshwater adaptive alleles in the marine populations we first extracted SNPs falling within 'sensitive' EcoPeaks and/or adaptive loci identified by Jones et al (2012). To ensure the distribution of allele frequencies was not biased towards genomic regions containing several closely linked adaptive loci or by adaptive regions tagged by multiple array SNPs, we merged the bed intervals of adaptive loci within 50kb of each other and estimated allele frequency using a single SNP from each merged region (44 adaptive regions total).

The frequency of freshwater adaptive alleles ranged from 0-0.07 (RABS), 0-0.09 (GLSP) and 0-0.14 (RSBY), with median values less than 0.01 for all RSBY and GLSP and less than 0.001 for RABS (**Fig S15.1**). Rabbit Slough fish are less likely to carry freshwater adaptive alleles than fish from Glacier Split and Resurrection Bay suggesting they are less affected by introgression with local freshwater populations. We note that the frequency of alleles at each locus is highly correlated among populations with the exception of two loci at comparatively high frequency in RSBY and GLSP, but low frequency in RABS (the chrI inversion at chrI:26.5Mb, and chrV:8.35Mb). This hints at some fine grained population substructure in marine populations - Rabbit Slough is located in the Knik Arm of the Cook Inlet, and is more than 340km away as the fish swims from Glacier Spit and Resurrection Bay (both of which are more proximal to the North Pacific Ocean).

*Identifying Carriers of Freshwater Alleles*
For each individual, we summed the number of derived alleles at SNPs tagging adaptive loci and classified fish carrying one or more alleles to be putative "carriers" of freshwater-adaptive standing genetic variation. Direct migration of freshwater fish or F1 hybrids through the marine environment to neighbouring freshwater populations has been hypothesized to be an efficient mechanism for the spread of freshwater adaptive alleles (*35*), assuming the migrants could survive strong negative selection pressures while in the marine environment. In this survey of

1569 marine sticklebacks, we did not find any evidence of F1 hybrids or direct freshwater migrants. In contrast, we observed considerable diversity in the standing genetic variation carried among individuals within and between marine populations (**Fig S15.2, Fig S15.4, Fig S15.6**). As expected from the low allele frequency within each population freshwater-adaptive alleles were most commonly carried in heterozygous form at typically one locus. However, rare individuals carrying freshwater adaptive haplotypes at multiple loci across the genome were also observed (range 1-8 chromosomes of 13 studied). For example, two individuals in RSBY carry homozygous tracts of freshwater-adaptive alleles at multiple, but not all loci targeted by the genotyping array (**Fig S15.4**). Notably these individuals are a mosaic of freshwater homozygous, heterozygous and marine homozygous tracts across their genome indicating they arise from recent admixture and backcrossing among marine and freshwater ecotypes. This highlights the importance of hybridization and admixture as a source of standing genetic variation that can be used as a substrate for rapid adaptation in the future.

*Extent of Freshwater Haplotype Blocks Carried by Marine Fish*

The size of freshwater haplotype blocks carried by marine individuals and the potential for the blocks to include freshwater adaptive variation at multiple loci across a chromosome) has direct implications for the rate of rapid adaptation in subsequent freshwater colonisations. Since the size of freshwater haplotype blocks will decay at a rate proportional to 1-recombination rate, the extent of the haplotype block along a chromosome, to the degree that it can be determined from the SNPs tagged by this array, is an indicator of the amount of time the freshwater adaptive haplotype has been in the marine environment. F1 hybrids are expected to be heterozygous at adaptive loci across the entire length of each chromosome, while subsequent generations will carry smaller tracts of heterozygosity. The SNP genotyping array tagged multiple adaptive loci on chromosome IV spanning 14Mb enabling us to explore the extent of freshwater haplotype blocks carried by marine fish.

Of 1484 chromosomes IV sampled from RABS marine fish, only one fish showed a contiguous run of heterozygosity that spanned SNPs in a small 20kb region at the EDA locus chrIV:12821275-12841217 (**Fig S15.2**). This individual is homozygous for marine alleles at other adaptive loci on chrIV indicating the freshwater haplotype block observed has been present in the marine population for many generations. The paucity of any other marine fish carrying extensive freshwater adaptive haplotype tracts on chrIV in RABS contrasts with our samples from RSBY and GLSP, and may indicate lower geographic proximity to marine-freshwater hybrid zones and/or stronger selective purging of freshwater alleles in RABS marine fish. In RSBY, we observed extensive haplotype blocks spanning at least 4 distinct adaptive loci tagged on the array over as much as 11.8Mb in four of 1256 chromosomes IV (0.0032%; **Fig S15.4**). Further, in GLSP carriers, a single individual out of 398 chromosomes sampled (0.0025%) was found to be heterozygous at 96% of chrIV SNPs tagged indicating it carries freshwater adaptive alleles at as many as six adaptive loci on this chromosome (Fig **S15.6**). Combined, these results indicate that large freshwater-adaptive haplotype blocks spanning multiple adaptive loci can be found in the marine population at very low frequency. As a source of partially assembled (linked) adaptive cassettes have the potential to dramatically increase the rate of adaptation in new freshwater habitats.

*Linkage Disequilibrium Among Adaptive Loci*

Since divergent adaptation to marine and freshwater environments is highly polygenic involving loci on almost all chromosomes, we explored the extent to which marine sticklebacks carry freshwater adaptive alleles on multiple chromosomes and looked for evidence of inter-chromosomal linkage disequilibrium (LD). The frequency dependency of some LD estimators (eg $r^2$) can make comparisons across loci challenging - especially for paucimorphisms (rare alleles) whose frequencies may differ by an order of magnitude across loci (*36*). In addition to $r^2$, we therefore also calculated D' (*37*) in which the two-loci linkage disequilibrium is standardized using the allele frequencies at the loci in question. Calculations were performed in `R` with unphased genotype data using the library `Pould` (*38*).

Previous studies reported evidence for considerable linkage disequilibrium among loci in Alaskan marine sticklebacks from Rabbit Slough and Resurrection Bay (*39*). However, the low frequency of freshwater adaptive alleles in these populations (<1%), makes the conclusions of the previous study questionable given the modest sample size used (18 individuals from Resurrection Bay, and 14 individuals from Rabbit Slough). Further, the findings are most likely false positives caused by artificial population substructure. The authors analysed fish from Rabbit Slough and Resurrection Bay together as a single population, assuming no geographic substructure exists among their samples, yet our analysis of allele frequencies at adaptive loci (above) show such structure very likely exists.

Here, with considerably larger samples sizes of 742, 628, 199 individuals for Rabbit Slough, Resurrection Bay and Glacier Spit respectively, we find evidence for only modest linkage disequilibrium among adaptive loci across the genome. Our analyses are still relatively underpowered given that most interchromosomal allelic associations are driven by very rare carriers (eg in Rabbit Slough, the heatmap in **Fig S15.3** suggest strong LD among chrXI and chrXIX ($r^2$), that is driven by a single carrier individual who is heterozygous for alleles on both chrXI and chrXIX (**Fig S15.2**). Further, the D' standardized measure of LD suffers from the problem of finding complete LD=1 when a rare allele is only ever sampled in association with one of the alleles at the second locus. Combined, we find little evidence for inter-chromosomal linkage disequilibrium among adaptive loci and stress the importance of very large sample sizes required (perhaps 10 fold larger than our current sample sizes) in order to explore this phenomena among paucimorphisms with sufficient power.

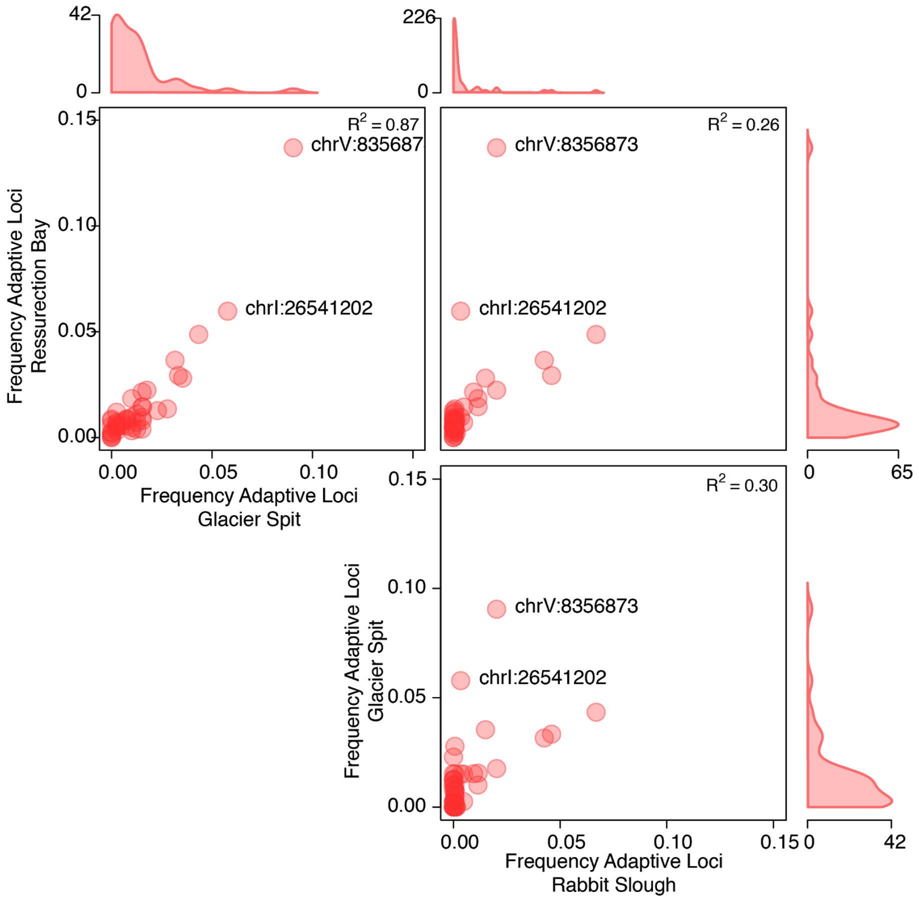


**Fig_S15.1 Frequency of freshwater derived alleles in three Alaskan marine populations.** Each point represents an adaptive locus with frequency estimated from a single SNP per locus tagging the divergent marine and freshwater haplotypes. Density plots show the distribution of frequencies at the 44 adaptive loci analyzed.

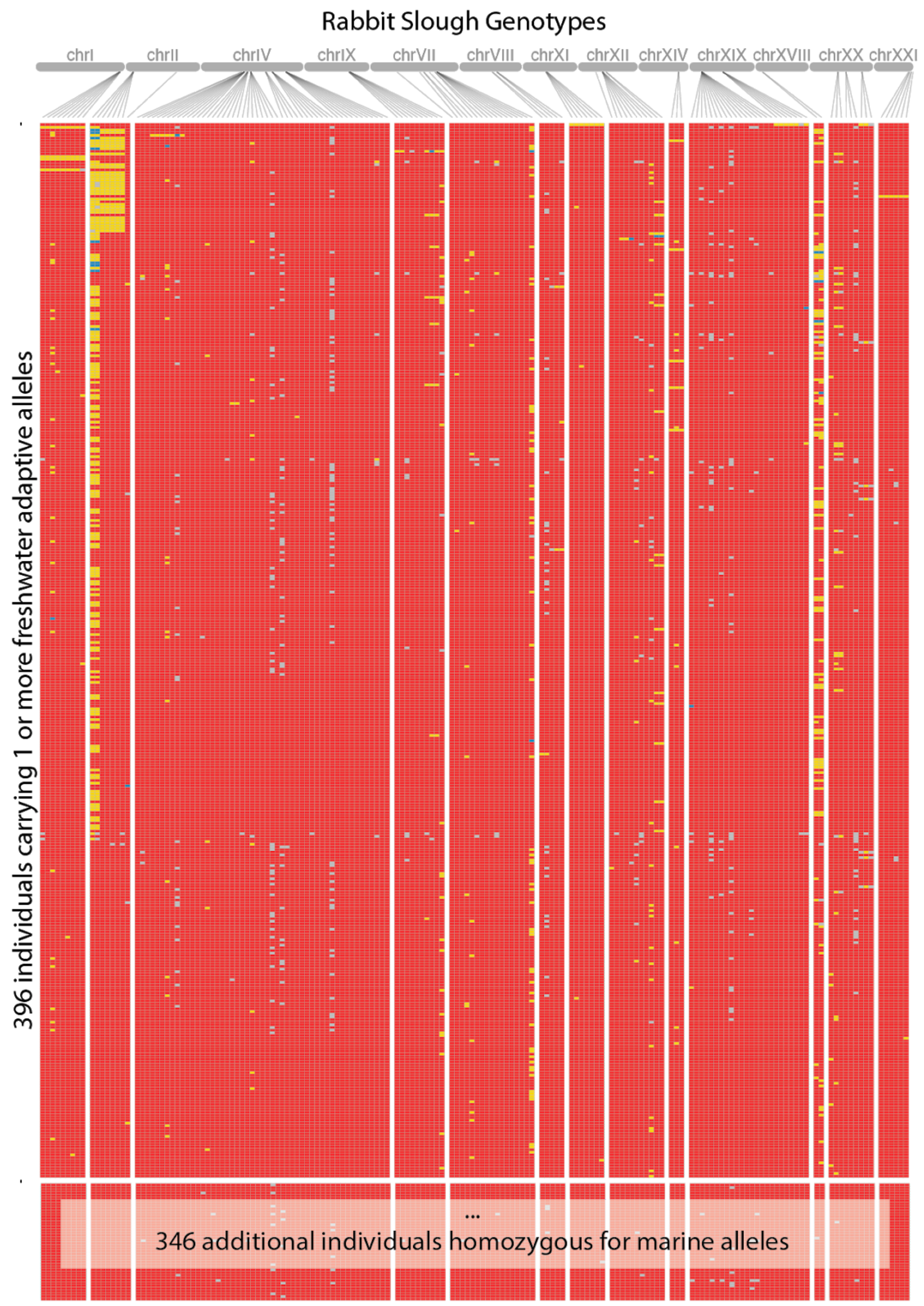


**Fig_S15.2** A visual genotype showing 742 Rabbit Slough marine fish at tagged adaptive loci across the genome. Rows represent individuals (sorted from top to bottom in descending order based on the total number of derived alleles each individual carries); columns represent SNPs. Red = homozygous for marine (major) allele; Blue = homozygous for freshwater (derived/minor) allele; Yellow = heterozygous; Grey = missing data;

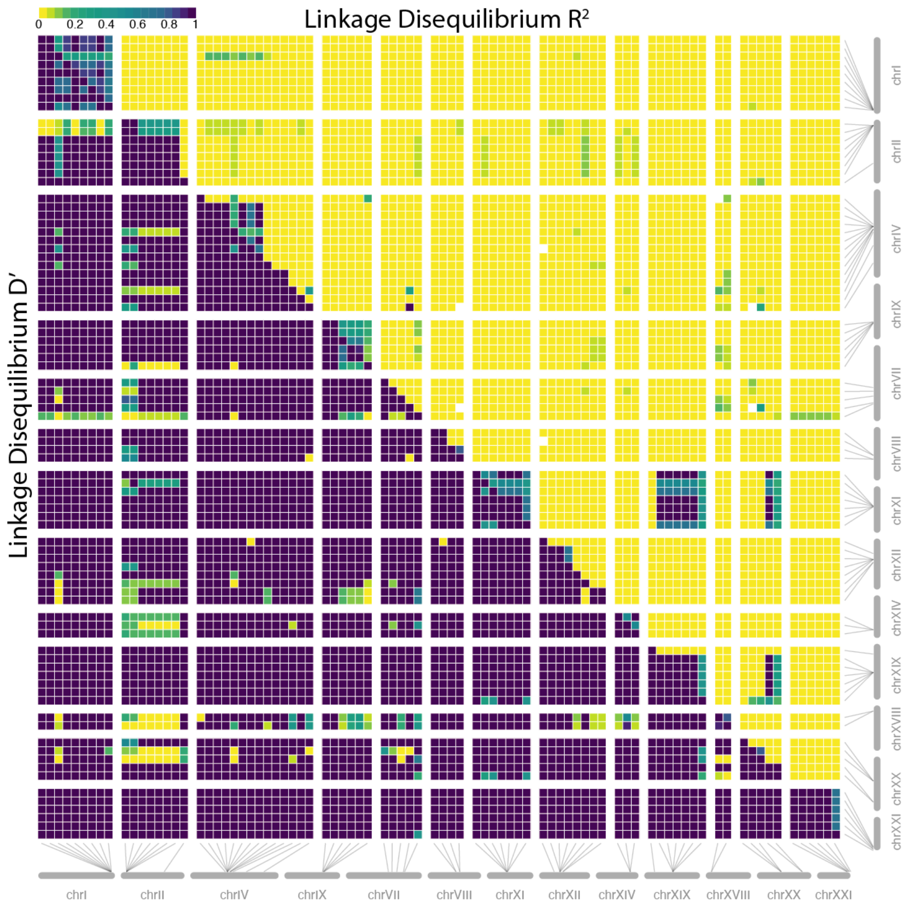


**Fig_S15.3** Estimates of linkage disequilibrium among adaptive alleles across the genome of Rabbit Slough fish. Upper triangle shows $r^2$, while lower triangle shows D'. SNPs that were fixed for one allele in the focal population were excluded from the analysis. Darker colors indicate strong LD.

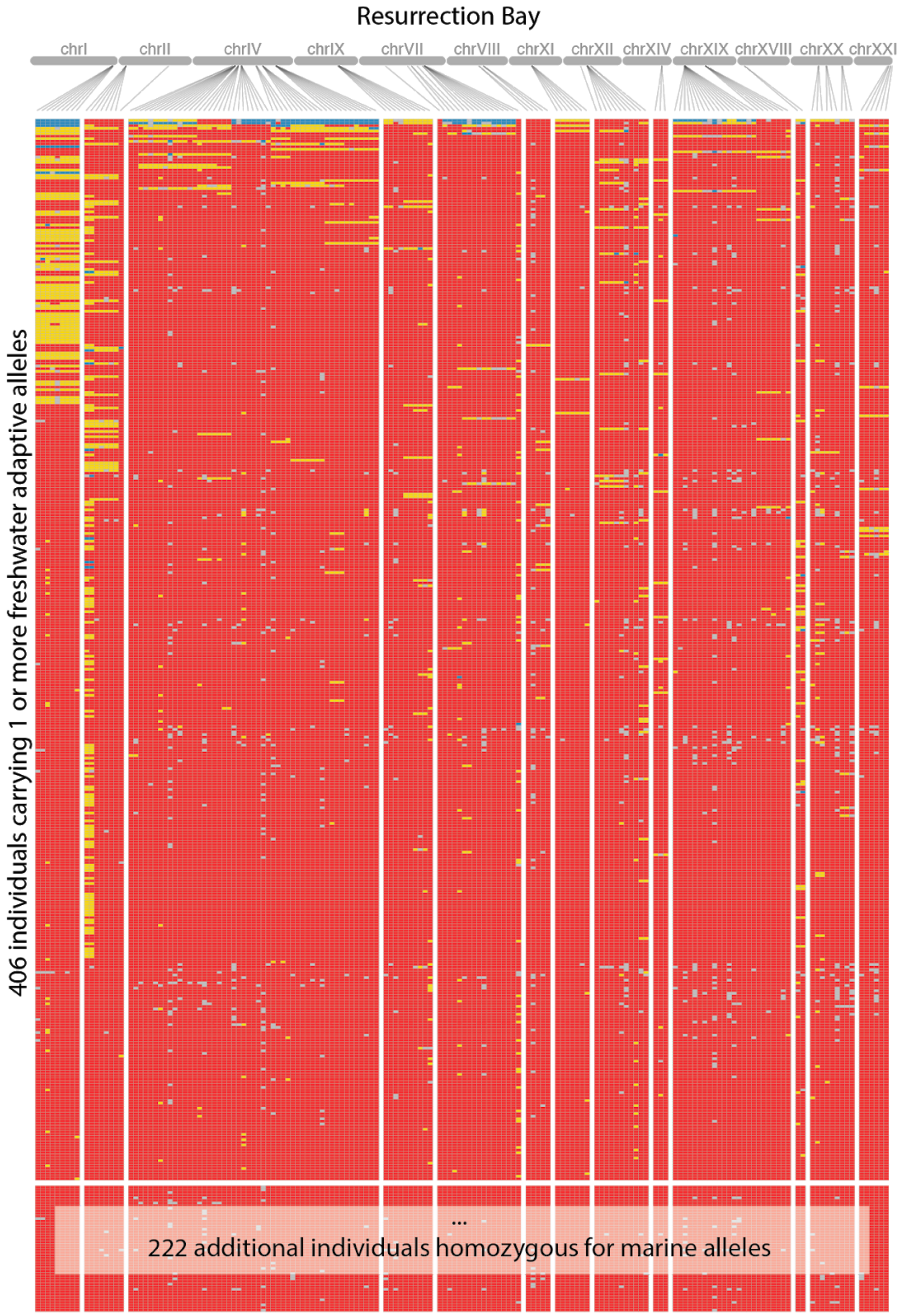


**Fig_S15.4** A visual genotype showing 628 Resurrection Bay marine fish at tagged adaptive loci across the genome. Rows represent individuals (sorted from top to bottom in descending order based on the total number of derived alleles each individual carries); columns represent SNPs. Red = homozygous for marine (major) allele; Blue = homozygous for freshwater (derived/minor) allele; Yellow = heterozygous; Grey = missing data;

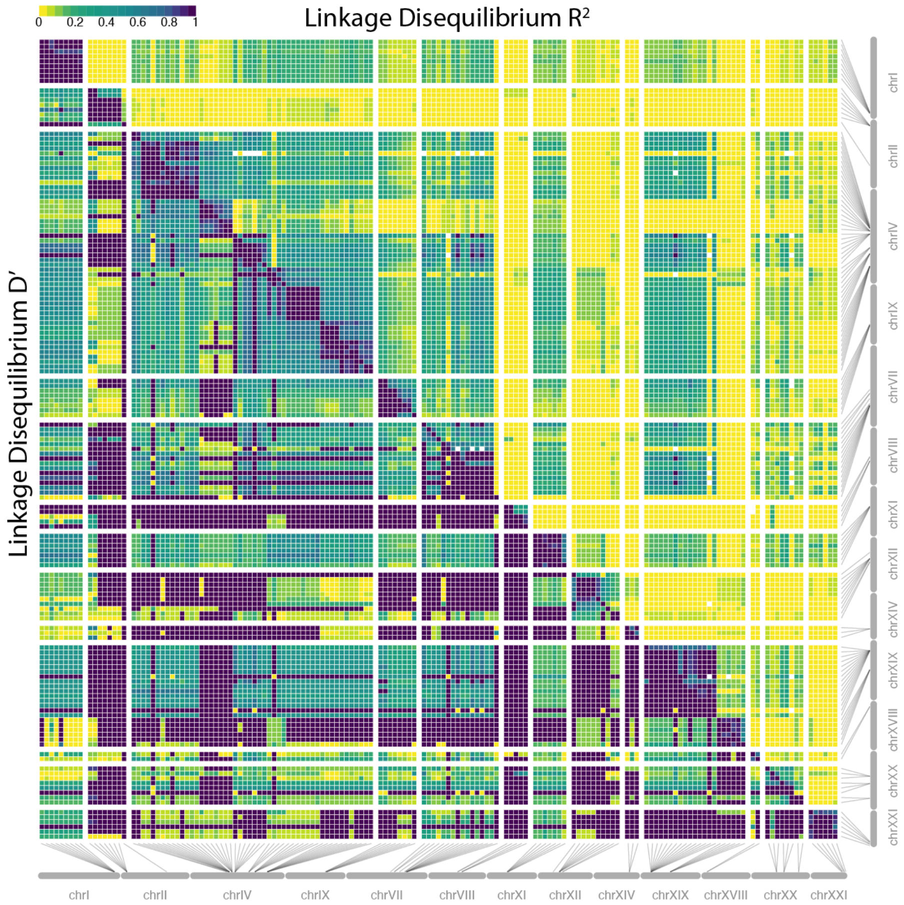


**Fig_S15.5** Estimates of linkage disequilibrium among adaptive alleles across the genome of Resurrection Bay fish. Upper triangle shows $r^2$, while lower triangle shows D'. SNPs that were fixed for one allele in the focal population were excluded from the analysis. Darker colors indicate strong LD.

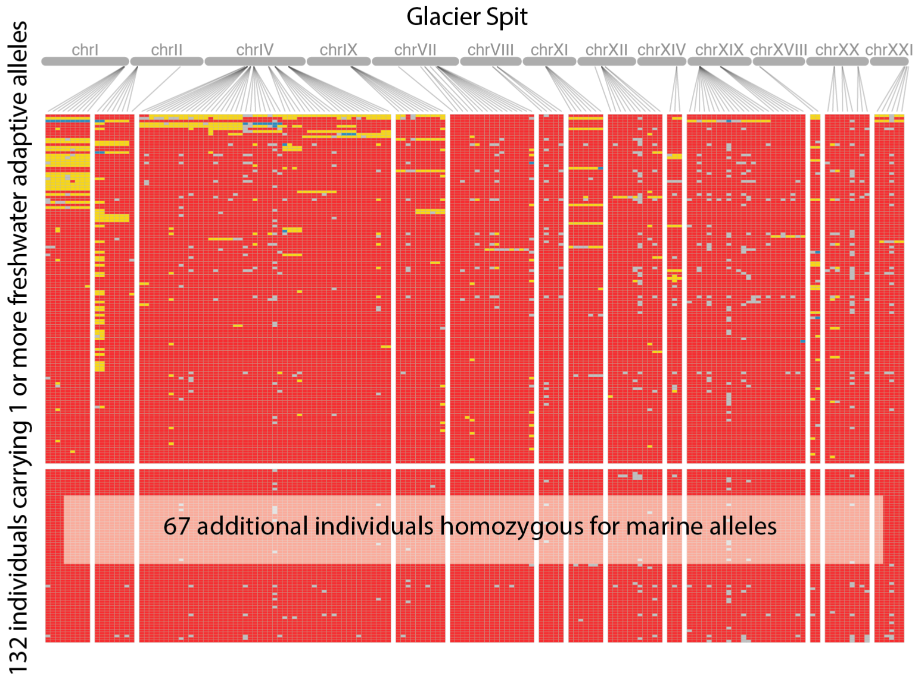


**Fig_S15.6** A visual genotype showing 742 Glacier Spit marine fish at tagged adaptive loci across the genome. Rows represent individuals (sorted from top to bottom in descending order based on the total number of derived alleles each individual carries); columns represent SNPs. Red = homozygous for marine (major) allele; Blue = homozygous for freshwater (derived/minor) allele; Yellow = heterozygous; Grey = missing data;

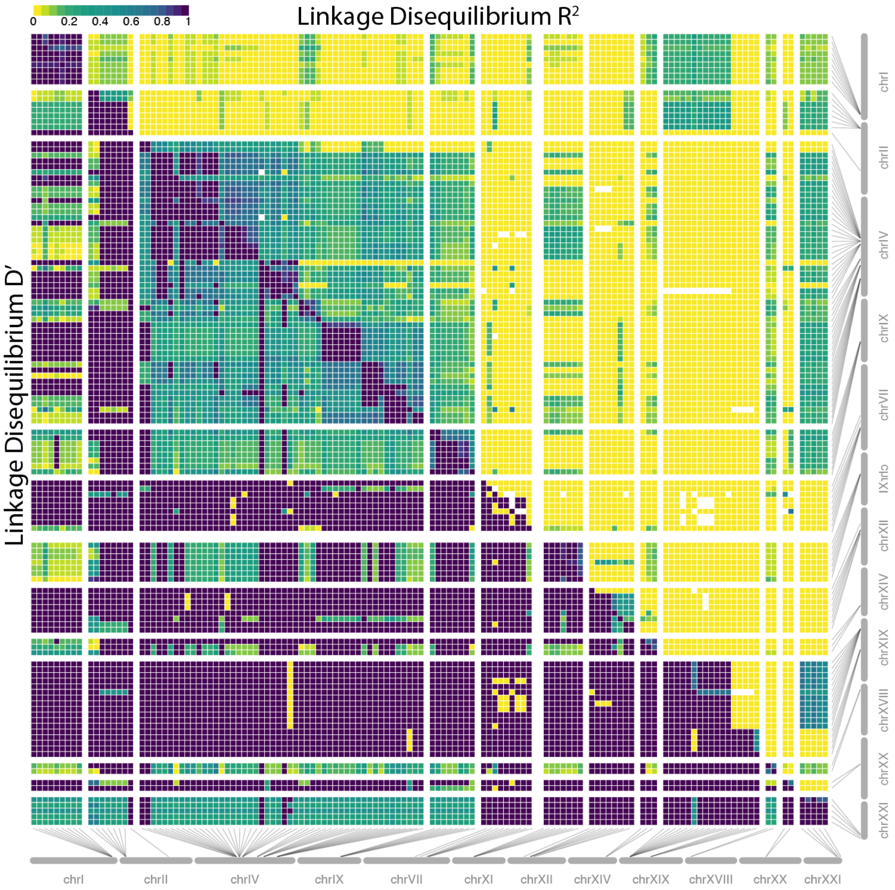


**Fig_S15.7** Estimates of linkage disequilibrium among adaptive alleles across the genome of Glacier Spit fish. Upper triangle shows $r^2$, while lower triangle shows D'. SNPs that were fixed for one allele in the focal population were excluded from the analysis. Darker colors indicate strong LD.

#### 16. COMPARING WGS, ARRAY AND POOL-SEQ ALLELE FREQUENCY ESTIMATES

In order to examine the reliability of our Pool-Seq allele frequency estimates we took advantage of having comparable estimates from Rabbit Slough via our whole genome sequencing and SNP-array data. When comparing our Pool-Seq estimates at ~4 million SNPs to the equivalent estimates based on 20 high coverage whole genomes, we found a highly significant positive correlation ($r = 0.975$, $p << 0.01$ ) with no noticeable bias in terms of under or overestimation (the best fit linear regression was essentially $y = x$, see Fig S16.1a). We then compared the Pool-Seq estimates to allele frequencies derived from 751 Rabbit Slough genotyped on the SNP-array. After filtering out A<>T and C<>G SNPs to avoid strand flipping effects in the array data and those with a missing rate of greater than 5%, we had 186 SNPs to compare between the two Rabbit Slough data sets. We again found a highly significant relationship when comparing Pool-Seq estimates and SNP array data ($r = 0.99$, $p << 0.01$) with no clear biases (**Fig S16.1**).

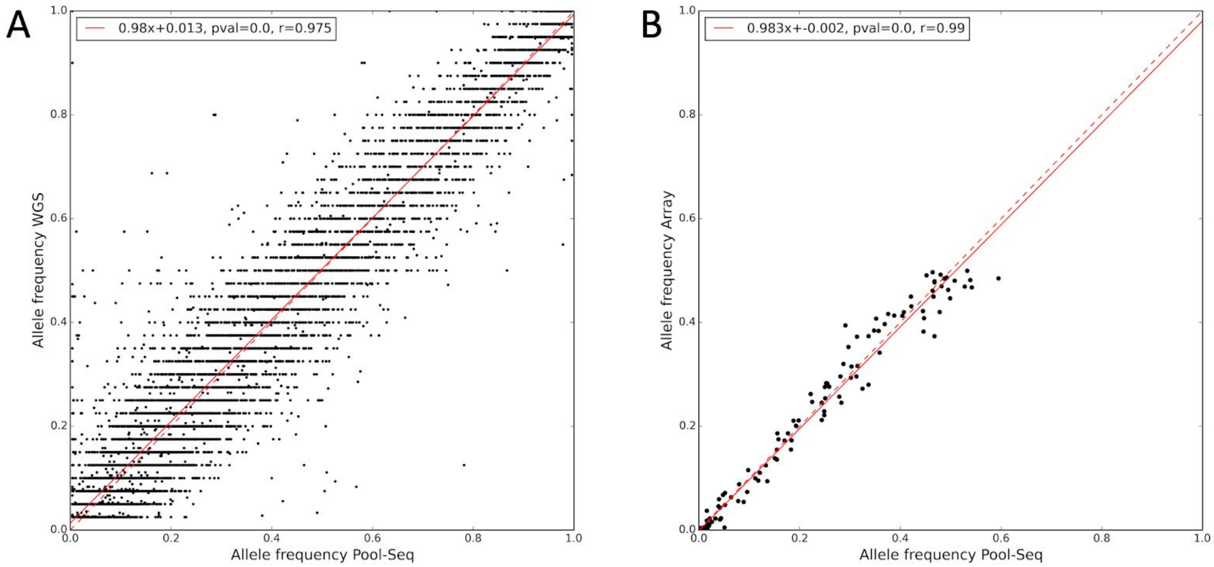


**Fig_S16.1:** Plot of Rabbit Slough SNP allele frequencies estimated from Pool-Seq versus a) 20 whole genomes and b) 751 SNP-array samples. Red dotted line shows expected correspondence, red solid line shows best fit linear regression model. In the case of a) only 10,000 randomly chosen SNPs are plotted, and allele frequencies refer to non-reference values, while in the case of b) they refer to minor allele frequency.

#### 17. RECOMBINATION MAP CONSTRUCTION

*Population genetic analysis of contemporary Rabbit Slough genomes*. We filtered down our Rabbit Slough variants to a set of high quality SNPs based on the VQSR "pass" tranche and each individual possessing coverage that was within half and double the genome-wide mean coverage (nb_snps = 3,577,312). We further filtered out SNPs with a minor allele frequency less than 5%, removed SNPs in high LD using `PLINK` (v1.90b3.38) (*40*, *41*) (`--indep-pairwise 50 5 0.2`), and excluding chrXIX (final number of snps = 548,955). Kinship analysis using the `--genome` function in PLINK showed no evidence of biological relatedness amongst specific samples (pairwise PI_HAT values ranged from 0.04 and 0.06, mean 0.05). Principal Component Analysis using `smartpca` (version: 10210) (*42*) showed no evidence of major population structure when visualized along the first four PCs (**Fig S17.1**), while the smallest cross-validation error during an model-based clustering analysis (`ADMIXTURE` (*43*) Version 1.3.0) was observed for K=1 (**Fig S17.2**).

*Recombination map construction.* Given the general population genetic homogeneity of our Rabbit Slough genomes, we used all 20 to construct a rho-based recombination map from our Rabbit Slough genomes following a similar methodology to Schanfelter et al. (*44*), though with some minor modifications. Read-aware haplotype phasing was performed on the 20 Rabbit Slough whole genomes using the 3,577,312 SNPs using `ShapeIt` (v2.r837) (*45*) using the same parameters as Schanfelter et al.(*44*) (`--main 2000 --burn 200 --prune 210 --states 1000`).

We similarly constructed our ancestral sequence based on the *Gasterosteus wheatlandi* (*Gw*) re-sequencing from Yoshida et al. (*28*) and the *Pungitius pungitius* (*Pp*) re-sequencing from White et al. (*46*), except we mapped reads (after running `AdapterRemoval`) to our gasAcu1-4 reference genome using `bwa mem` (v0.7.15-r1140) (*23*) under default setting followed by `Stampy` (v1.0.32) (*47*) rather than `bowtie` (*48*). A substitution rate of 0.1 was used for Pp and 0.05 was used for Gw. The former was based on a 13 million year divergence time of GasAcu and Pp (*49*) and then assuming a yearly substitution rate of $7.1\times10^{-9}$ (*50*), and the latter set intermediate to this based on previously observed divergence (*28*) and the know topology (*51*). The combined `bwa-Stampy` approach resulted in substantially higher mapping sensitivity than the original mapping based on `Bowtie` (46% vs 69.6% for *Pp* and 74.2% v 87.2% *Gw*). Variants were called from gvcfs using `GATK Haplotypecaller` as described above. When constructing the ancestral sequence, if both outgroup species possessed alternative homozygote genotypes with a read depth of at least 3 and a genotype quality score of 30, then the ancestral allele was converted to this alternate allele. When only one outgroup species was callable and possessed an alternative homozygote genotype, if our Rabbit Slough genomes were variable for this same allele then the ancestral allele was converted to this alternate allele. The reconstructed ancestral sequence and associated files can be found here (https://drive.google.com/drive/folders/1tk2p0VMV2S212WXEW_Weqm8JPnSnkYQL?usp=shar

ing). Mutation matrices were constructed from this ancestral sequence and normalized as in Chan et al.(*52*).

Custom python scripts were used to construct the required input files for use by `ldhelmet` (v1.9) (*52*), utilizing the same workflow and parameters as Schanfelter et al. (*44*). Note that SNPs across all 20 Rabbit Slough genomes were used to calculate these maps, except for chrXIX, where we restricted our analysis to the 10 female samples. `find_confs` was run with a window size of `-w 50` to create a haplotype configuration file. To compute the likelihood lookup table `table_gen` was run with the grid of $\varrho$ values set to `-r 0.0 0.5 3.0 1.0 10.0` and $\boldsymbol{\theta}$ set to `-t 0.006`. This was estimated from a site frequency spectrum (sfs) generated for our 20 Rabbit Slough genomes by `angsd` (v0.922) (*53*), which utilizes genotype likelihoods to take into account SNP calling error and generate an sfs that is largely unbiased to variable coverage across genomes and positions. Repetitive DNA and transposons were masked when running `angsd`. Eleven padé coefficients were calculated using pade with `-t 0.006` and -`x 11`. A recombination map was then estimated by running rjmcmc with the following settings, `-w 50 -b 10.0 --burn_in 100000 -n 1000000`. The final $\varrho$-scaled recombination map was determined from the 50% percentile of the sampling distribution, with interpolation used to determine $\varrho$ between SNPs.

We compared our newly calculated Rabbit Slough (RS) map to the existing Lake Washington (LW) and Puget Sound (PS) maps from Schanfelter et al. (*44*) lifted over to GasAcu1-4. When comparing $\varrho$ estimates across 2k non-overlapping windows, we found a significant correlation between RS and both LW (r=0.295, p<<0.01) and PS (r=0.269, p<p<<0.01), though the correlation between LW and PS (r=0.433, p<p<<0.01) was unsurprisingly stronger (**Fig S17.3**). When comparing $\varrho$ visually across chromosomes via a LOWESS regression (conducted via the `statsmodels.nonparametric.lowess` function in python using 1% of the data for each y value and a delta of 0.01 × the number of 2kb windows per chromosome) we found that in general there was a high degree of similarity with regard to recombination landscape for all three populations (**Fig S17.4**).

$\varrho$-based maps were converted to genetic distance based on the pedigree-based linkage map generated by Glazer et al.(*54*) from genotyping-by-sequencing data. The map of Glazer et al. is very low resolution, based on 2,839 markers that correspond to regions that are each hundreds of thousands base pairs long and a previous build of the stickleback genome (BROADS1.75). We found a liftover to *gasAcu1-4* troublesome with many of these regions due to their large sizes. Therefore we located the corresponding region in our current reference genome for each marker by blasting the central 2000bp of each marker via `blastn`. We successfully identified the corresponding *GasAcu1-4* sequence for 879 markers. We then followed the procedure of Shanfelter et al.(*44*), which itself was based on Smukowski Heil et al. (*55*) to convert our map to genetic map units. Briefly, individual conversion factors are calculated for each chromosome by calculating the average cM/Mb rate across each individual marker-marker region and dividing this by the equivalent regions in the $\varrho$-based map. We find overall a high and significant correlation between our $\varrho$-based map and the linkage map of Glazer et al. (*54*) (r=0.853, p<<0.01) (**Fig S17.5**).

Recombination hotspots were identified based on the same criteria as Shanfelter et al.(*44*) with the mean $\varrho$ in 2kb windows sliding every 1kb compared to the mean rate from the 40kb flanking either side of the target window. We utilized thresholds of 5, 10 and 20x to determine whether a window was a local hotspot compared to its flanking region and merged any overlapping 2kb hotspot windows, resulting in 4,191, 1,941 and 864 windows respectively. This compares to the 4,265 windows identified by Shanfelter et al.(*44*) when using the 5k criteria.

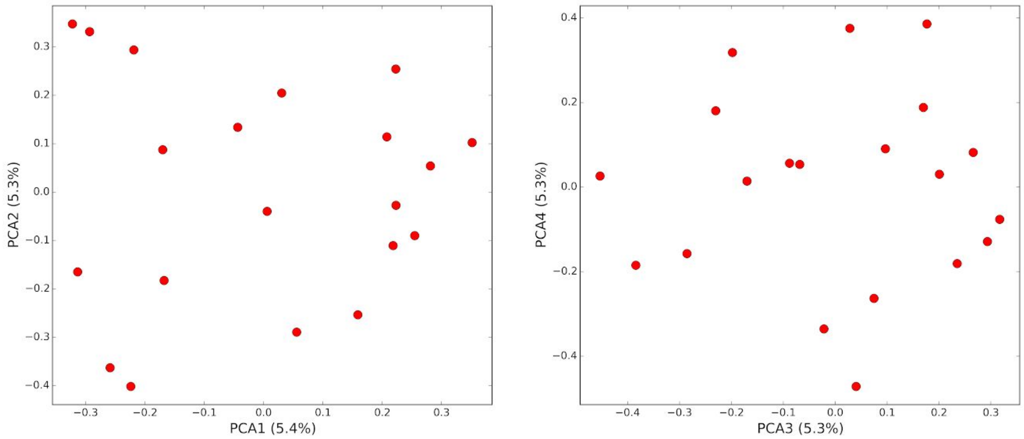


**Fig_S17.1:** First four PCs of PCA of Rabbit Slough whole genomes.

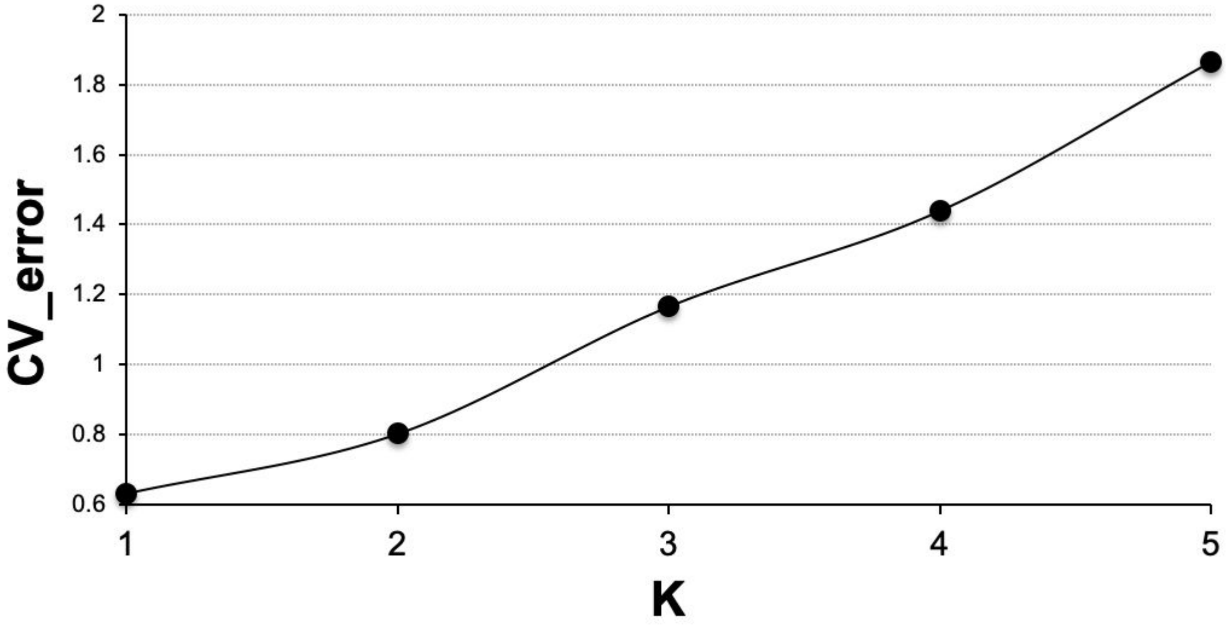


**Fig_S17.2:** Cross-validation analysis of model-based clustering analysis of Rabbit Slough whole genomes for first 5 Ks.

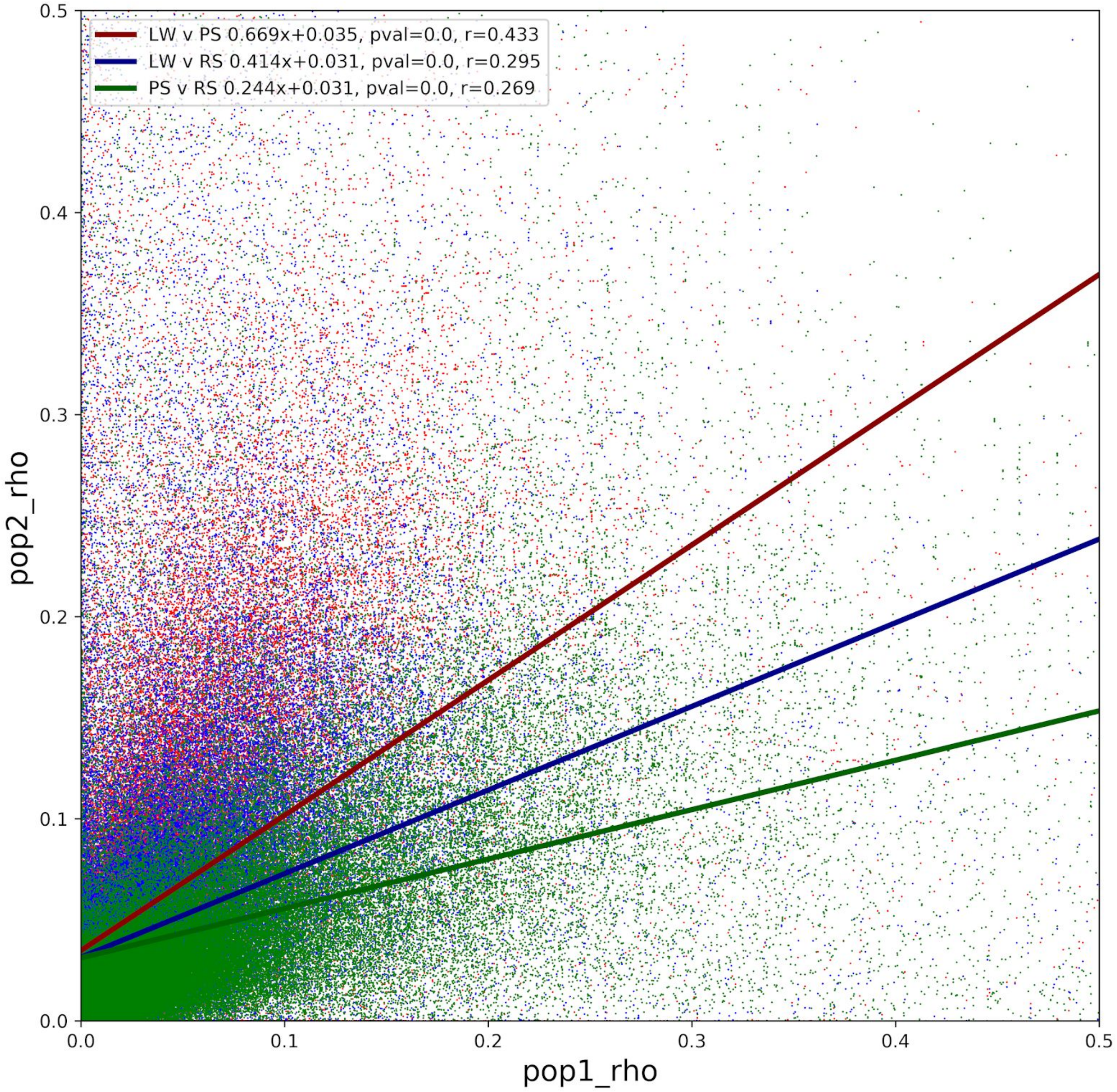


**Fig_S17.3:** Plot of correlation of estimated rho for 2kb windows across the entire genome for Rabbit Slough (RS), Lake Washington (LW) and Puget Sound (PS)

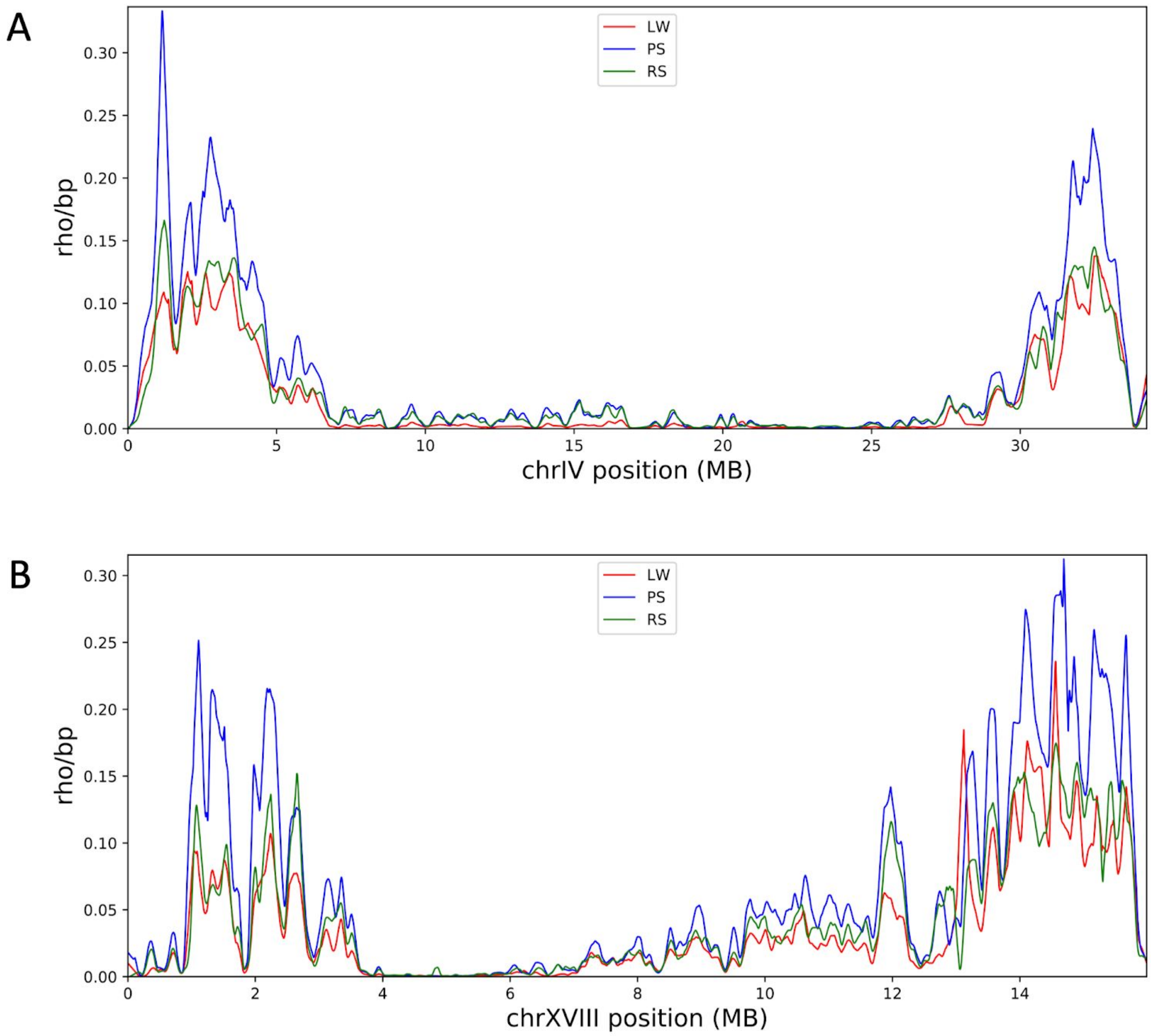


**Fig_S17.4:** Recombination landscape based on 2kb windows across chromosomes (A) IV and (B) XVIII for Rabbit Slough (RS), Lake Washington (LW) and Puget Sound (PS)

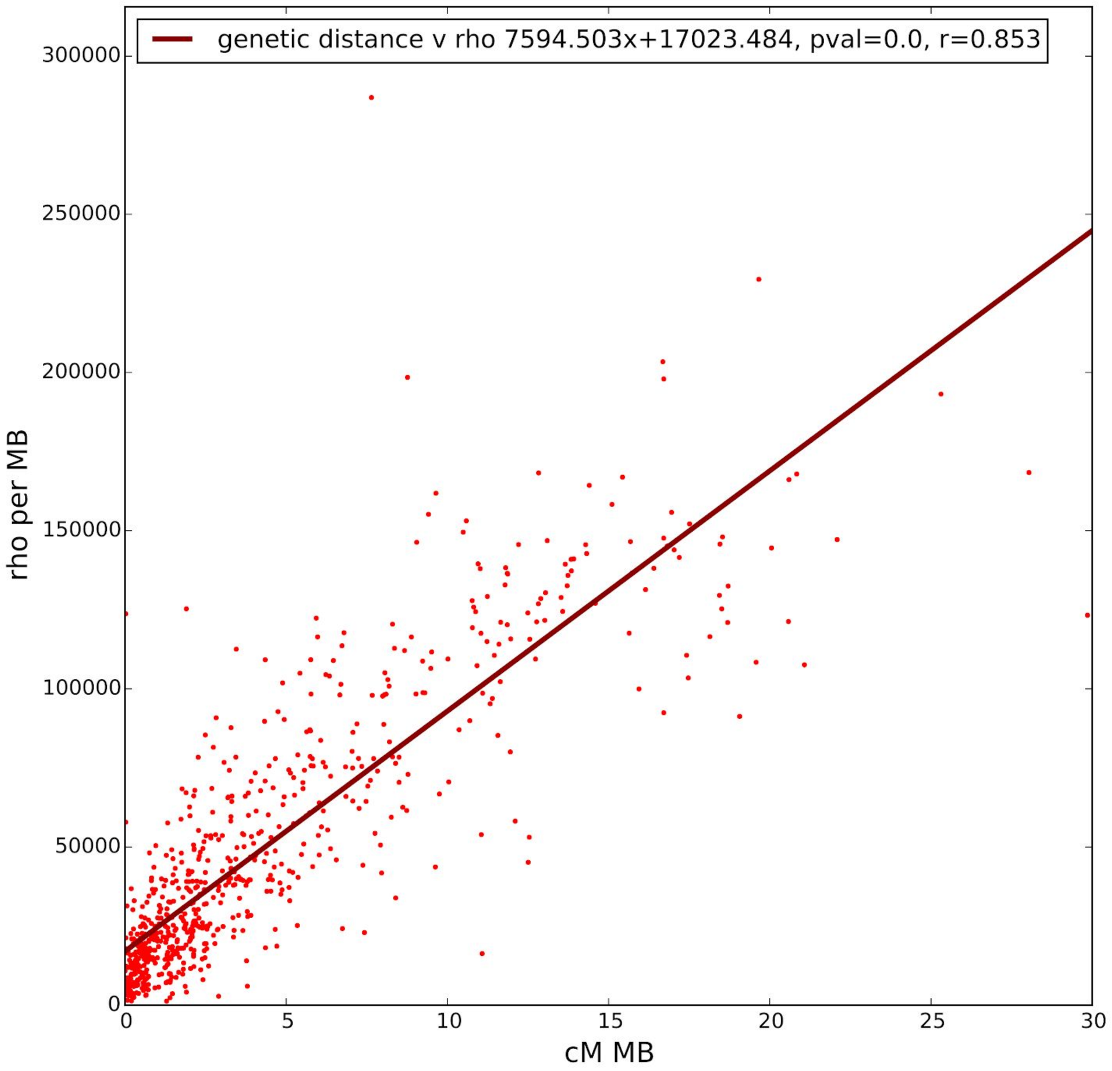


**Fig_S17.5:** Correlation between our rho-based map of recombination rate and the linkage map of Glazer et al.(*54*). Each point represents the genetic distance between two GBS markers from Glazer et al. and their equivalent value in our LD-based map.

### 18. $F_{ST}$ ANALYSIS OF OF CONTEMPORARY EVOLUTION SAMPLES

We estimated chromosome-specific $F_{ST}$ for each Pool-Seq population against its youngest counterpart. We utilised the ratio of averages method implemented in Bhatia et al (*56*) and based on Hudson et al.'s (*57*) estimator, applying this to SNPs ascertained in the Pool-Seq populations. We excluded chrXIX from this analysis due to uncertainty in chromosome number along its entire length.

When examining $F_{ST}$ between CH2009 and each subsequent Cheney Pool-Seq population, we see essentially three groups of chromosomes: those experiencing rapid evolution with $F_{ST}$ values increasing from 0 from the founding of the lake to above 0.1 after only 8 years (CH2017) (chrIV, chrXI, chrXXI, chrI, chrVIII), those experiencing intermediate evolution with final $F_{ST}$ > 0.03 (chrXII, chrXX, chrVIII, chrV, chrXIII, chrIX and chr X) and those experiencing slower evolution with final $F_{ST}$<0.025 (chrXVI, chrII, chrIII, chrVI, chrXVII, chrXV, chrXIV, chrXVIII) (**Fig S18.1**). As a point of comparison, we note that the $F_{ST}$ values we observe for the five rapidly evolving chromosomes after only 8 years are of similar magnitude to those observed between modern African and non-African human populations, which have probably evolved over at least 2000 generations (*56*).

We observe a similar overall magnitude of change when comparing SC2011 to each subsequent Scout Pool-Seq population (**Fig S18.2**, **Fig S18.3**). However, even though this time-series is two years behind Cheney, chrIV demonstrates an even higher final $F_{ST}$ in Scout 2017. In addition, chrI appears to be evolving somewhat slower and chrXII somewhat quicker in Cheney.

When comparing LB1999 to each subsequent Loberg Pool-Seq we see a much reduced rate of evolution, with the maximum $F_{ST}$ between LB1999 and LB2017 being ~0.035 (**Fig S18.4** and **Fig S18.5**). Contrasting Scout and Cheney with Loberg suggests that much of the chromosome evolution occurs within the first 10 years founding freshwater lakes with anadromous fish. However, it should also be noted that those chromosomes that change most rapidly in the first decade tend to also be the most rapidly evolving in the following two decades, albeit at a reduced pace. Indeed, there is a significant correlation ($p<0.01$) between both CH2009 v CH2017 $F_{ST}$ and SC2011 v SC2017 $F_{ST}$ compared to RS1999 v RS2017 $F_{ST}$. (**Fig S18.6** and **Fig S18.7**).

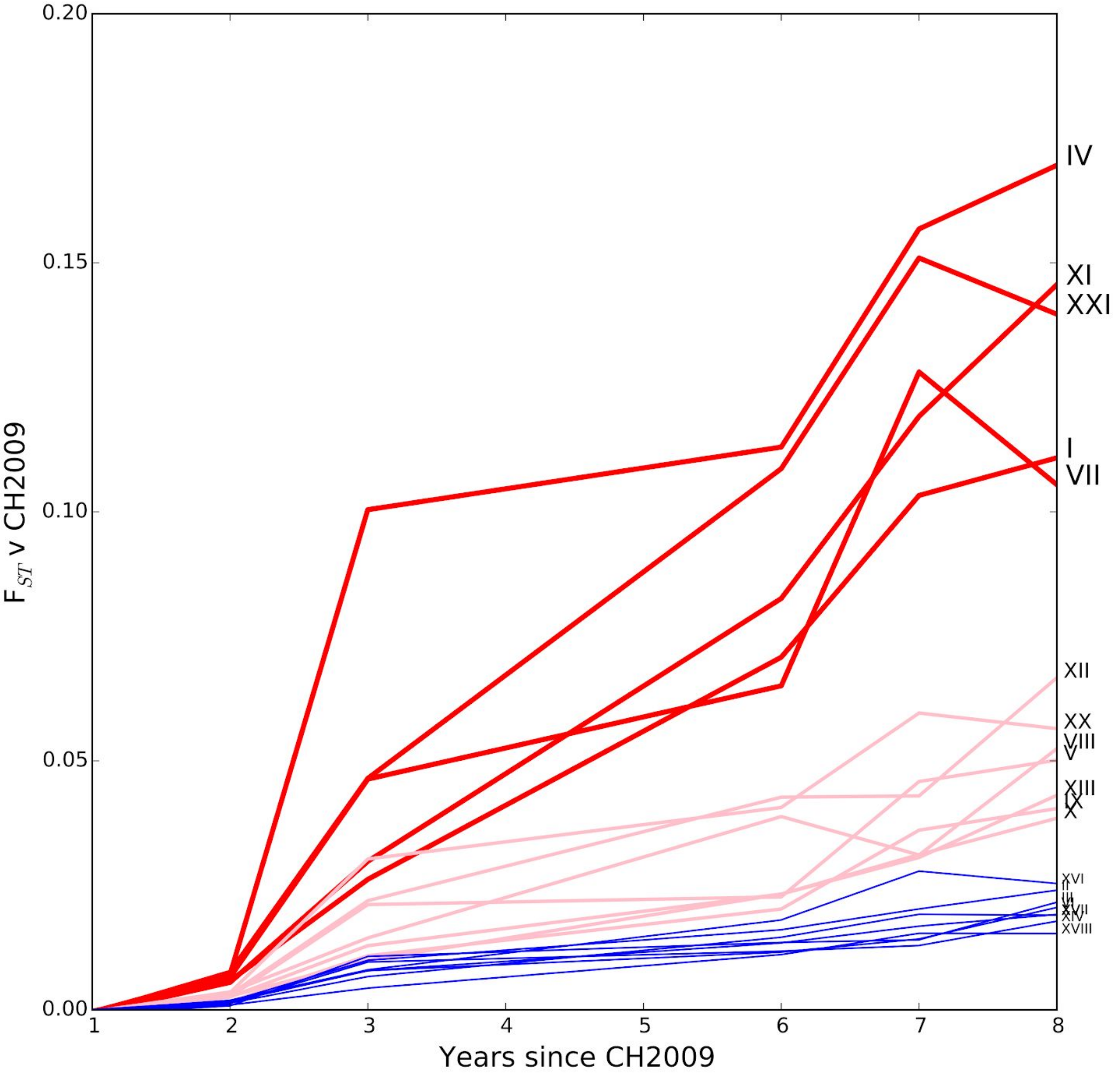


**Fig_S18.1**: Chromosome-specific pairwise FST when comparing CH2009 v to all subsequent Cheney Pool-Seq populations. Chromosomes defined as rapidly, intermediate and slowly evolving are coloured red, pink and blue respectively.

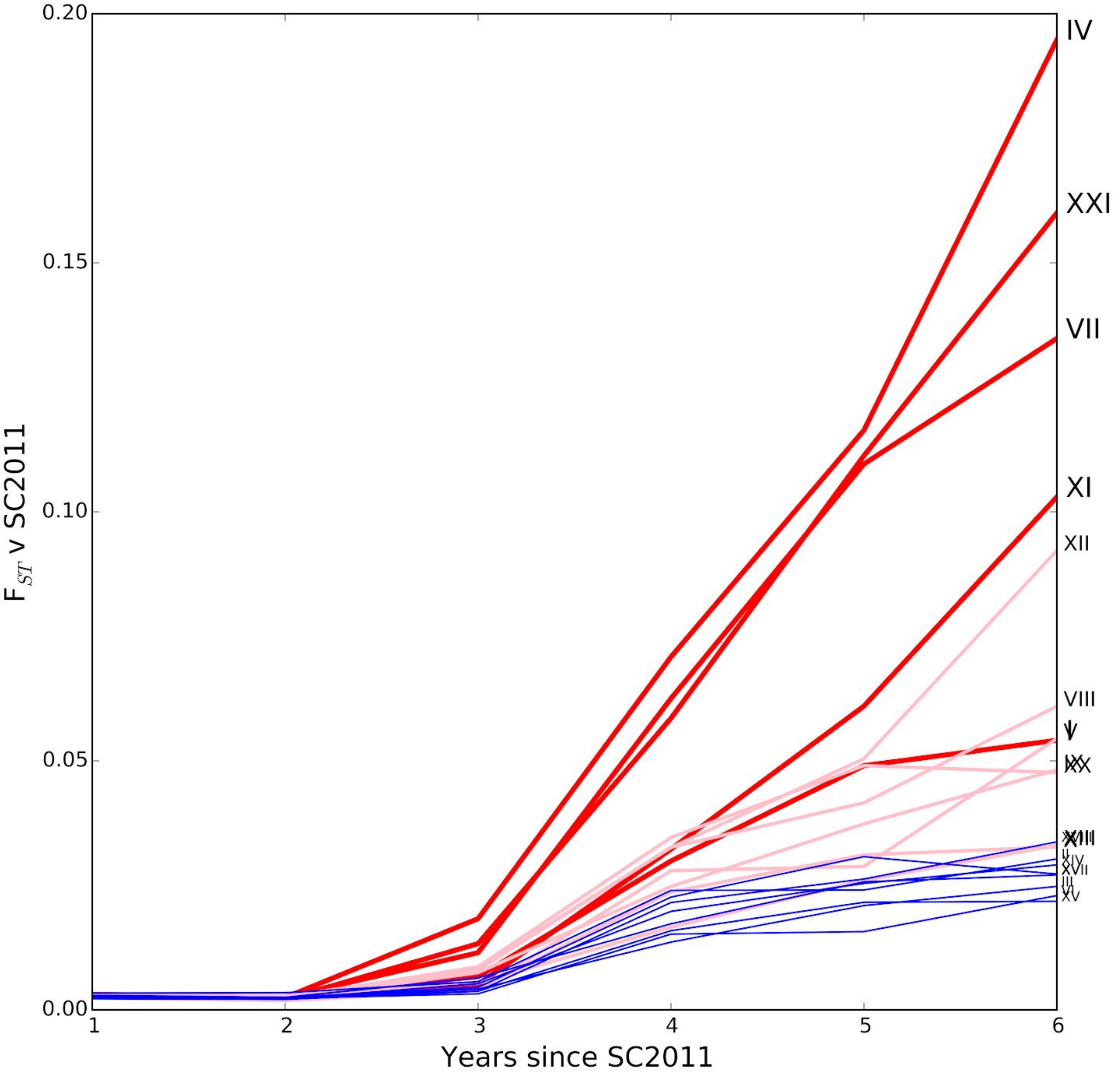


**Fig_S18.2**: Chromosome-specific pairwise FST when comparing SC2011 v to all subsequent Scout Pool-Seq populations. Chromosomes defined as rapidly, intermediate and slowly evolving in Chenery are coloured red, pink and blue respectively.

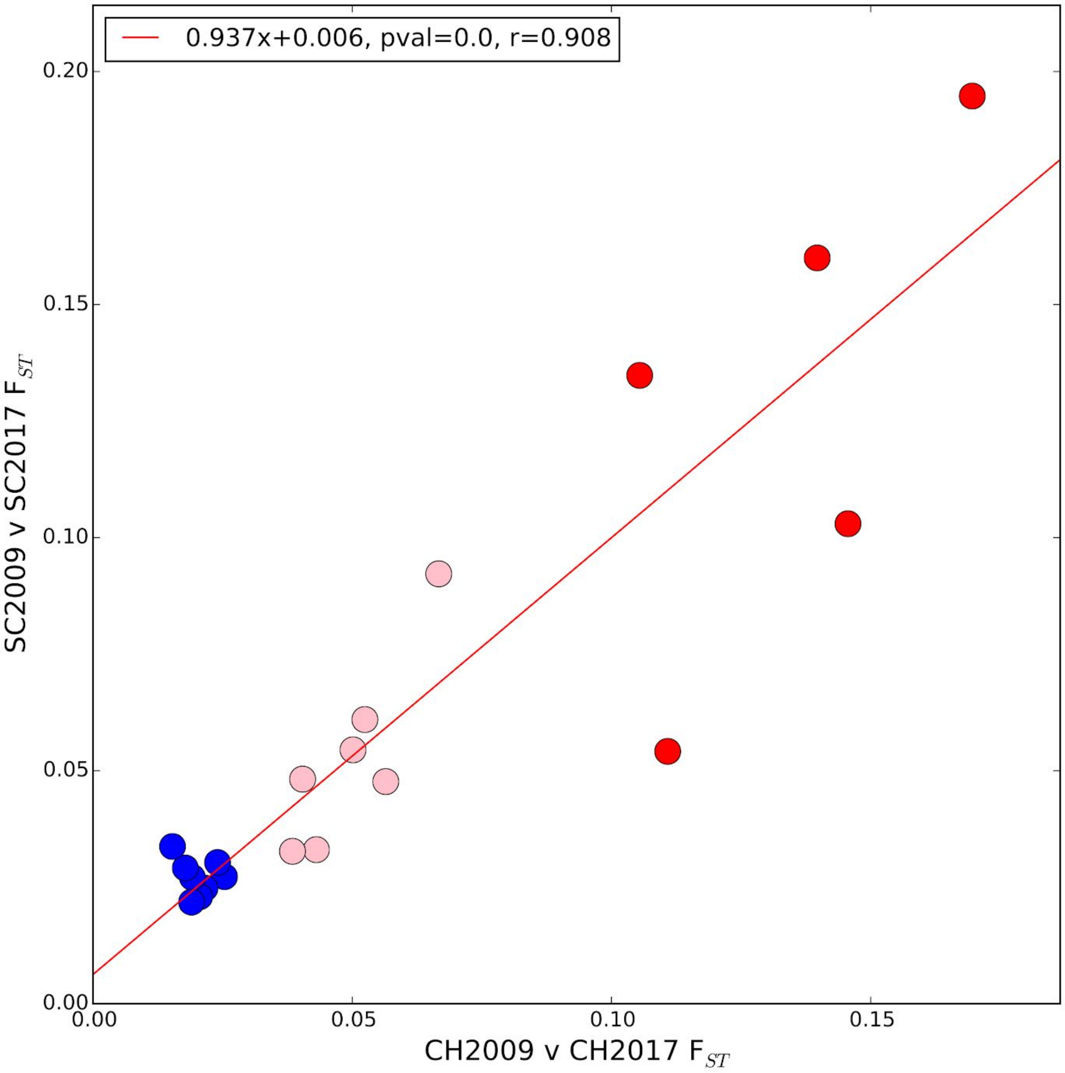


**Fig_S18.3**: Correlation of individual chromosome FSTs between first and last year sampled from Cheney and Scout Pool-Seq populations. Chromosomes defined as rapidly, intermediate and slowly evolving in Chenery are coloured red, pink and blue respectively. Red line represent best fit linear regression with inferred parameters shown in top left legend.

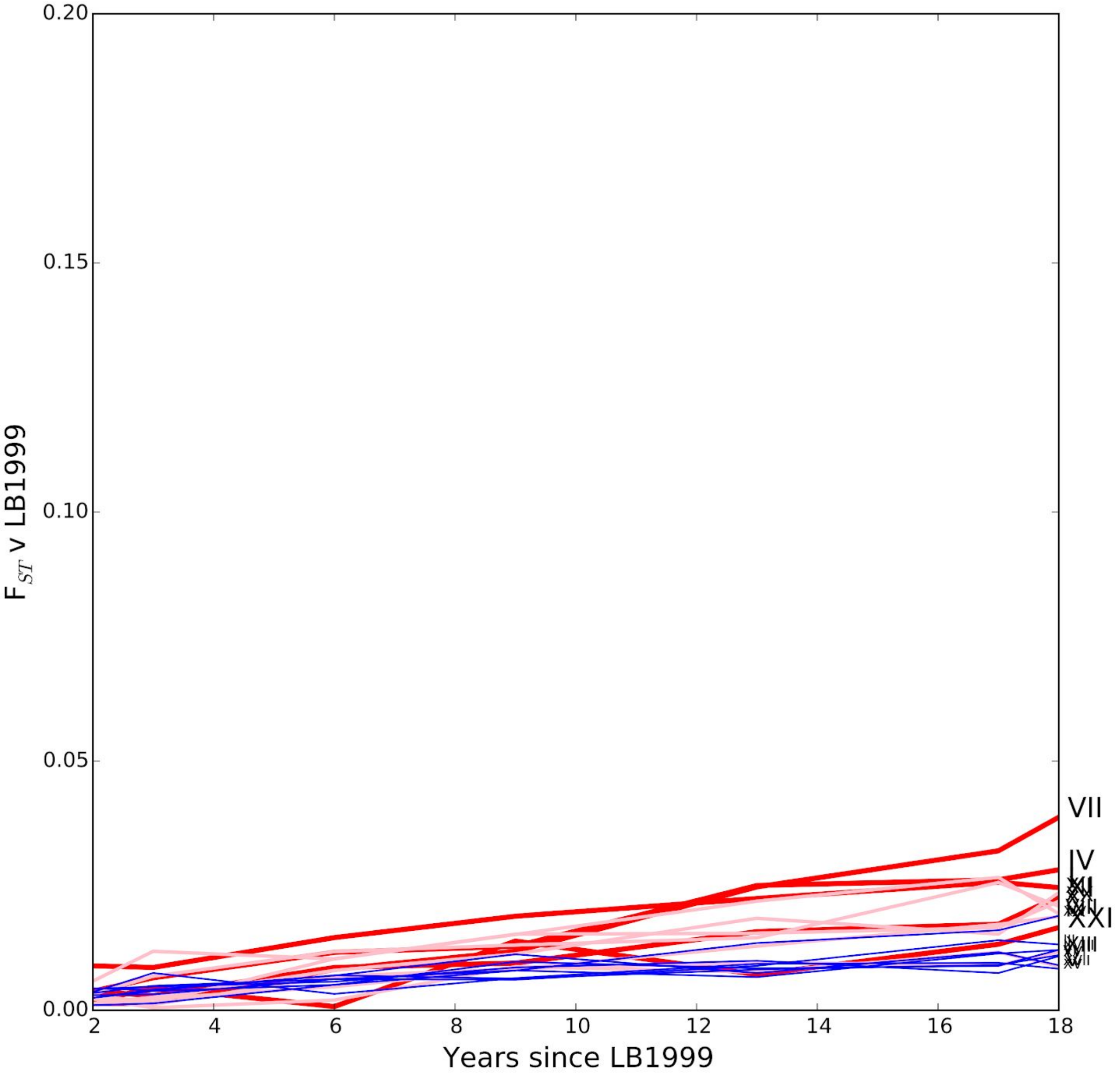


**Fig_S18.4**: Chromosome-specific pairwise FST when comparing LB1999 v to all subsequent Loberg Pool-Seq populations. Chromosomes defined as rapidly, intermediate and slowly evolving in Chenery are coloured red, pink and blue respectively. Scale of Y-axis matches Fig_S18.1.

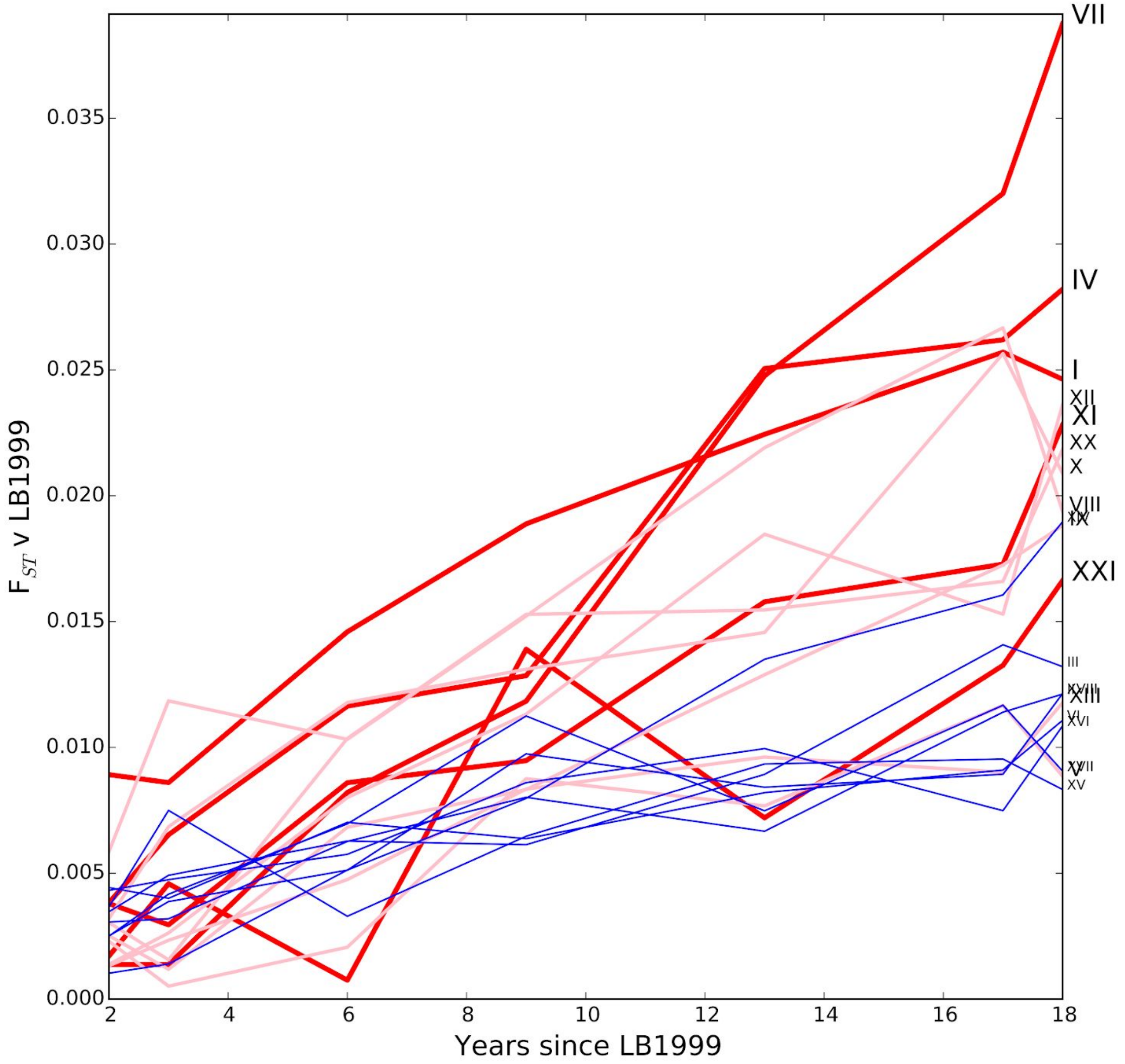


**Fig_S18.5**: Chromosome-specific pairwise FST when comparing LB1999 v to all subsequent Loberg Pool-Seq populations. Chromosomes defined as rapidly, intermediate and slowly evolving in Chenery are coloured red, pink and blue respectively. Scale of Y-axis is expanded compared to Fig_S18.4.

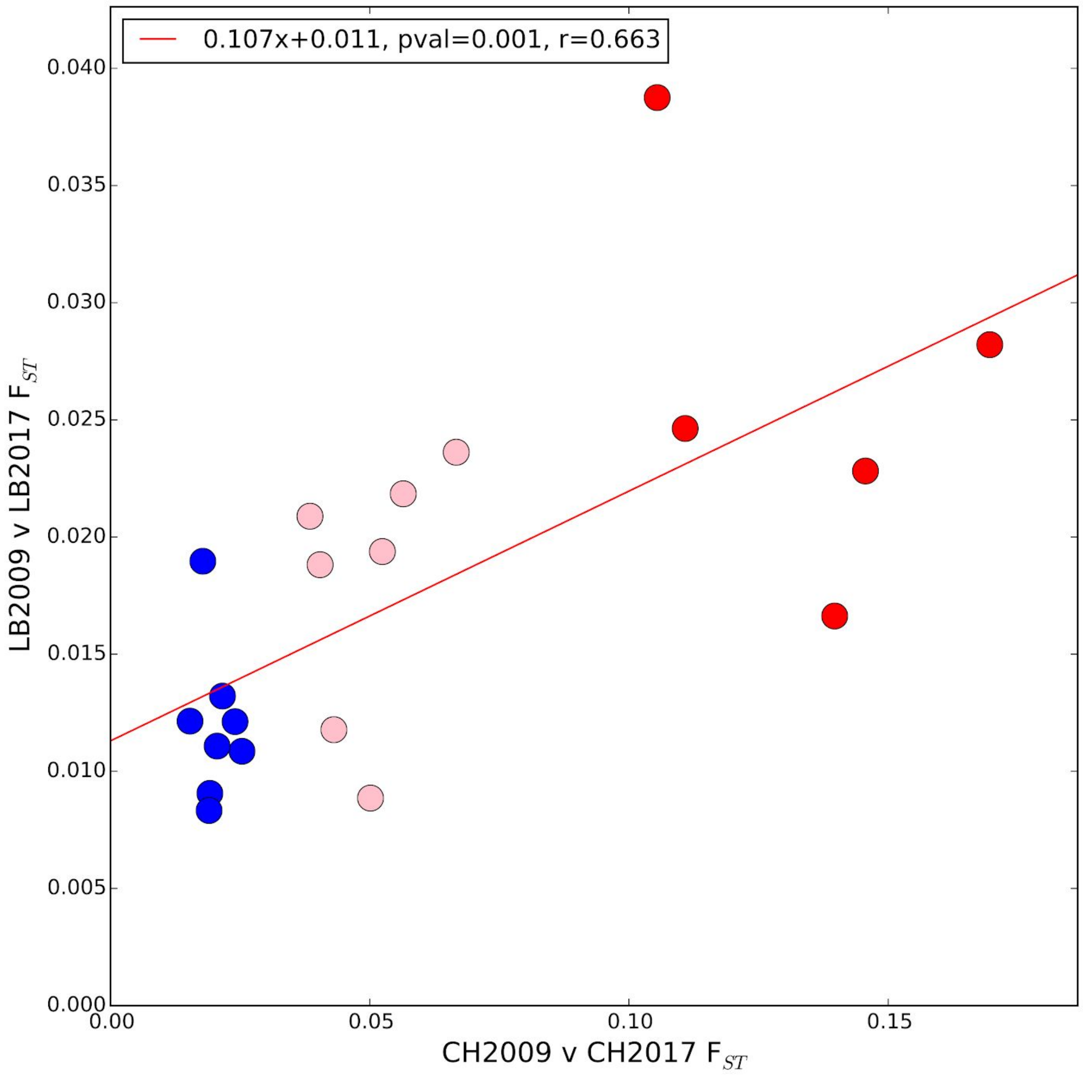


**Fig_S18.6**: Correlation of individual chromosome FSTs between first and last year sampled from Cheney and Loberg Pool-Seq populations. Chromosomes defined as rapidly, intermediate and slowly evolving in Chenery are coloured red, pink and blue respectively. Red line represent best fit linear regression with inferred parameters shown in top left legend.

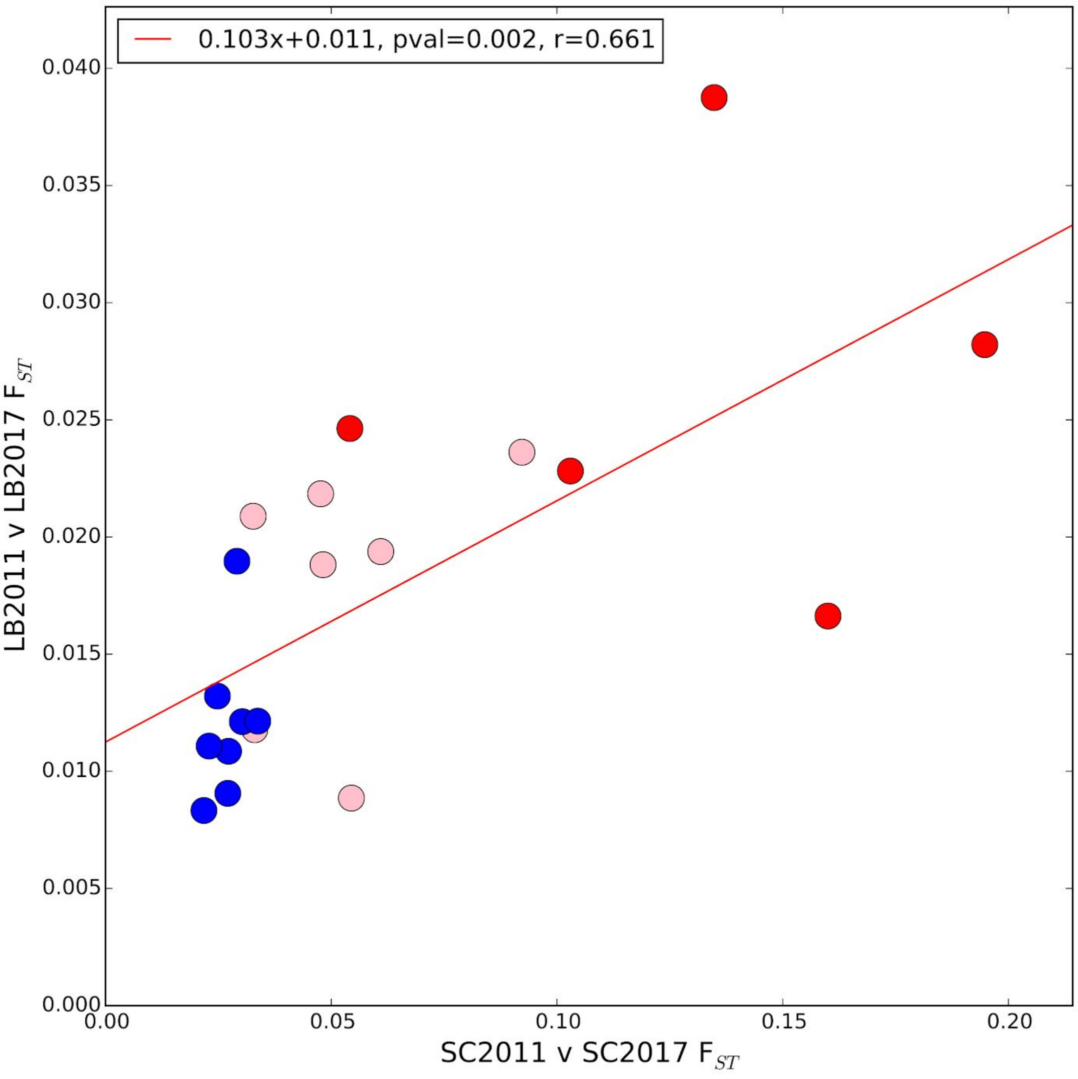


**Fig_S18.7**:. Correlation of individual chromosome FSTs between first and last year sampled from Scout and Loberg Pool-Seq populations. Chromosomes defined as rapidly, intermediate and slowly evolving in Chenery are coloured red, pink and blue respectively. Red line represent best fit linear regression with inferred parameters shown in top left legend

#### 19. $N_e$ INFERENCE FOR CONTEMPORARY EVOLUTION LAKES

Our time series data presented the opportunity to estimate changes in effective population size using the variance in allele frequencies between time points, with larger variance in allele frequency at neutral SNPs representing decreasing $N_e$ (i.e. increased drift). We used the method of moments estimator of Jonas et al. (*58*), which is well suited to Pool-Seq data as it models not only deviations from the underlying population allele frequency due to sampling, but also due to added variance caused by estimate the sample frequencies from variable read coverage across across samples. However, we do note that our use of the estimator does not account for any unevenness in DNA concentrations when pooling of samples, which will lead to further overdispersion. We could likely only confidently account for this using replicate Pool-Seq experiments of the same individuals, though effect on estimating allele frequencies is likely to be minimal given the high sample number and coverage of our pool (*59*). Regardless, during interpretation we focus on relative changes in $N_e$ rather than absolute values, which would be underestimated due to any unaccounted for overdispersion.

We applied this method to all pairs of successive Pool-Seq experiments (so CH2009 v CH2010, CH2010 v CH2011, etc.), utilizing biallelic SNPs ascertained in the long term samples and with a minor allele frequency of at least 5% in the oldest population of a pair and calculating separate estimates for each chromosome.Given that $N_e$ will be affected by any positive or negative selection, we in particular played closest attention to chrXV, which shows the least evidence of rapid freshwater adaptation and thus in theory should reflect shifts in allele frequency due to genetic drift and thus population demographic effects.

When examining our time series via boxplots where each point is the $N_e$ for individual chromosomes, we observe that both Cheney and Scout show a decrease and then increase in $N_e$, with harmonic means for chrXV across the entire time series of 150 and 120 respectively (**Fig S19.1** and **Fig S19.2**). Interestingly, these dips in $N_e$ show a marked concordance to Catch Per Unit Effort estimates for these two lakes from Bell et al. (*7*), a proxy for estimating the census population size of the lake (**Fig S19.1** and **Fig S19.2**). This suggests that population decreases early on in the history of these two new freshwater are affecting genetic variation in a corresponding manner. The Loberg time series shows a somewhat larger overall $N_e$ (chrXV harmonic mean of 250), with some evidence of a small increase over time until recently, though there is considerable variation across chromosomes (**Fig S19.3**).

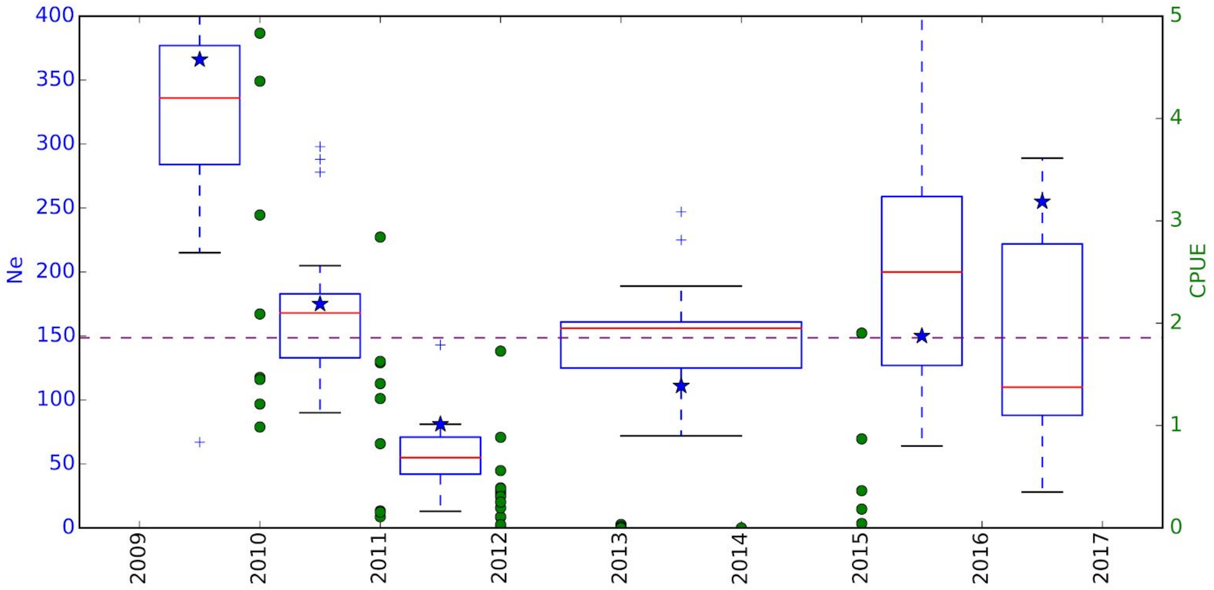


**Fig S19.1:** Left axis (blue): Boxplots of chromosome-specific estimates of Ne between successive Pool-Seq experiments at Cheney Lake. Blue star shows estimates for chrXV. Purple line represents harmonic mean for chrXV. Note that we have reduced the Y-axis to exclude sections above the interquartile for some paired years to ease viewability. Right axis (green): Estimates of catch per unit effort (CPUE) via fish per trap-hour taken from Bell et al. 2016 (*7*).

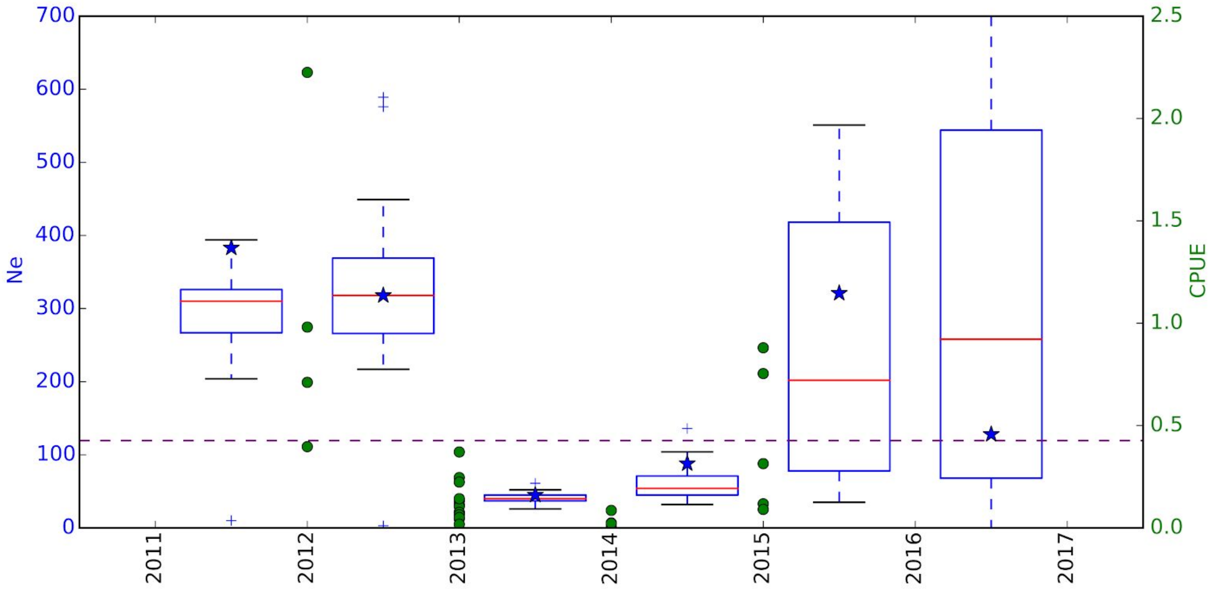


**Fig S19.2:** Left axis (blue): Boxplots of chromosome-specific estimates of Ne between successive Pool-Seq experiments at Scout Lake. Blue star shows estimates for chrXV. Purple line represents harmonic mean for chrXV. Note that we have reduced the Y-axis to exclude sections above the interquartile for some paired years to ease viewability. Right axis (green): Estimates of catch per unit effort (CPUE) via fish per trap-hour taken from Bell et al. 2016 (*7*).

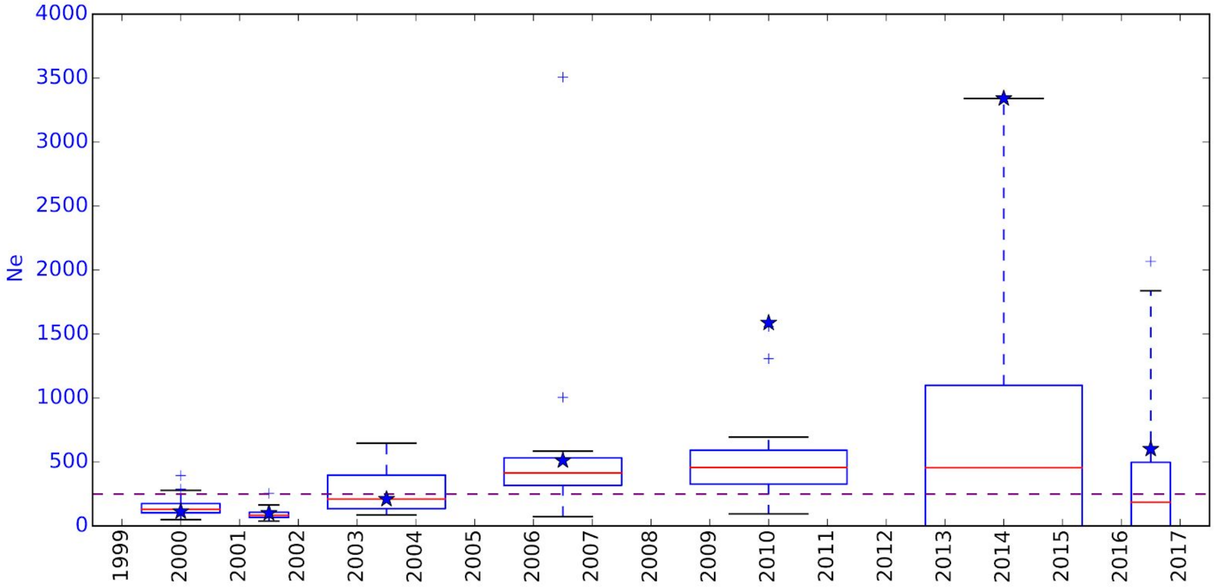


**Fig S19.3:** Boxplots of chromosome-specific estimates of Ne between successive Pool-Seq experiments at Loberg Lake. Blue star shows estimates for chrXV. Purple line represents harmonic mean for chrXV. Note that we have reduced the Y-axis to exclude sections above the interquartile for some paired years to ease viewability.

### 20. CMH ANALYSIS OF CONTEMPORARY EVOLUTION SAMPLES

In order to identify SNPs that had shown a significant change in allele frequency in our contemporary time-series data, we applied a modified Cochran–Mantel–Haenszel (CMH) test (*60*). CMH tests are commonly used in evolve and resequencing (E&R) experiments when comparing replicate Pool-Seq samples from two different time points (versus a Fisher's exact test which only examines a single replicate) and show good power compared to many other tests (*61*). The modified CMH test, in similar vein to the $N_e$ analysis described above, additionally accounts for the overdispersion caused by sampling and sequencing, while also controlling for the expected neutral drift between two time points, which better controls the resulting p-value distribution. This method does not account for the effect of uneven pooling of samples. However, in modelling drift in our modified CMH test, we utilize for each lake the harmonic mean of $N_e$ estimates based on each successive pair of time points (Loberg = 250, Cheney = 150, Scout = 120). As these $N_e$ estimates also did not account for possible uneven pooling, their values may in fact somewhat reflect the added overdispersion caused by it.

We applied this test to five combinations of lakes: all three lakes together (CH+SC+LB), Cheney and Scout together (CH+SC, this would reflect significant changes occurring within the earliest stage of freshwater adaptation), as well as each lake on its own (LB, CH, SC). We note that the first two combinations do not strictly represent replications as usually applied in evolve and resequence experiments, as each lake is sampled over different lengths of time, which the original code does not allow. We rewrote the code in python, allowing a different level of drift ($N_e$ ) for each individual lake when calculating the test statistic, as well as different time intervals. For Cheney and Scout we compared the earliest Pool-Seq time point (CH2009 and SC2011) to the latest time point (CH2017 and SC2017). In the case of Loberg for which we do not know the founding population precisely, we utilize RS2009 as a proxy for the starting population and LB2017 as the end point. As described above, Rabbit Slough is the most likely source of anadromous fish after Loberg was poisoned.

For each lake combination a modified CMH test statistic and associated p-value was calculated for any SNP identified in the established population set that for each Pool-Seq population had coverage between one-half and two times the mean coverage, and that was variable in at least one Pool-Seq population considered. The number of SNPs used ranged between 3,023,631 (CH+SC+LB) and 3,682,352 (LB) across the five combinations. We used a Bonferroni correction to identify significant SNPs at a 5% threshold.

For CH+SC+LB the number of significant SNPs is greatest on chrIV (n=46,215) but also noticeably prominent on chrVII (n=31,083) and XXI (n=28,299), while chrI, chrXI, chrXII and chrXX are somewhat more moderate. Barely any (<40) significant SNPs are found on chrVI, chrXV and chrXVIII (**Fig S20.1**). LB analyzed on its own largely reiterates the relative number of significant SNPs as CH+SC+LB, presumably reflective of it being the most evolved freshwater population with regards to time (**Fig S20.2**). It is noticeable that SC and CH analyzed separately show somewhat distinctive patterns, with chrIV also dominant when analyzing SC (**Fig S20.3**) but chrXI is more prominent for CH (**Fig S20.4**). Combining the signal for CH+SC shows the importance of chrIV in early freshwater adaptation, consistent with the genome-wide $F_{ST}$ results (**Fig S20.5**). As would be expected, there are more significant SNPs when utilizing multiple populations (e.g. CH+SC+LB) or populations with a long time-series (e.g. Loberg) in the CMH test.

Examining the distribution of CMH p-values across the genome using both physical and genetic distance (based on our Rabbit Slough-based recombination map), we see that our significant p-values cluster into clear peaks, while also noting that for chromosomes with large numbers of significant SNPs, using genetic distance tends to reduce the span of these peaks (i.e. they are much closer in genetic space compared to physical space) (**Fig S20.6-10**).

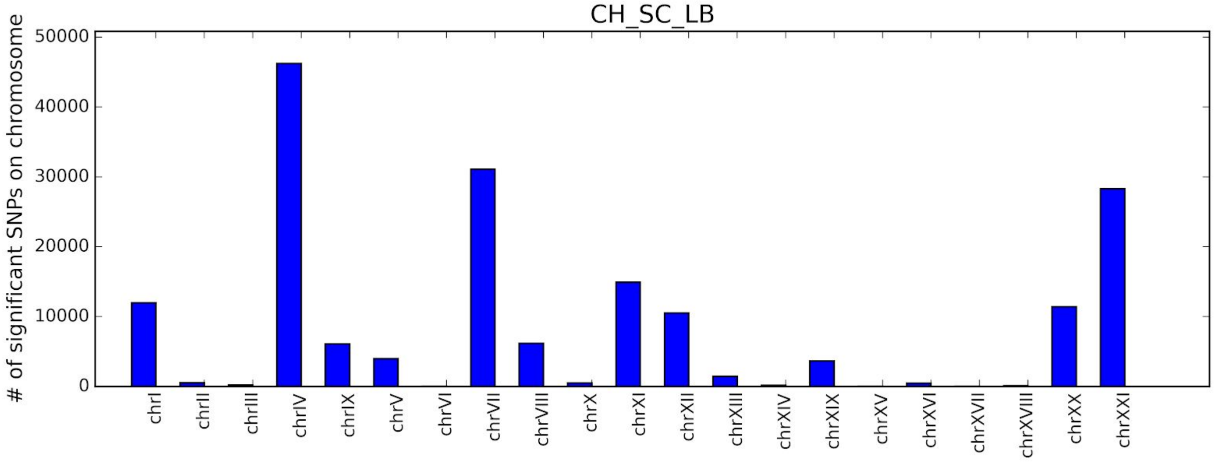


**Fig S20.1:** Number of significant SNPs for modified CMH test after Bonferroni correction for CH+SC+LB for each individual chromosome.

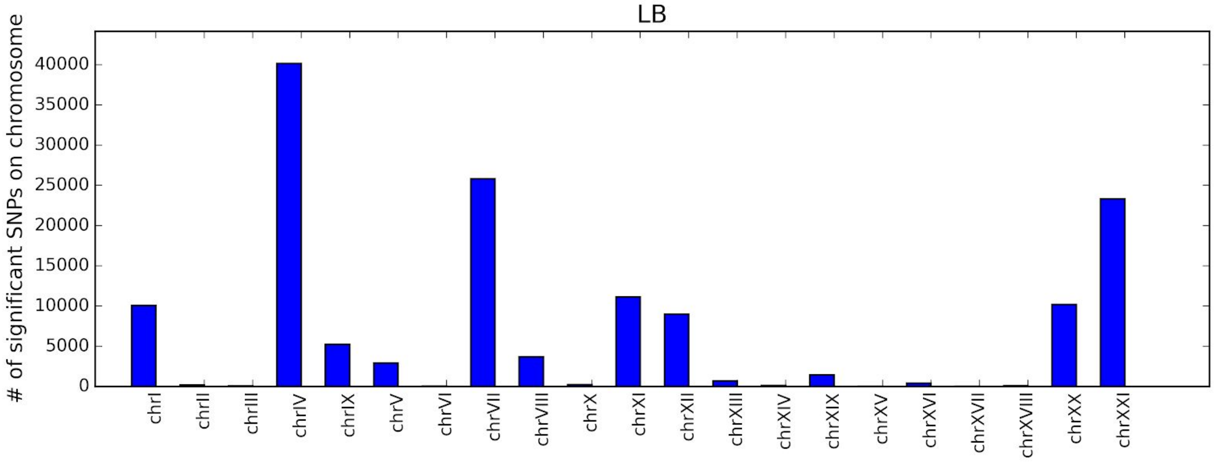


**Fig S20.2:** Number of significant SNPs for modified CMH test after Bonferroni correction for LB for each individual chromosome.

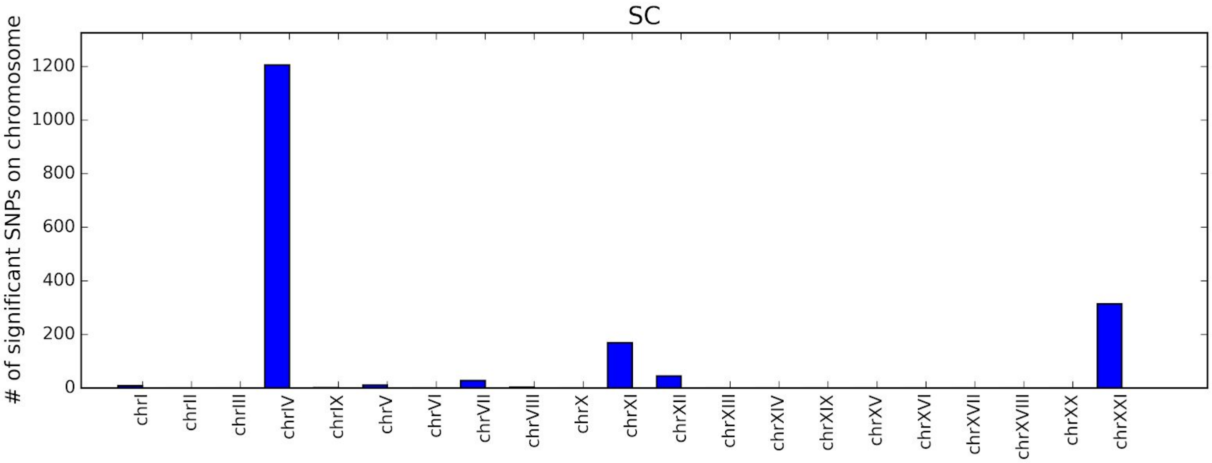


**Fig S20.3:** Number of significant SNPs for modified CMH test after Bonferroni correction for SC for each individual chromosome.

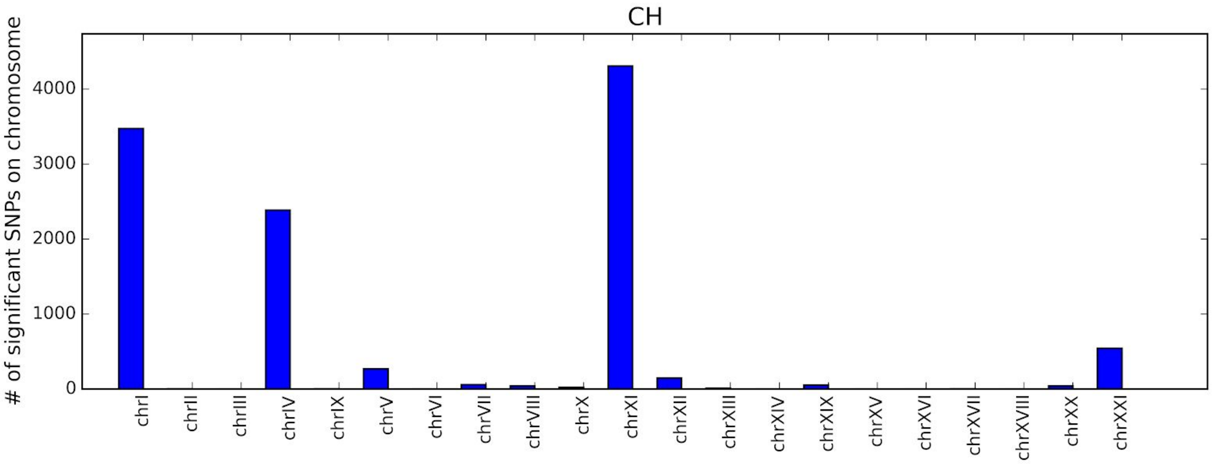


**Fig S20.4:** Number of significant SNPs for modified CMH test after Bonferroni correction for CH for each individual chromosome.

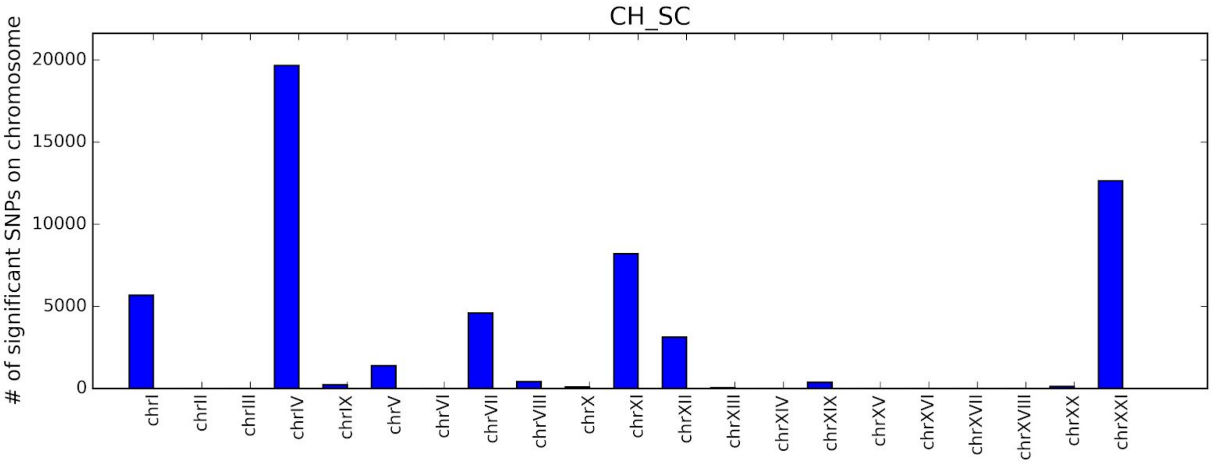


**Fig S20.5:** Number of significant SNPs for modified CMH test after Bonferroni correction for CH+SC for each individual chromosome.

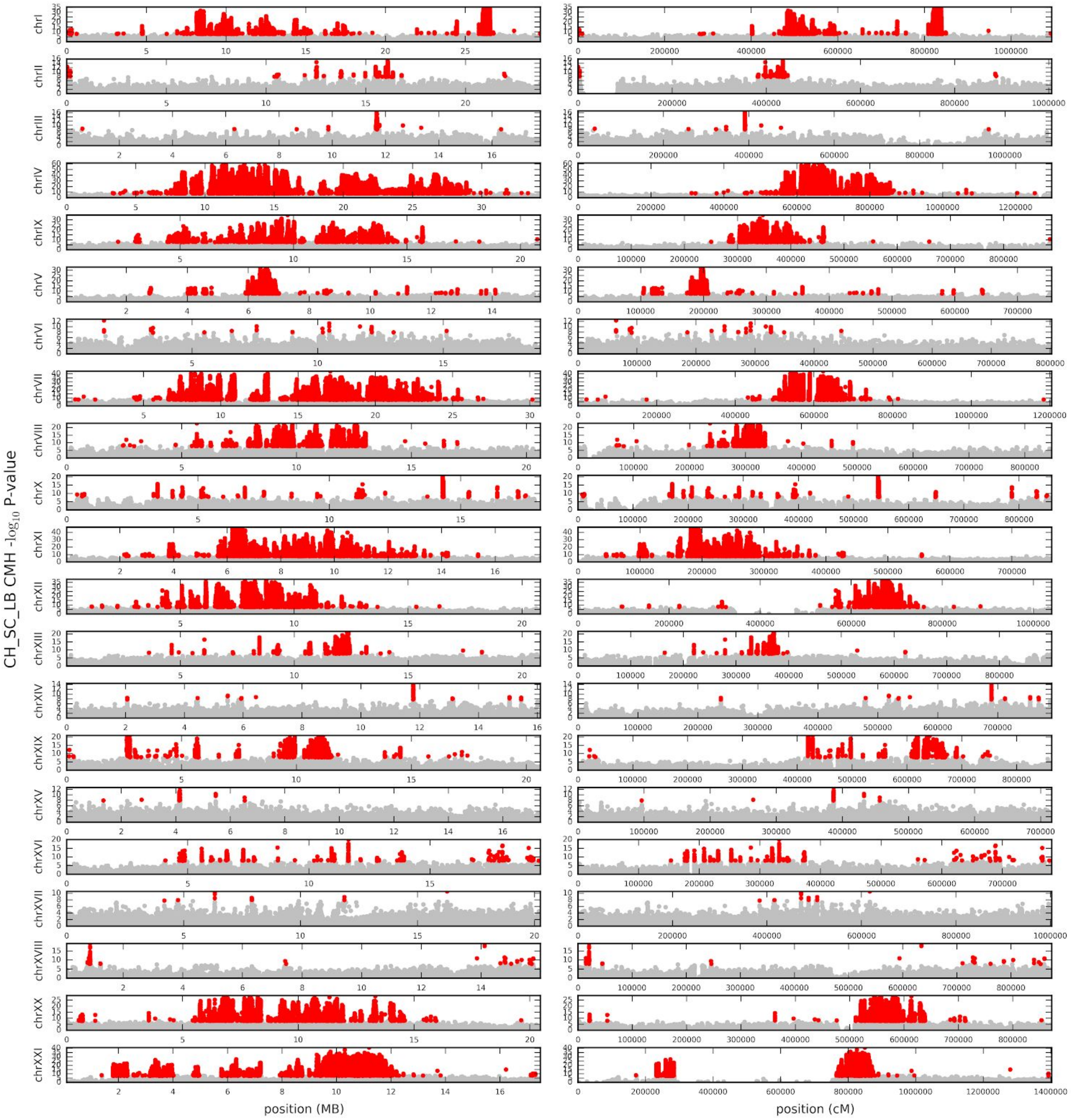


**Fig S20.6:** Genome-wide distribution of CMH -log10 p-values for each chromosome for CH+SC+LB, with position in physical distance (left column) and genetic distance (right column). Red circles represent significant SNPs after Bonferroni correction, silver circles represent all other SNPs. Note that Y-axis is scaled for the lowest p-value for each chromosome separately.

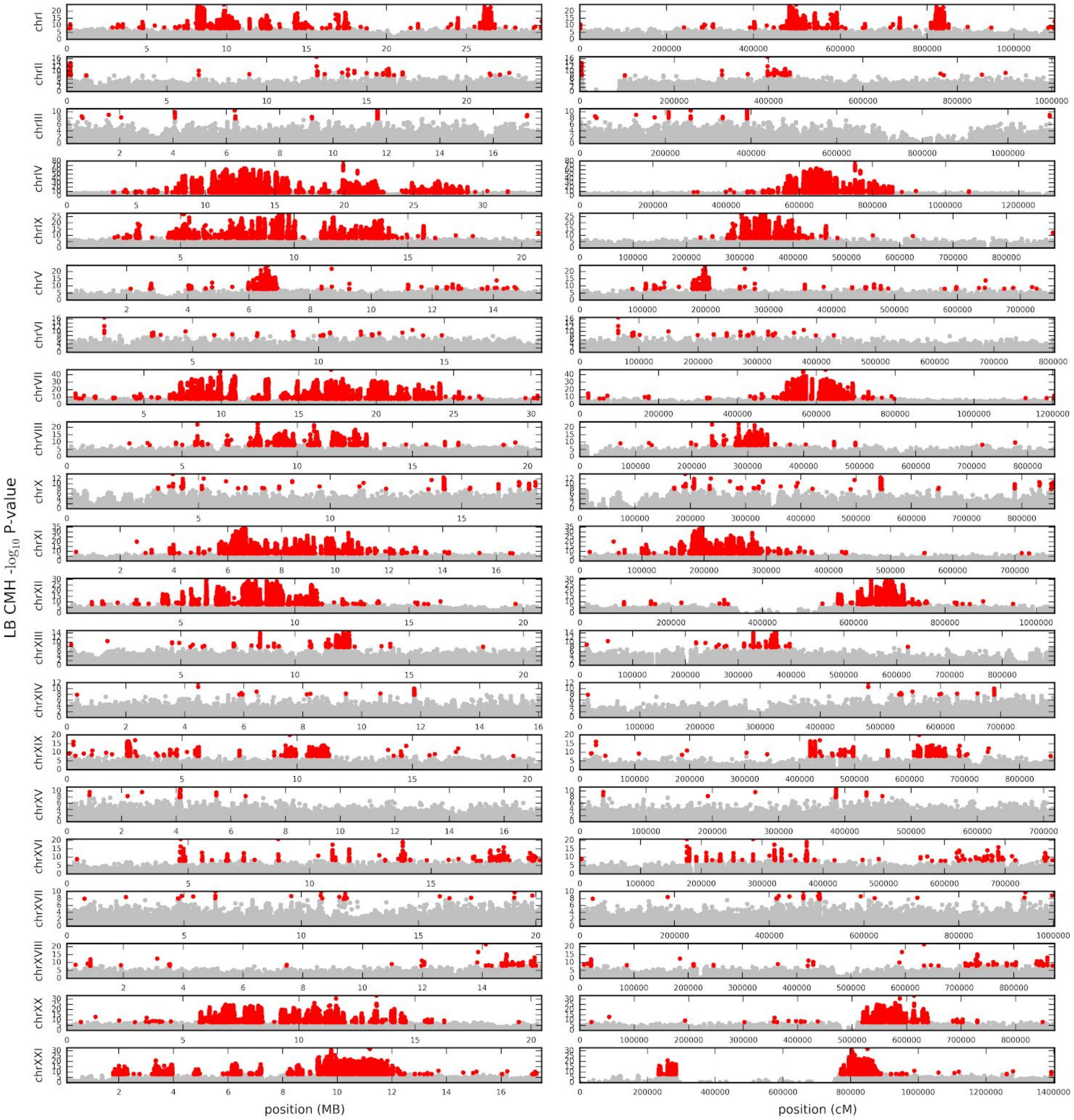


**Fig S20.7:** Genome-wide distribution of CMH -log10 p-values for each chromosome for LB, with position in physical distance (left column) and genetic distance (right column). Red circles represent significant SNPs after Bonferroni correction, silver circles represent all other SNPs. Note that Y-axis is scaled for the lowest p-value for each chromosome separately.

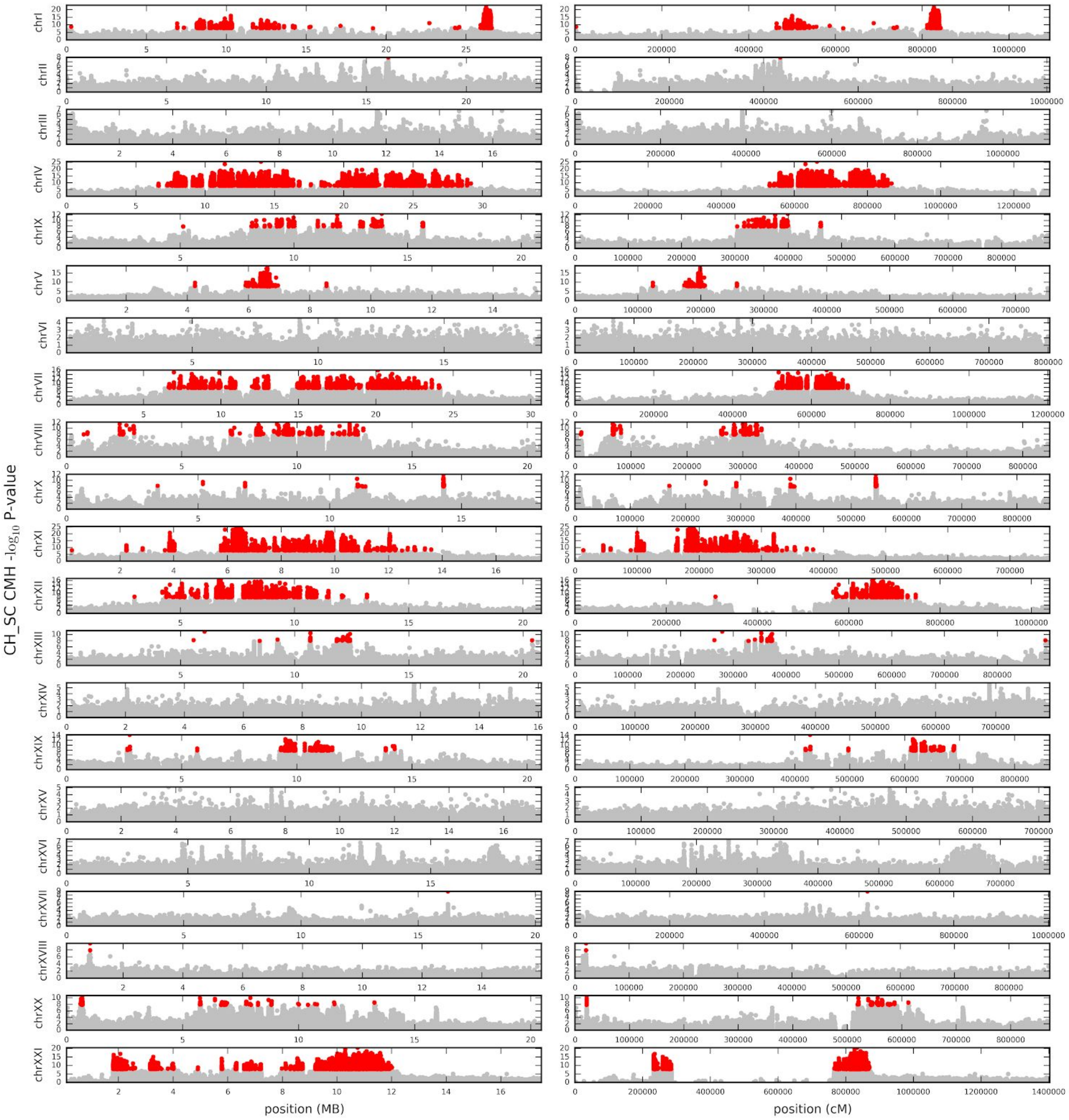


**Fig S20.8:** Genome-wide distribution of CMH -log10 p-values for each chromosome for CH+SC, with position in physical distance (left column) and genetic distance (right column). Red circles represent significant SNPs after Bonferroni correction, silver circles represent all other SNPs. Note that Y-axis is scaled for the lowest p-value for each chromosome separately.

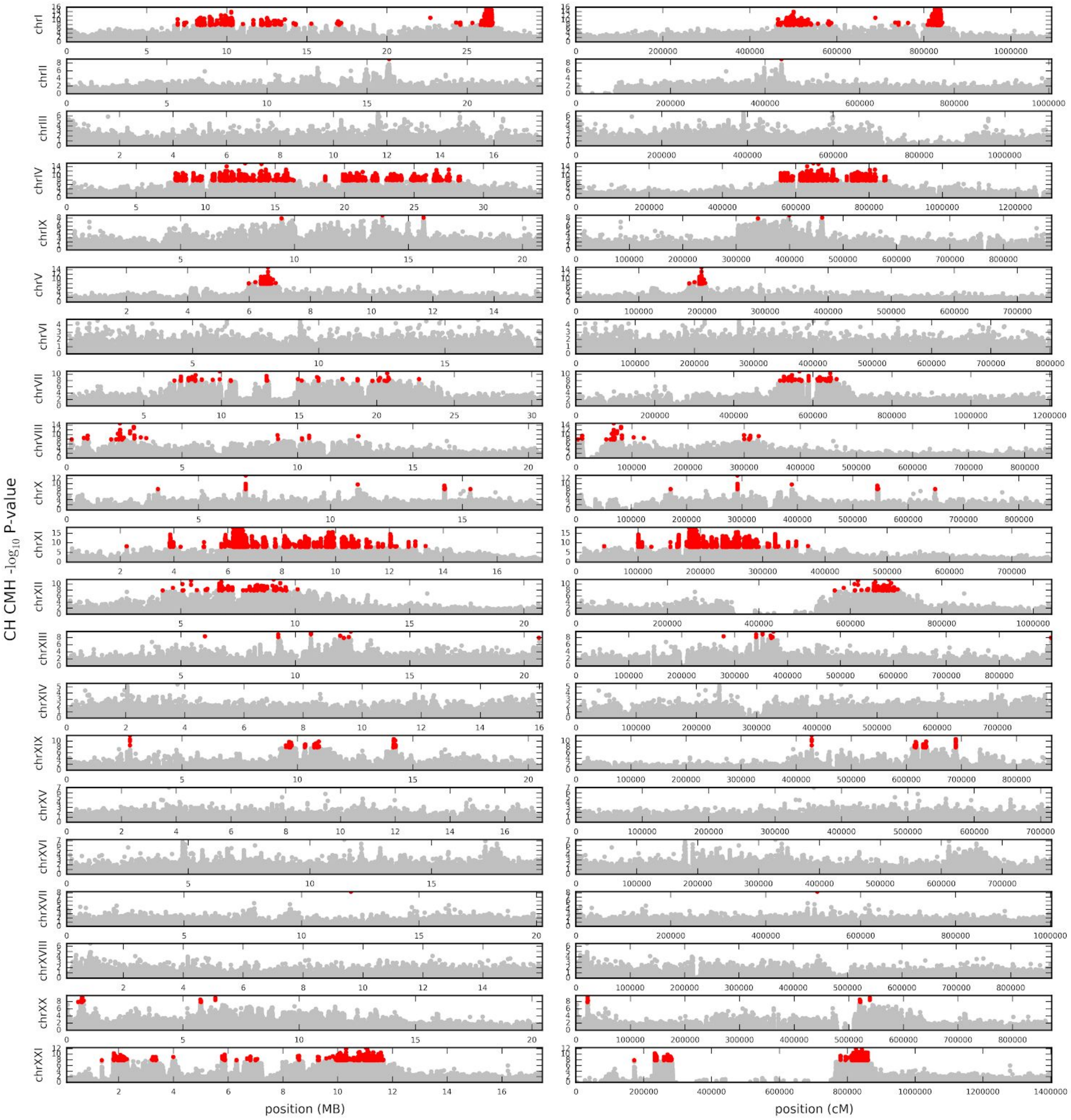


**Fig S20.9:** Genome-wide distribution of CMH -log10 p-values for each chromosome for CH, with position in physical distance (left column) and genetic distance (right column). Red circles represent significant SNPs after Bonferroni correction, silver circles represent all other SNPs. Note that Y-axis is scaled for the lowest p-value for each chromosome separately.

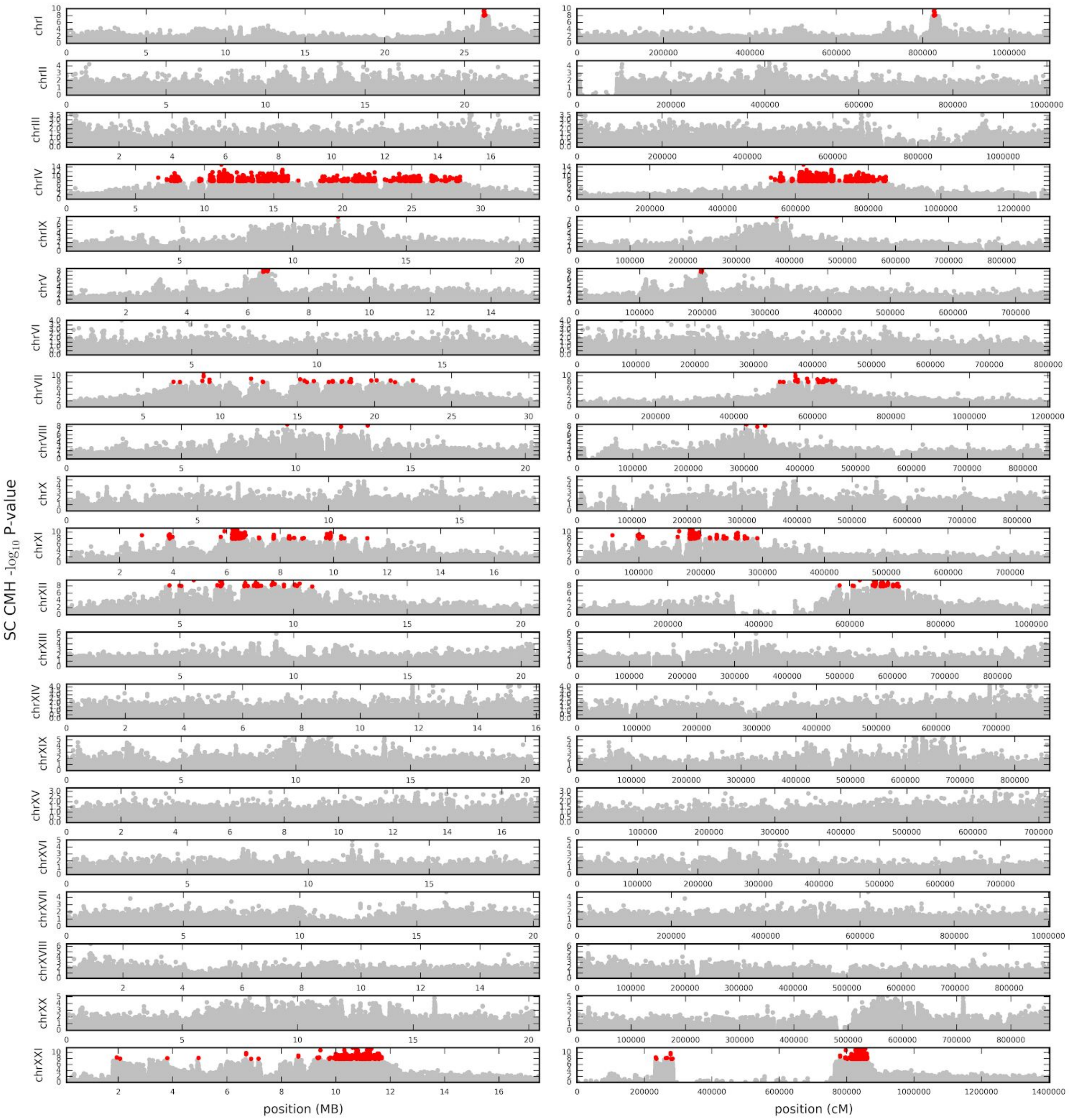

**Fig S20.10:** Genome-wide distribution of CMH -log10 p-values for each chromosome for SC, with position in physical distance (left column) and genetic distance (right column). Red circles represent significant SNPs after Bonferroni correction, silver circles represent all other SNPs. Note that Y-axis is scaled for the lowest p-value for each chromosome separately.

### 21. TEMPOPEAK IDENTIFICATION

We followed the same general EcoPeak identification methodology in order to define TempoPeaks from our CMH p-values. However, we restricted our peaks to those called using a Bonferroni correction to determine the minimum p-value peak threshold, which showed a good overlap with long term peaks. We found using a false discovery rate to be too permissive compared to calling EcoPeaks. We produced a set of sensitive and specific TempoPeaks for each of the five lake combinations (CH+SC+LB, CH+SC, LB, CH, SC). Sensitive TempoPeaks were based on a 5% Bonferroni-corrected p-value threshold merging SNP and window-defined peaks. Specific TempoPeaks were based on a 1% p-value threshold only considering window-defined peaks. A separate set of peaks was also constructed using physical and genetic distance to define windows and peaks. Based on our overall genetic map, the approximate recombination rate per generation per base is $4 \times 10^{-6}$, and thus the window size and sliding window size of 2500bp and 50bp is equivalent to 0.01cM and 0.0002cM.

Unsurprisingly, the most TempoPeaks are identified using all three contemporary lakes together, where we will have maximum power, or using Loberg, where the most time has passed since the founding of the population, while the least number of TempoPeaks are found when only examining Scout, where the least time has passed (**Table S21.1**). No significant specific TempoPeaks were found for Scout at all, though there were observable peaks in CMH scores that line up with significant TempoPeaks identified in other lake combinations (i.e. there is insufficient power to call them significant beyond the background of the rest of the genome at this stage of the lake's evolution) (**Fig S21.1 - Fig S21.10**).

As reflected in the distribution of SNPs with significant CMH p values and the genome-wide increase in $F_{ST}$ over time, a large proportion of peaks are found on chrIV across all lake combinations, though other chromosomes appear to also be almost equally prominent when only looking at Loberg, suggesting that chrIV is driving the earliest stages of freshwater adaptation as reflected by Cheney and Scout, but that other chromosomes "catch up" later on (**Fig S21.11 - Fig S21.15**).

Using genetic distance always results in fewer peaks than the equivalent test with physical distance, by ~20-40% in the case of sensitive TempoPeaks and ~60% for specific TempoPeaks (**Table S21.1**). It appears that some regions of the genome that appear as multiple distinct peaks in physical distance are likely single peaks when taking into account variation in the recombination rate, which is also reflected in the total size of TempoPeaks being of similar magnitude using genetic versus physical distance (2.8% versus 3.8% of the genome for CH+SC+LB for specific TempoPeaks **Table S21.2**) and with the genetic distance TempoPeaks generally being large despite the reduced number (**Table S21.3**). TempoPeak sizes vary considerably. When using genetic distance and looking at CH+SC+LB specific TempoPeaks using genetic distance, the median TempoPeak size is ~31kb, but with a 5th and 95% percentile that ranges from ~2.5 to 275kb, and the largest peak being ~1.5Mb long (chrIV: 20862645-22379208).

Almost all specific TempoPeaks identified in the contemporary evolution samples overlap with sensitive EcoPeaks in the geographic analysis (**Table S21.4**), strongly suggesting that these regions reflect true positives that exhibit consistent recurrent and rapid evolution in threespine stickleback. There is much less overlap for the sensitive TempoPeaks identified in CH+SC+LB and LB (on the order of 40%), presumably either due to the evolution of specific local peaks or false positives, though there is a noticeable improvement when looking at CH and SC. The presence of considerable false positives amongst the sensitive sets is also likely reflected in the total amount of the genome found in significant TempoPeaks, where for both CH+SC+LB and LB ~ 20% of the genome is found in sensitive TempoPeaks, while the this value is <3% in the case of specific TempoPeaks (**Table S21.2**)

When examining specific TempoPeaks defined by genetic distance (the most conservative conditions), we found 12 TempoPeaks in CH+SC that were absent in corresponding specific LB analysis. These could be rapidly evolving loci that were not selected for during the accidental colonization of Loberg, perhaps because they were absent in the initial founders (**Fig S21.16**). However, we also note that these are present in the LB sensitive TempoPeak set, and so may reflect limited statistical power due to slower evolution.

**Table S21.1.** Number of sensitive and specific TempoPeaks

| # of peaks | Sensitive (physical) | Sensitive (genetic) | Specific (physical) | Specific (genetic) |
|---|---|---|---|---|
| CH+SC+LB | 524 | 392 | 344 | 142 |
| LB | 589 | 467 | 246 | 99 |
| CH+SC | 271 | 159 | 86 | 30 |
| CH | 193 | 119 | 9 | 4 |
| SC | 102 | 60 | 0 | 0 |

**Table S21.2.** Total genomic size of sensitive and specific TempoPeaks

| # of peaks | Sensitive (physical) | Sensitive (genetic) | Specific (physical) | Specific (genetic) |
|---|---|---|---|---|
| CH+SC+LB | 97,975,057 | 129,580,111 | 17,568,150 | 13,038,873 |
| LB | 85,857,279 | 121,512,167 | 11,224,050 | 6,843,058 |
| CH+SC | 53,262,302 | 76,011,011 | 3,268,900 | 1,527,516 |
| CH | 30,676,157 | 53,913,015 | 313,850 | 136,680 |
| SC | 22,319,671 | 40,373,872 | 0 | 0 |

**Table S21.3.** Attributes of CH+SC+LB TempoPeak sizes

| Peak sizes (bp) | Sensitive (physical) | Sensitive (genetic) | Specific (physical) | Specific (genetic) |
|---|---|---|---|---|
| mean | 186,723 | 331,439 | 53,570 | 93,323 |
| median | 50,926 | 41,970 | 29,425 | 31,980 |
| 5th percentile | 1 | 1 | 3,208 | 2,544 |
| 95th percentile | 851,923 | 1,633,693 | 163,955 | 275,830 |

**Table S21.4.** Percentage of overlap between TempoPeaks and sensitive EcoPeaks.

| % overlap | Sensitive (physical) | Sensitive (genetic) | Specific (physical) | Specific (genetic) |
|---|---|---|---|---|
| CH+SC+LB | 38 | 34 | 94 | 95 |

| | | | | |
|---|---|---|---|---|
| LB | 38 | 31 | 93 | 95 |
| CH+SC | 73 | 70 | 99 | 100 |
| CH | 76 | 69 | 100 | 100 |
| SC | 85 | 86 | NA | NA |

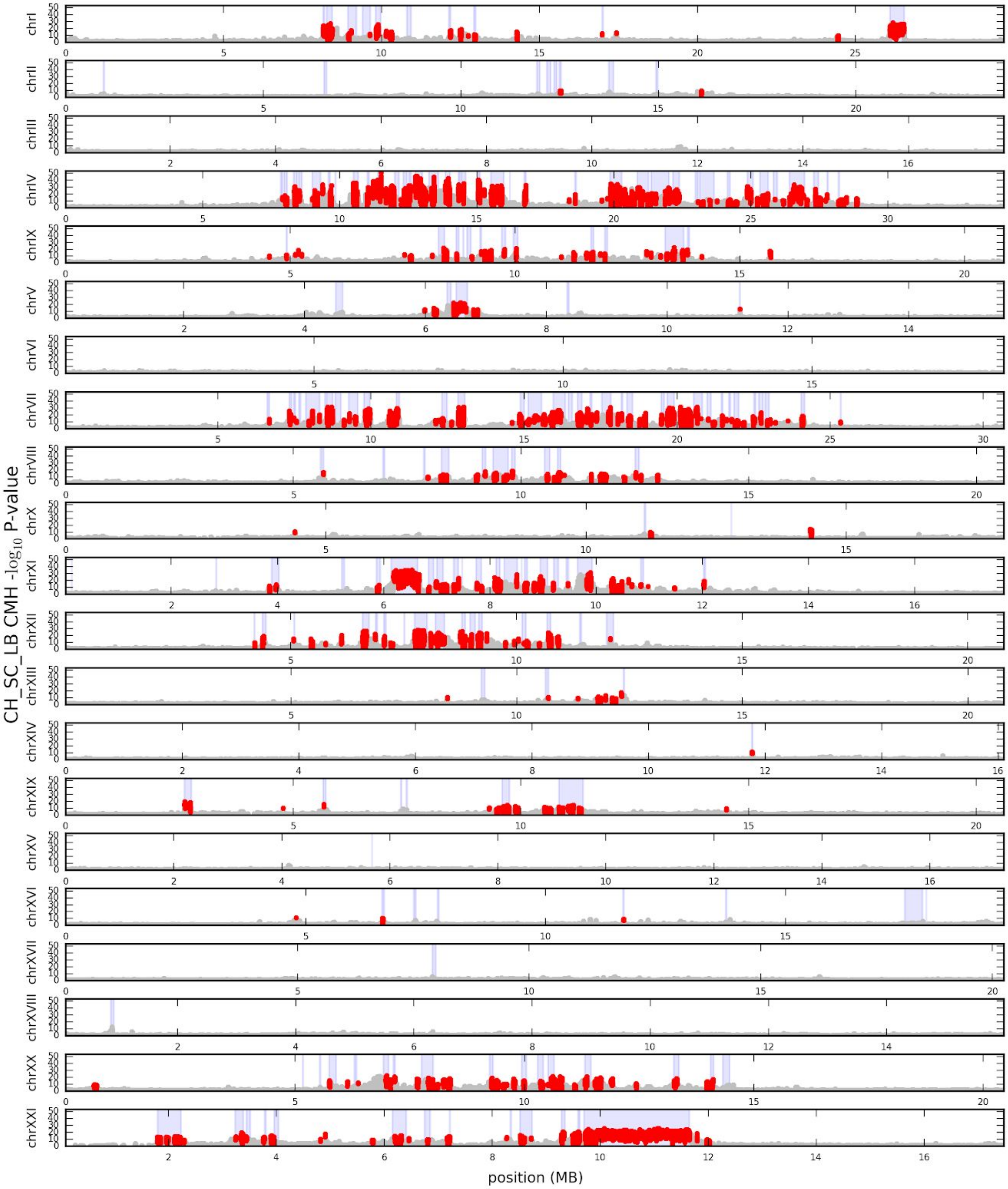


**Fig_S21.1:** Genome-wide distribution of mean CMH -log10 p-values in windows of 2500bp sliding every 50bp for each chromosome for CH+SC+LB, with position in physical distance. Red regions represent windows in significant specific TempoPeaks. Blue vertical regions are specific EcoPeaks

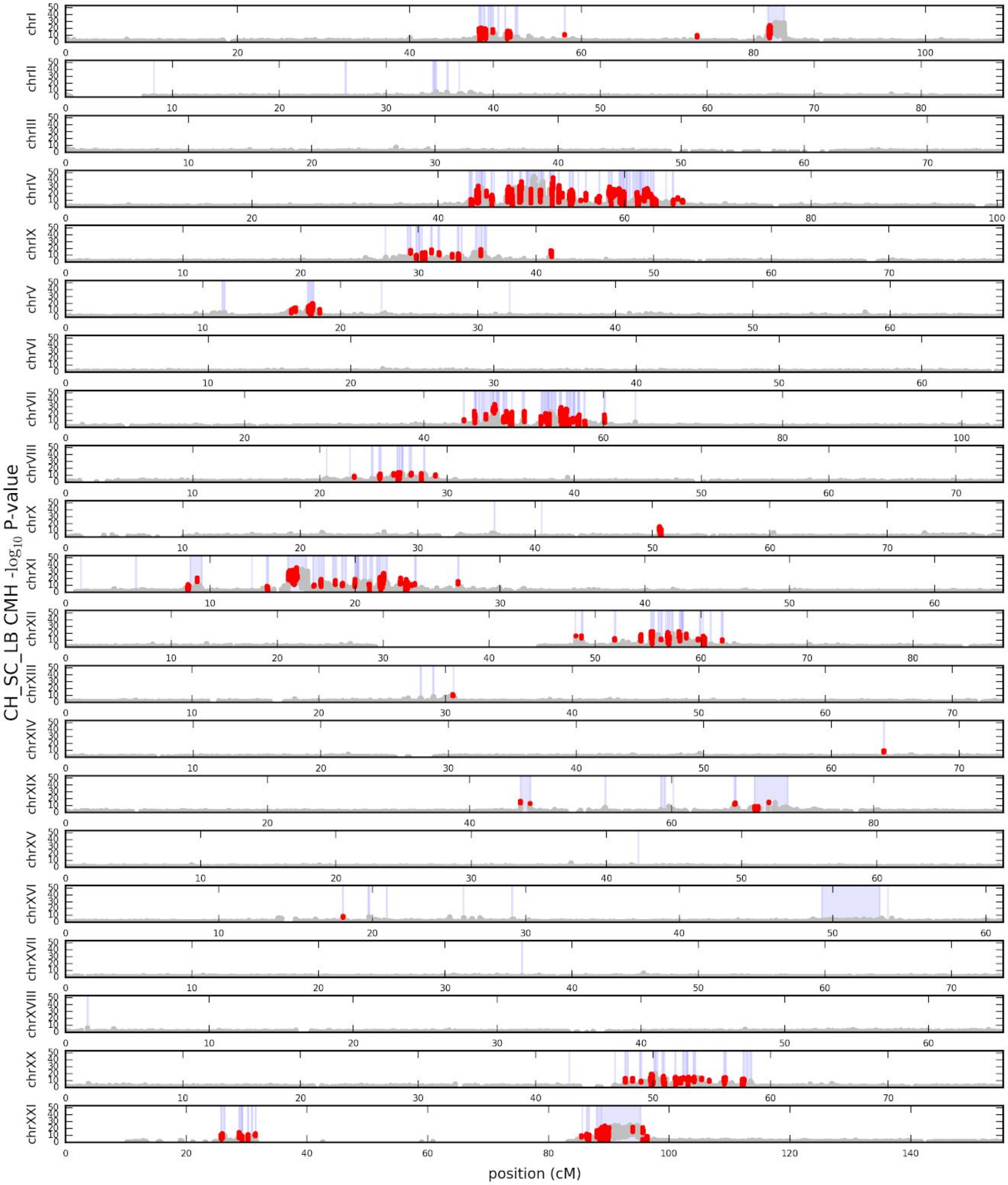


**Fig_S21.2:** Genome-wide distribution of mean CMH -log10 p-values in windows of 0.01cM sliding every 0.0002cM for each chromosome for CH+SC+LB, with position in genetic distance. Red regions represent windows in significant specific TempoPeaks. Blue vertical regions are specific EcoPeaks

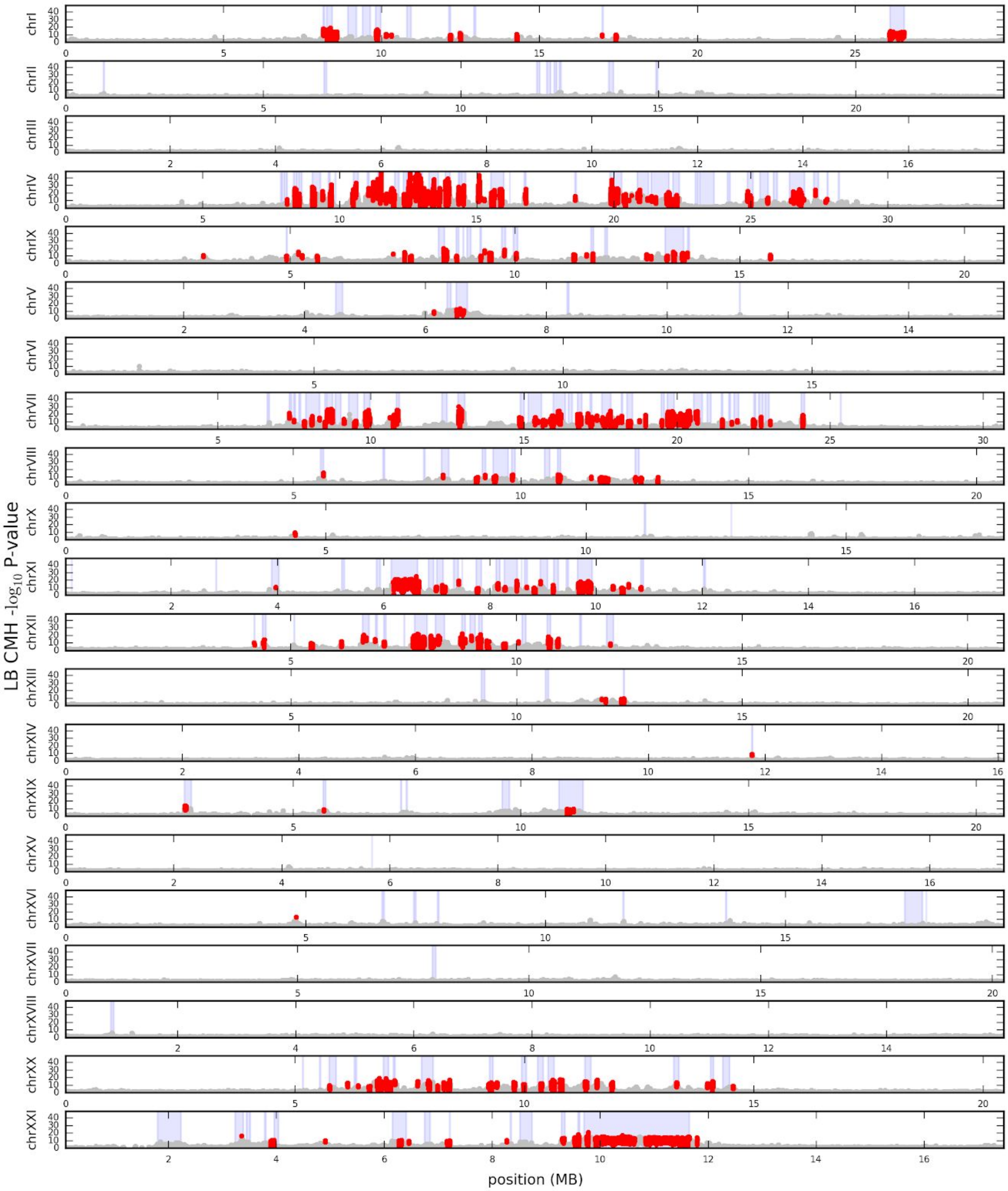


**Fig_S21.3:** Genome-wide distribution of mean CMH -log10 p-values in windows of 2500bp sliding every 50bp for each chromosome for LB, with position in physical distance. Red regions represent windows in significant specific TempoPeaks. Blue vertical regions are specific EcoPeaks

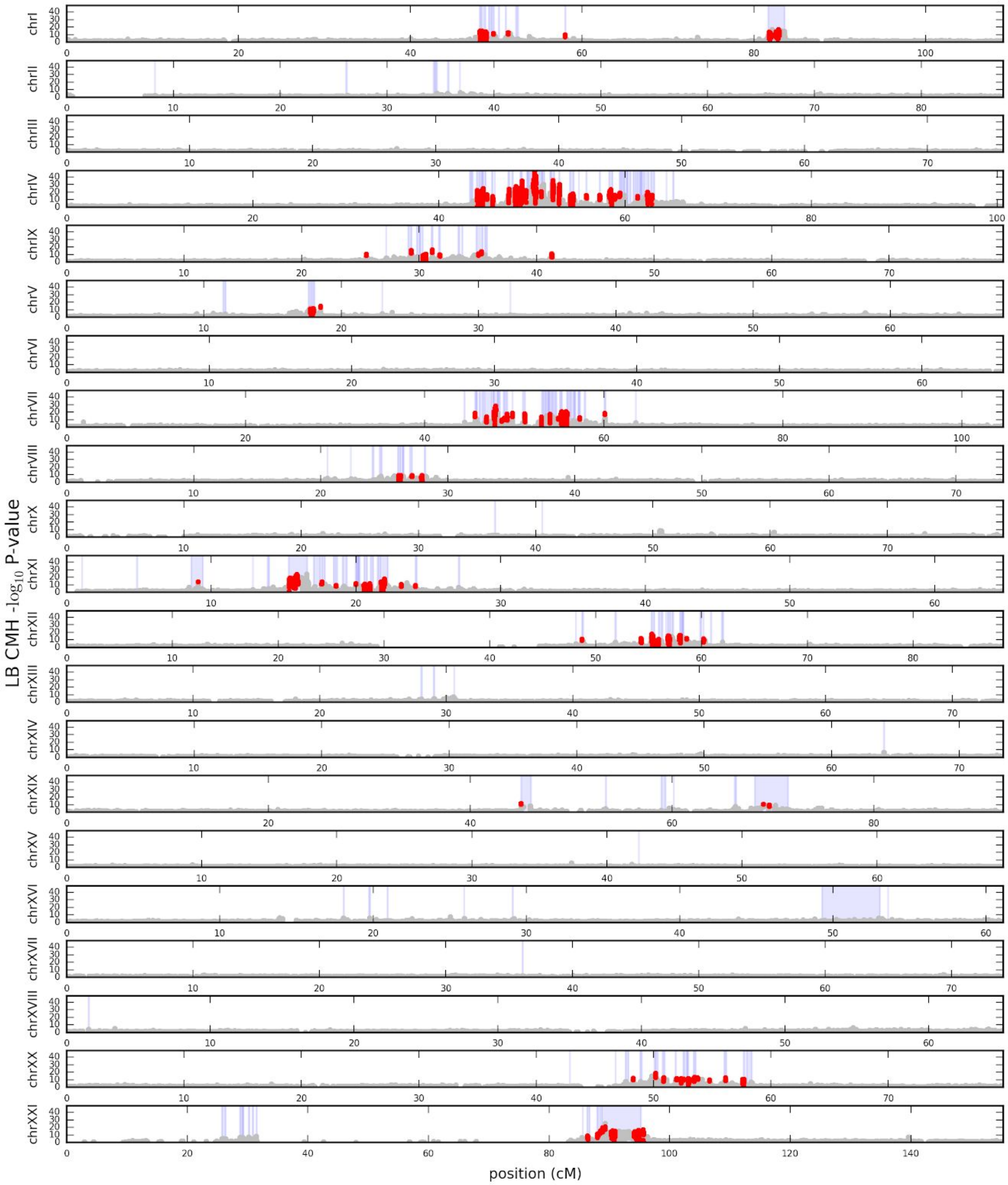


**Fig_S21.4:** Genome-wide distribution of mean CMH -log10 p-values in windows of 0.01cM sliding every 0.0002cM for each chromosome for LB, with position in genetic distance. Red regions represent windows in significant specific TempoPeaks. Blue vertical regions are specific EcoPeaks

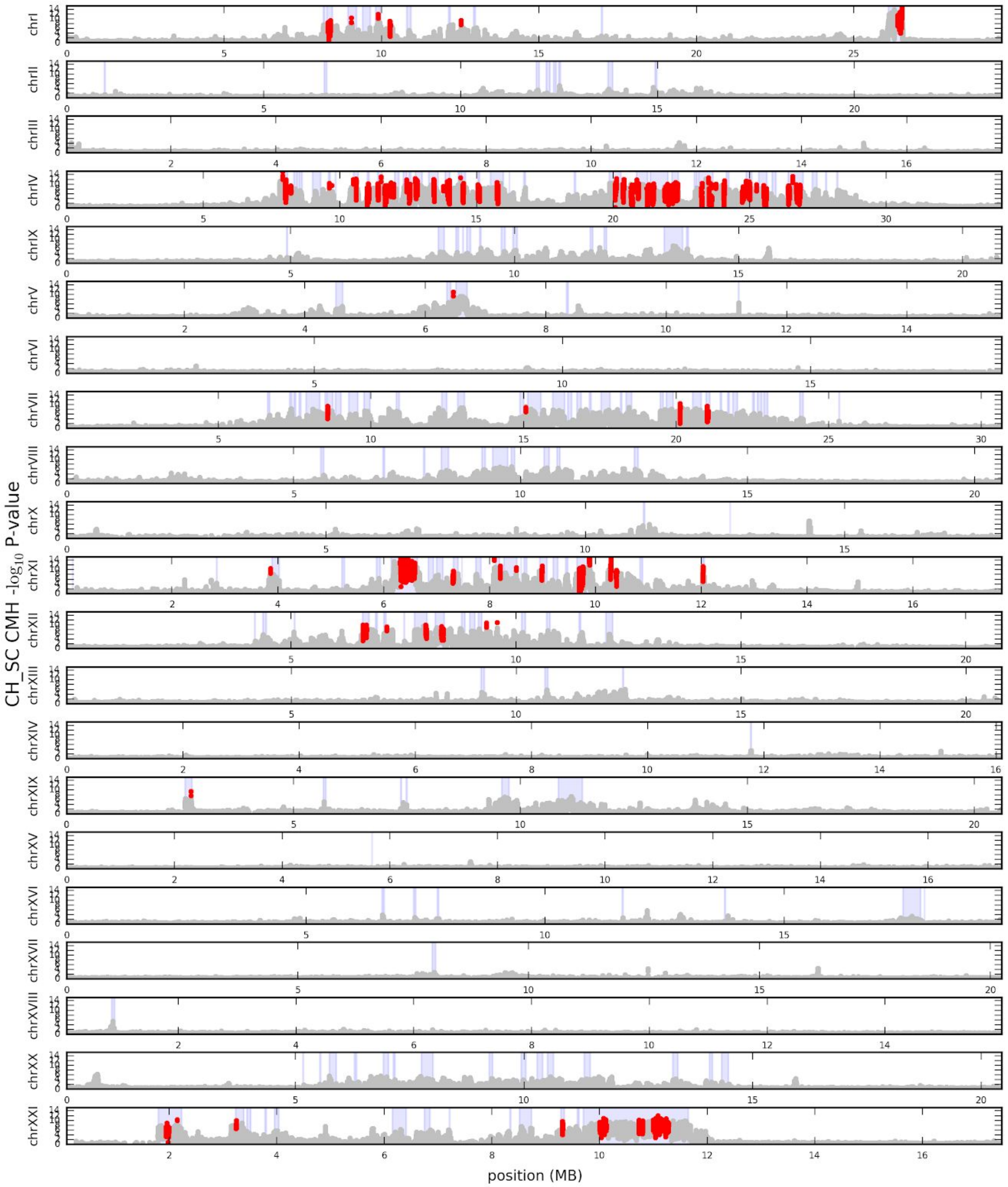


**Fig_S21.5:** Genome-wide distribution of mean CMH -log10 p-values in windows of 2500bp sliding every 50bp for each chromosome for CH+SC, with position in physical distance. Red regions represent windows in significant specific TempoPeaks. Blue vertical regions are specific EcoPeaks

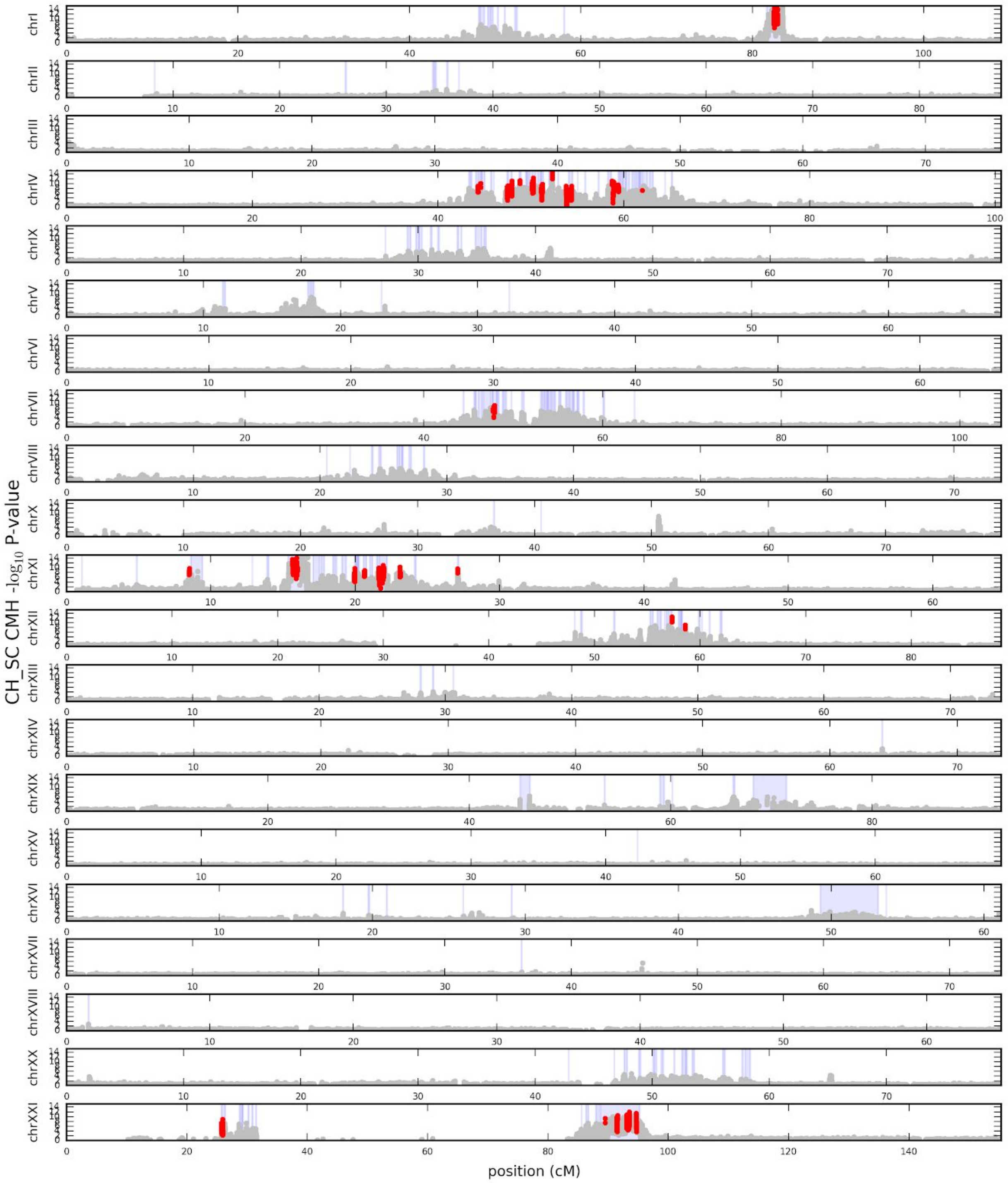


**Fig_S21.6:** Genome-wide distribution of mean CMH -log10 p-values in windows of 0.01cM sliding every 0.0002cM for each chromosome for CH+SC, with position in genetic distance. Red regions represent windows in significant specific TempoPeaks. Blue vertical regions are specific EcoPeaks

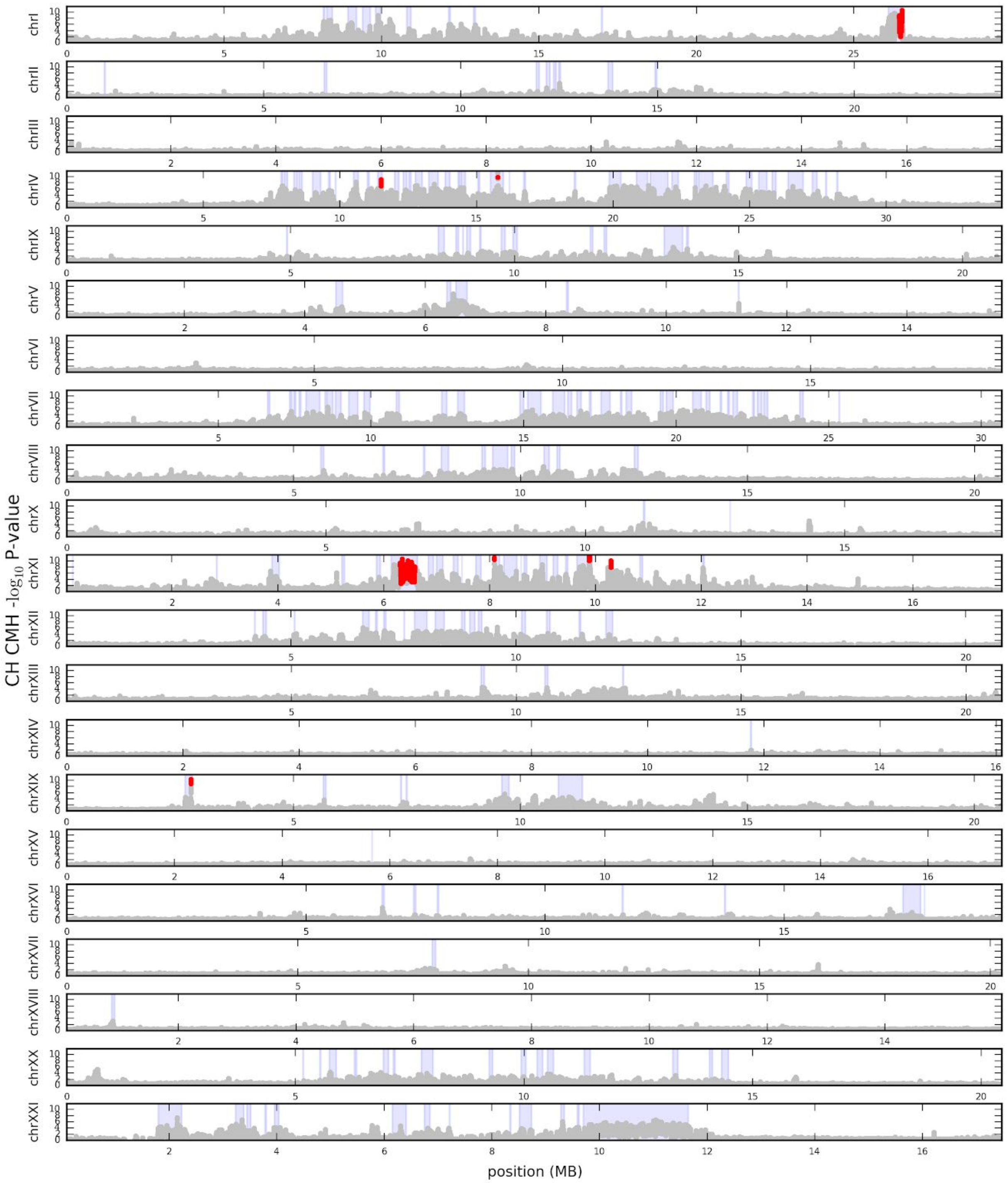


**Fig_S21.7:** Genome-wide distribution of mean CMH -log10 p-values in windows of 2500bp sliding every 50bp for each chromosome for CH, with position in physical distance. Red regions represent windows in significant specific TempoPeaks. Blue vertical regions are specific EcoPeaks

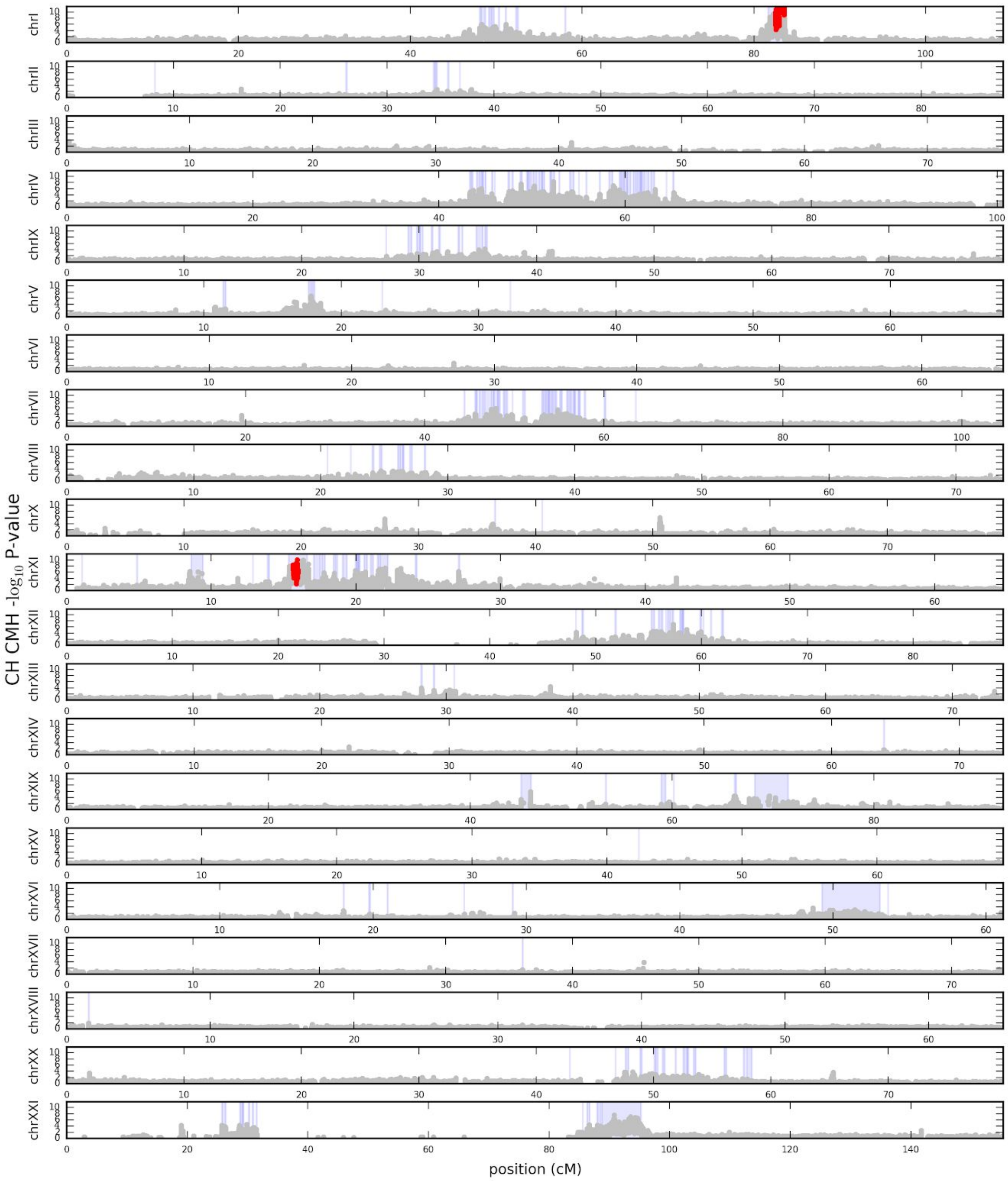

**Fig_S21.8:** Genome-wide distribution of mean CMH -log10 p-values in windows of 0.01cM sliding every 0.0002cM for each chromosome for CH, with position in genetic distance. Red regions represent windows in significant specific TempoPeaks. Blue vertical regions are specific EcoPeaks

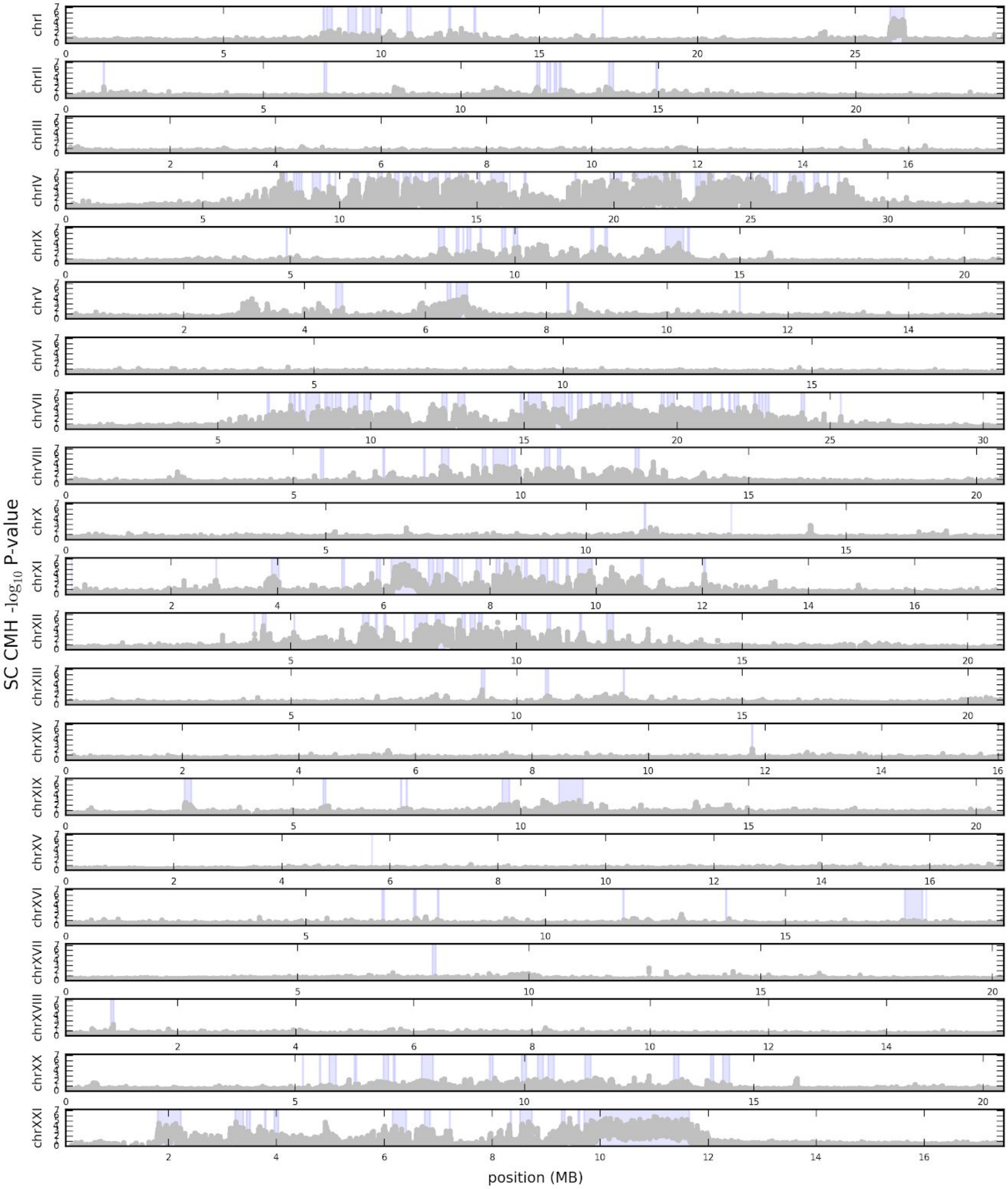


**Fig_S21.9:** Genome-wide distribution of mean CMH -log10 p-values in windows of 2500bp sliding every 50bp for each chromosome for SC, with position in physical distance. Red regions represent windows in significant specific TempoPeaks. Blue vertical regions are specific EcoPeaks

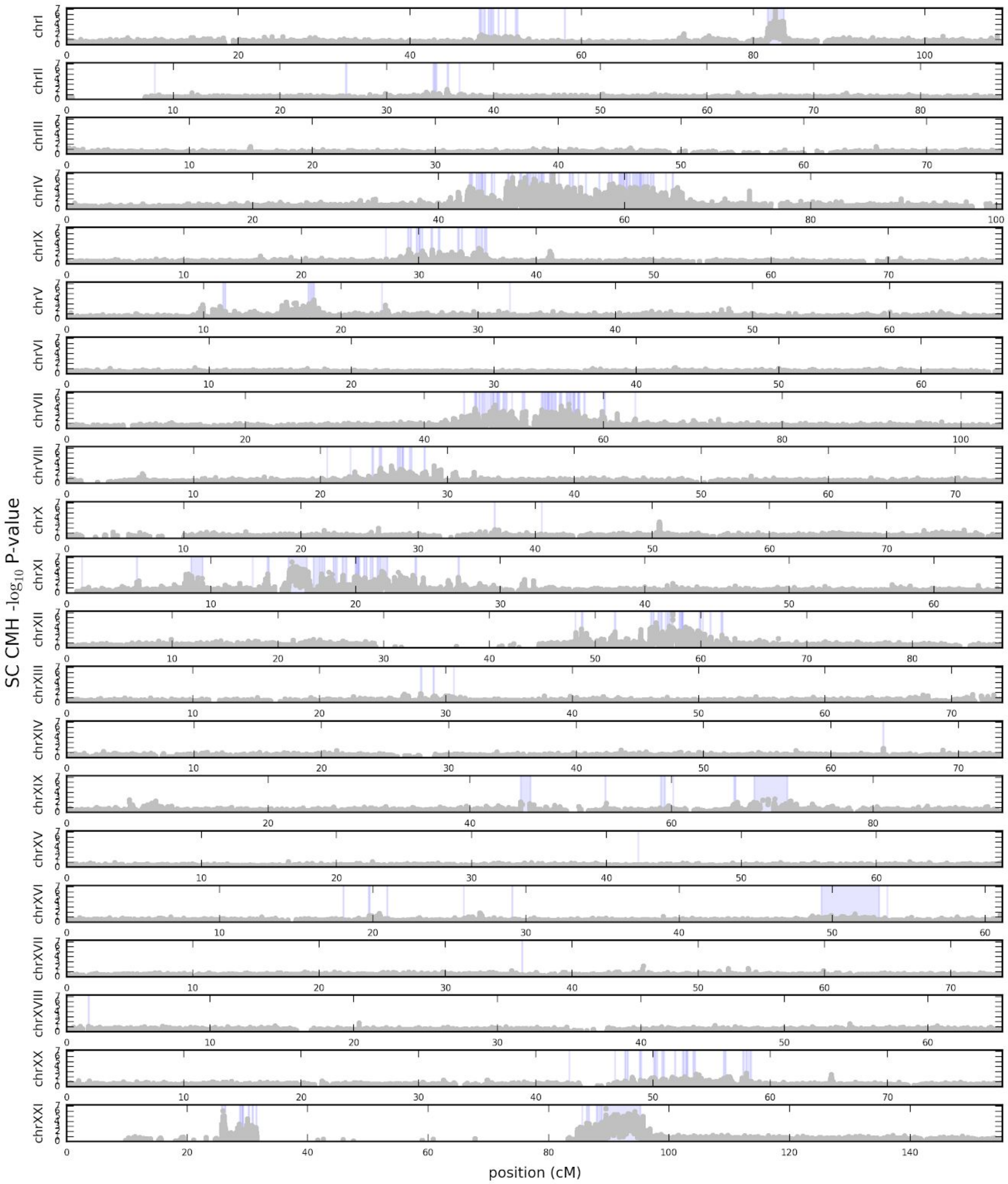


**Fig_S21.10:** Genome-wide distribution of mean CMH -log10 p-values in windows of 0.01cM sliding every 0.0002cM for each chromosome for SC, with position in genetic distance. Red regions represent windows in significant specific TempoPeaks. Blue vertical regions are specific EcoPeaks

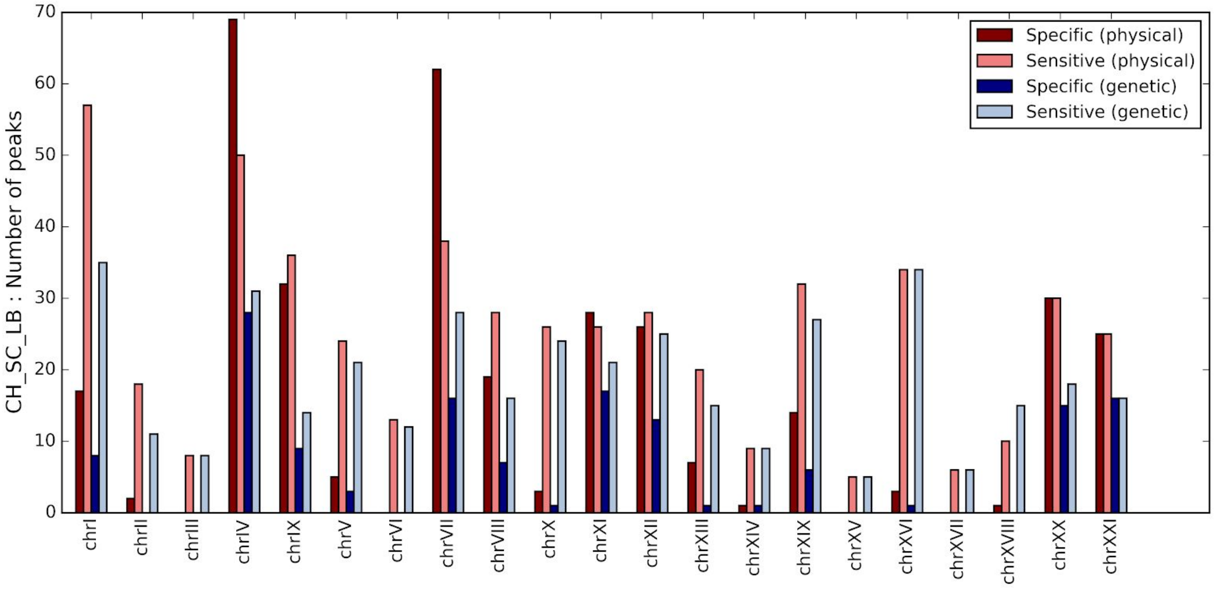


**Fig_S21.11:** Number of significant specific and sensitive TempoPeaks based on physical and genetic distance for CH+SC+LB.

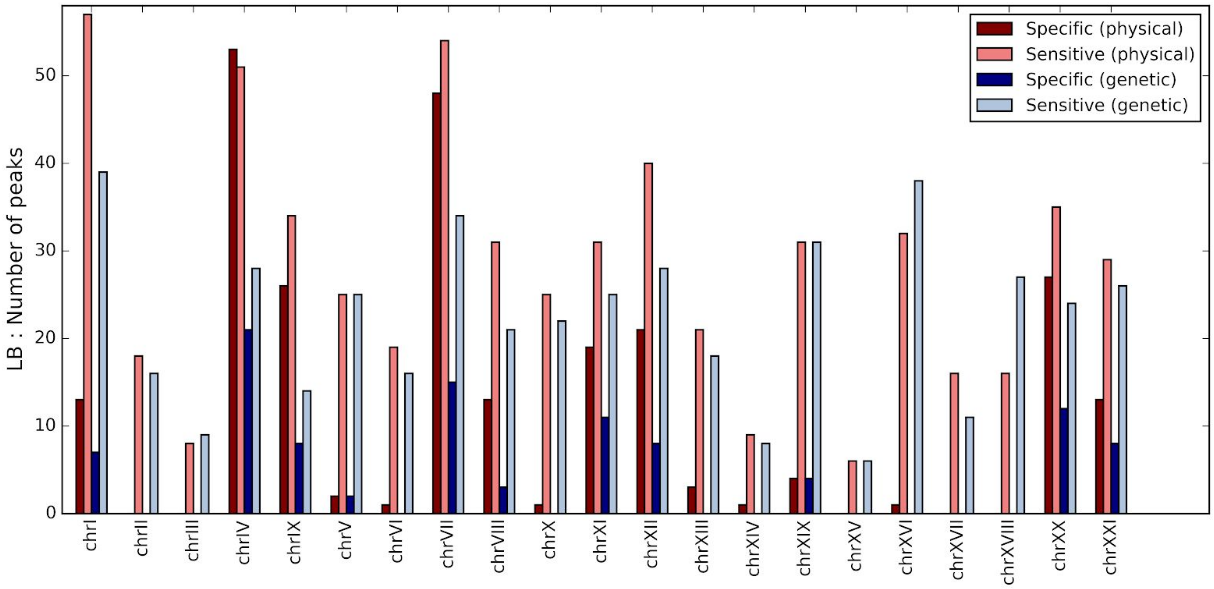


**Fig_S21.12:** Number of significant specific and sensitive TempoPeaks based on physical and genetic distance for LB.

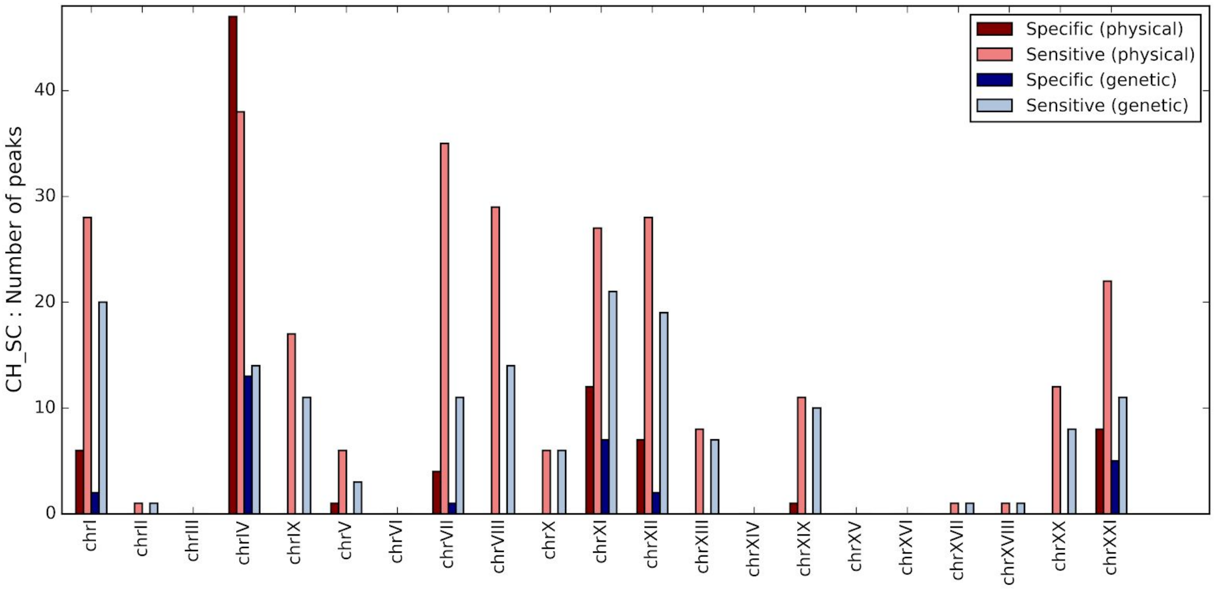


**Fig_S21.13:** Number of significant specific and sensitive TempoPeaks based on physical and genetic distance for CH+SC.

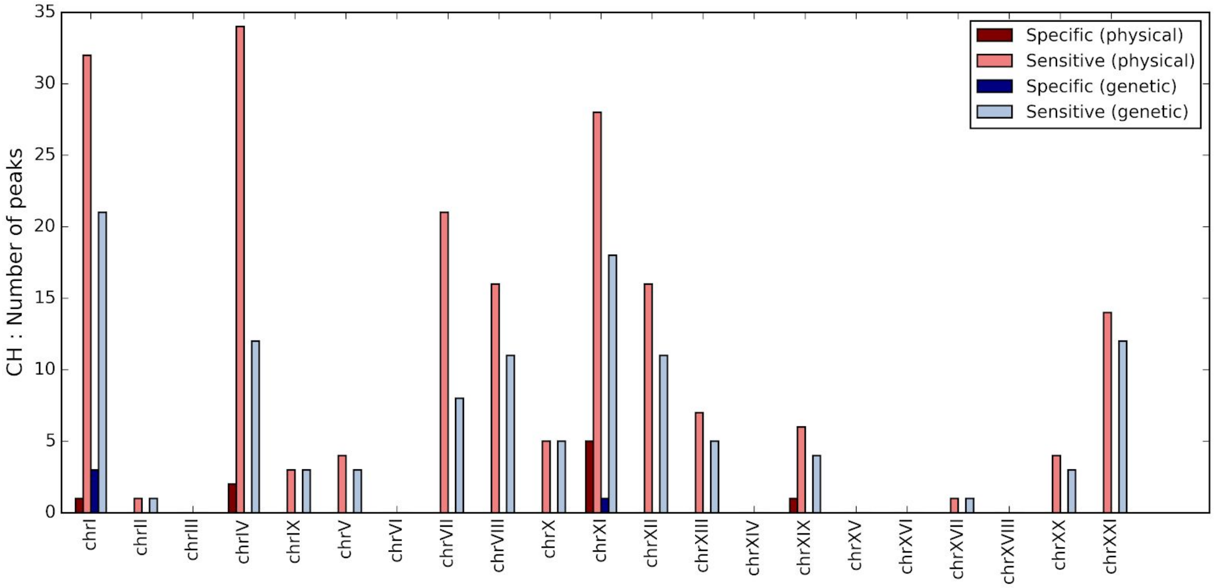


**Fig_S21.14:** Number of significant specific and sensitive TempoPeaks based on physical and genetic distance for CH

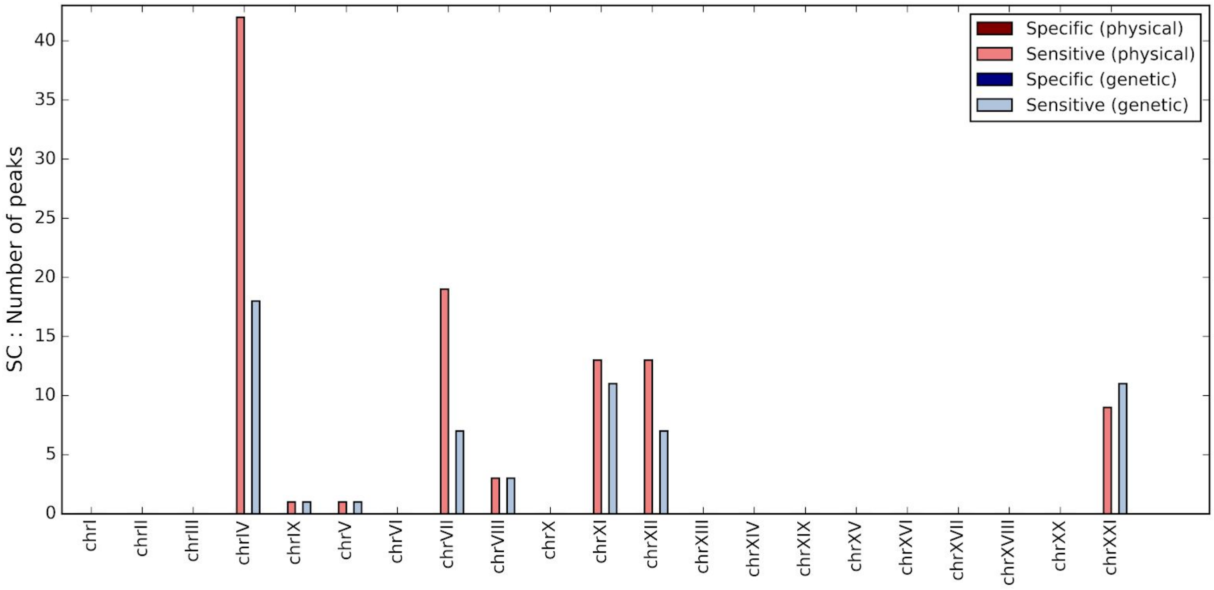


**Fig_S21.15:** Number of significant specific and sensitive TempoPeaks based on physical and genetic distance for SC.

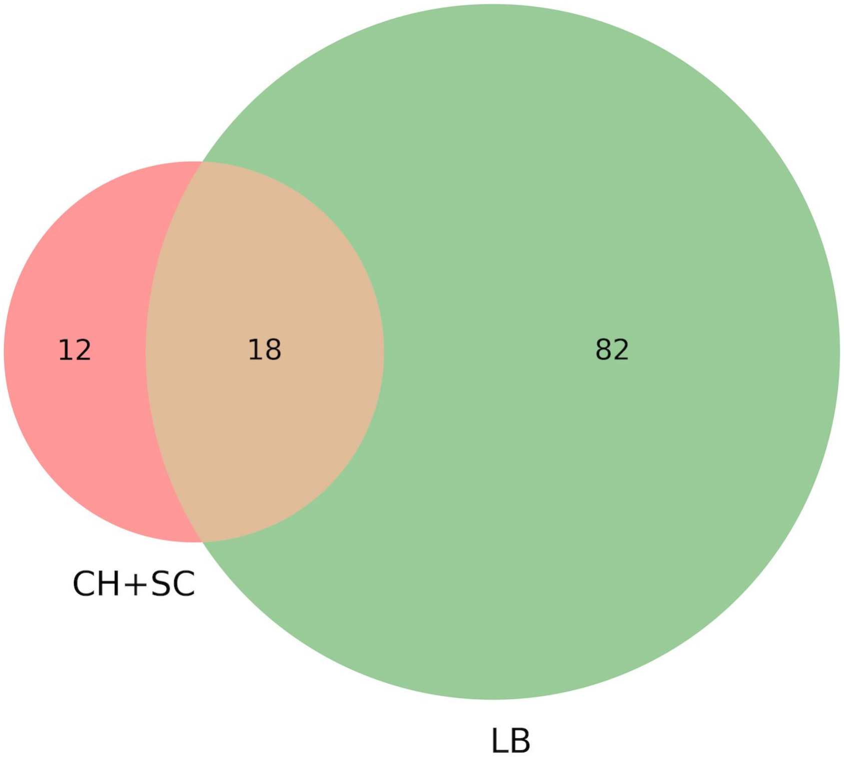


**Fig S21.16.** Venn diagram showing number of private and shared significant peaks when defined using our specific p-value criteria and genetic distance

#### 22. COMPARING TEMPOPEAKS AND RECOMBINATION HOTSPOTS

When plotting our specific window-based TempoPeaks based on genetic distance alongside recombination hotspots defined as being >20× the background rate, we observed notable overlap between clusters of TempoPeaks on individual chromosomes and hotspots (**Fig S22.1**). This overlap was particularly apparent on chrIV. Looking closer at the major clusters on this chromosome, we observe that hotspots often lie in between TempoPeaks (**Fig S22.2** and **Fig S22.3**). Thus, while clusters of TempoPeaks generally lie in low recombination regions, there appear to be hotspots between these TempoPeaks, perhaps to allow rapid assembly of selected haplotypes and avoid Hill-Robertson interference (*62*).

We formally tested this relationship for pairs of TempoPeaks and hotspots by identifying all pairs of successive TempoPeaks on the same chromosome (as defined by the CH+SC+LB specific criteria using genetic distance) separated by a maximum distance of 4cM (equivalent to ~1Mb) as measured from the closest boundaries of each TempoPeak, and noting how many such pairs possessed a recombination hotspot within the interval. We then performed a bootstrap procedure to test how unusual this result was by randomly choosing pairs of genomic positions matching the genetic distance between the real set of TempoPeaks (the only condition being that for each pair of real TempoPeaks, the random positions were chosen from the same chromosome) and noting how often hotspots were found in the intervals of these random pairs. We repeated this process 10,000 times and generated an empirical p-value by comparing the number of TempoPeaks with hotspots in the intervals for the real data to the randomly chosen positions.

From the real data there were 120 successive TempoPeaks separated by a maximum distance of 4cM, 83 of which we found a recombination hotspot within the interval. Amongst our 10,000 randomly selected sets of matching peaks, the maximum number of randomly chosen position pairs with hotspots in between was 37 with a mean of 21.7, suggesting that our result was highly non-random ($p<0.0001$). Using physical distance instead with a maximum distance between loci of 1Mb produces a similar extreme result that cannot be replicated with randomly chosen pairs of loci ($p<0.0001$), with 60/94 pairs of observed TempoPeaks possessing a recombination hotspot in the interval, versus a maximum of 59 in our randomly chosen pairs and a mean of 39.7.

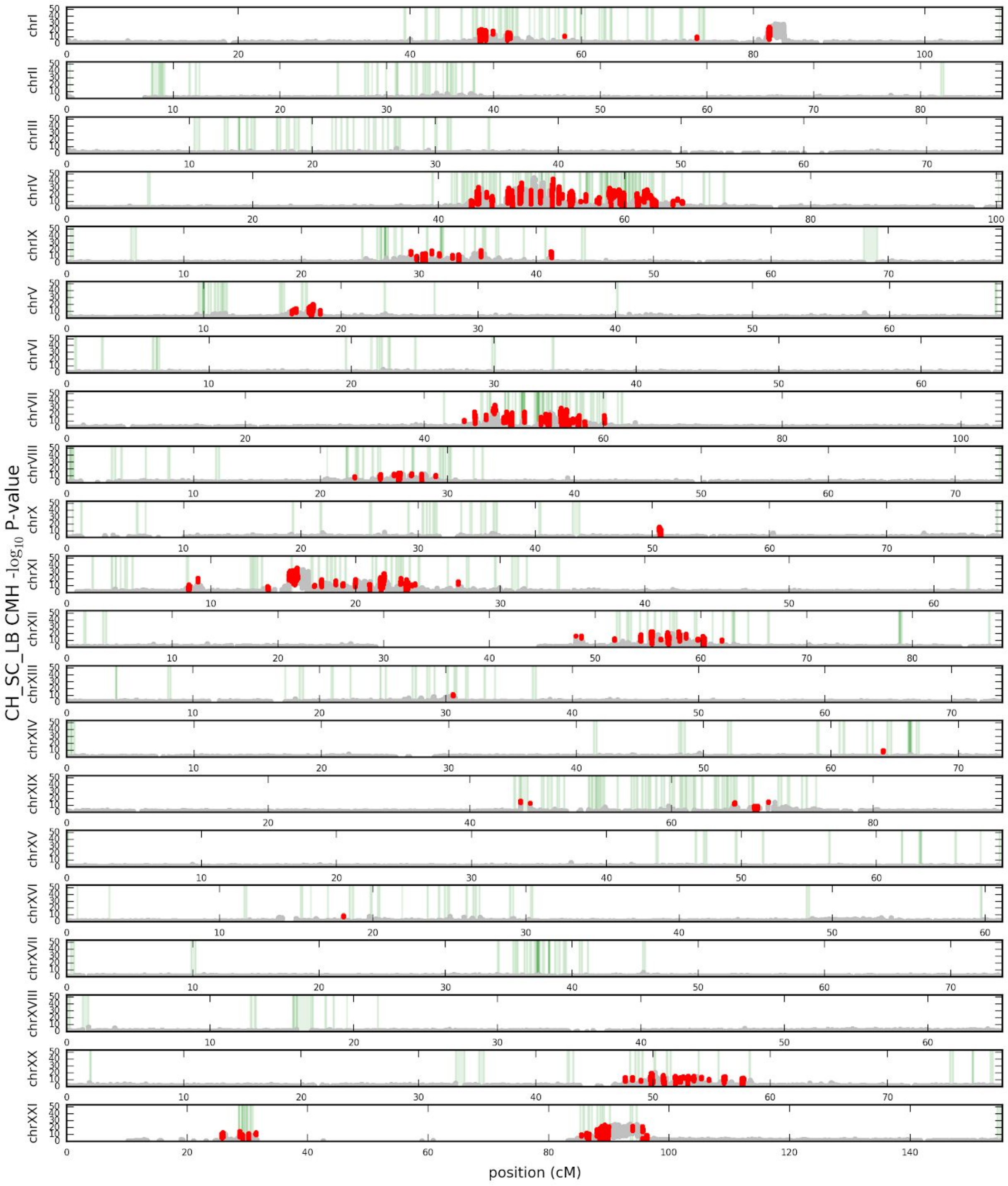


**Fig S22.1:** Genome-wide distribution of mean CMH -log10 p-values in windows equivalent to 2500bp sliding every 50bp for each chromosome for CH+SC+LB, with position in genetic distance. Red regions represent windows in specific TempoPeaks. Green vertical regions are recombination hotspots that are defined as 20× the background rate.

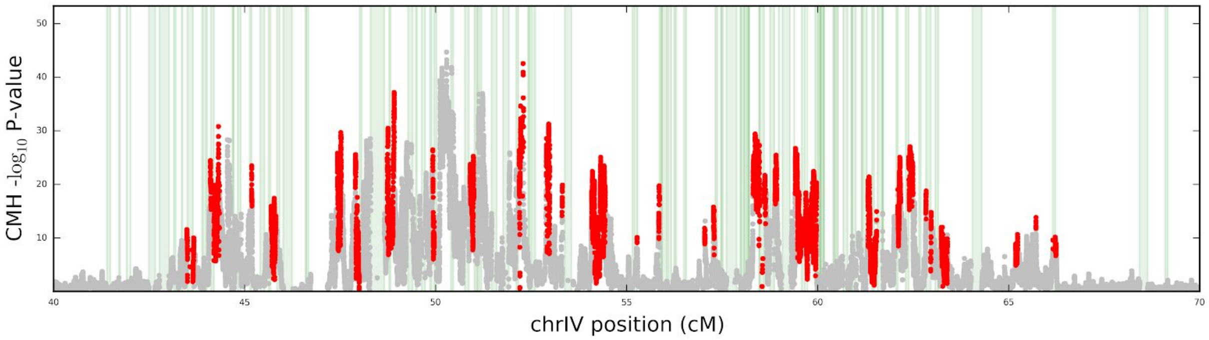


**Fig S22.2:** Distribution of mean CMH -log10 p-values in windows equivalent to 2500bp sliding every 50bp for each chromosome for CH+SC+LB, with position in genetic distance, focusing in on chrIV from 40 to 70cM. Red regions represent windows in specific TempoPeaks. Green vertical regions are recombination hotspots that are defined as 20× the background rate.

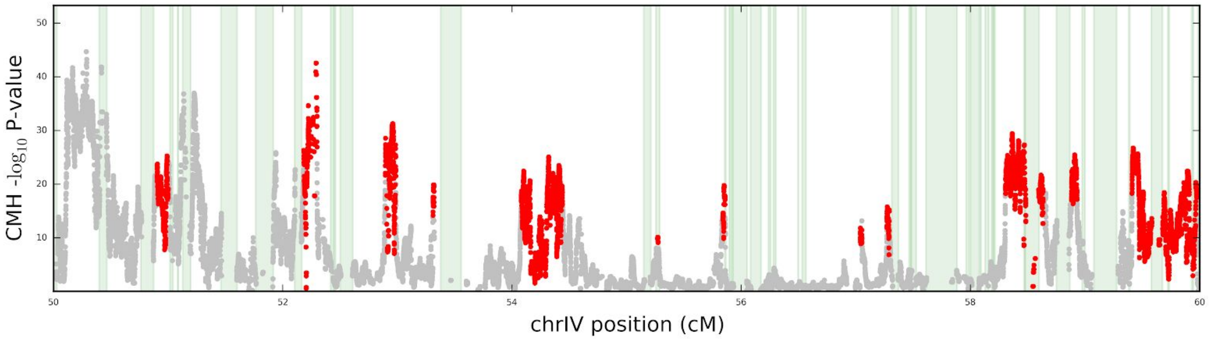


**Fig S22.3:** Distribution of mean CMH -log10 p-values in windows equivalent to 2500bp sliding every 50bp for each chromosome for CH+SC+LB, with position in genetic distance, focusing in on chrIV from 50 to 60cM. Red regions represent windows in specific TempoPeaks. Green vertical regions are recombination hotspots that are defined as 20× the background rate.

### 23. ESTIMATING TEMPOPEAK SELECTION COEFFICIENTS ASSUMING SNP INDEPENDENCE

Our replicate time series data provided the possibility of estimating selection coefficients ($s$) for individual SNPs by modelling allele frequency trajectories over time. We applied the method described in Taus et al. (*63*) to estimate $s$ for the SNP with the largest CMH score in each significant TempoPeak. This method implements a deterministic approach to estimate $s$, which we consider justified for our data due to the extremely rapid allele frequency increases involved (implying large $s$ values) and that selection is occurring on standing genetic variation that is already at some notable frequency in the starting populations ($p0$). We estimate $s$ and $p0$ both assuming the dominance coefficient, *h*, is 0.5, as well as estimating *s*, $p0$ and *h* simultaneously. For the former we perform a linear least-squares regression (LLS) (performed in Python using `scipy.stats.linregress`) to the logit transformed allele frequencies for each time point relative to the number of generations (assuming one 1 year per generation for stickleback) since the founding of the population (i.e. the colonization of Cheney, Scout and Loberg). The slope and y-intercept then provide estimates of $s$ and $p0$, respectively. We do not perform the bias correction described by Taus et al.(*63*), instead simply noting that our estimates may be a slight underestimate of the true value under this specific model, which as we will see below in the section on Deep Neural Networks, is not a very likely one for our data. When estimating $h$ alongside $s$ and $p0$, we performed a nonlinear least squares (NLS) optimization (performed in Python using `scipy.optimize.curve_fit`) of allele frequencies relative to the number of generations. The model function for the NLS was constructed based on iterating forward the well known expression for change in allele frequency forward in time for one generation assuming infinite population size (*64*) :

$$\Delta p = \frac{pq[p(w_{11}-w_{12})+(q(w_{12}-w_{22})]}{p^2 w_{11}+2pqw_{12}+q^2 w_{22}}$$

where $w_{11} = 1$, $w_{12} = 1\text{-}s(1\text{-}h)$ and $w_{22}$=1-$s$. Usually implementing such a function would be computationally difficult when considering thousands of generations, but the number of total generations required to iterate forward in our case is small (no more than 30 generations).

We applied these methods to all specific and sensitive TempoPeaks for all 5 subsets of lakes and for both physical and genetic distance. We utilized sample allele frequencies previously estimated by the method of Lynch et al.(*24*) for each given SNP as a proxy for population allele frequencies. Given the large $s$ values estimated here, we expect this to not to largely affect our analysis, and Taus et al.(*63*) demonstrated that coverage likely has little effect on estimation.

When examining only significant CH+SC peaks, allele frequencies for the most significant SNP within each peak often rose from close to 0 to above 50% within 8 generations, an extremely rapid rate of change for a vertebrate species. Thus $s$ estimates were generally very high, ranging from 0.36-0.76 with a mean 0.54 for the 74 CH+SC specific TempoPeaks determined using physical distance (**Fig S23.1**, **Table S23.1**) and 0.44-0.70 with a mean of 0.56 for the 30 specific TempoPeaks determined using genetic distance (**Fig S23.2**, **Table S23.2**). Examining the 344 and 142 CH+SC+LB specific TempoPeaks (thus extending the fit to ~30 generations since founding) identified under the same physical and genetic distance criteria respectively resulted in a wider range (0-0.64 for physical distance, 0.08-0.68 for genetic) but still wide a very high mean of 0.3 and 0.33 (**Fig S23.3, Fig S23.4**, **Table S23.3, Table S23.4**). Visually, using *h*=0.5 under the LLS generally provided a poor fit to the observed data, especially when considering all three populations, with this model usually being unable to capture the earliest stages of freshwater founding where allele frequencies increase much faster than possible under semi-dominance. For the vast majority of loci, a $h$ value close to 1 provided the best fit of the observed data.

In the previous section we identified 12 specific TempoPeaks that were significant when analyzing CH+SC but not LB when analyzing both under specific criteria. The CH+SC peaks did overlap LB sensitive peaks as well as long-term peaks suggesting that they may still be under selection in Loberg despite their absence amongst the specific set of peaks, though this must be balanced by our sensitive set likely being overly aggressive (~20% of the genome is considered to experience a significant allele frequency change). When estimating *s* values from the CH+SC allele frequency trajectories, *s* ranged from 0.44 to 0.69 with a mean of 0.53, and thus allele frequencies in CH+SC are indeed increasing rapidly at these 12 loci. Examination of allele frequency trajectories of the same SNPs in Loberg at these peaks showed a mixed pattern (**Fig S23.5**). Some SNPs demonstrated rapidly increasing frequencies in CH+SC as well as high frequencies in Loberg. Further inspection of these loci showed that the lack of significant peak in Loberg could have resulted from from the specific Loberg peak being slightly shifted upstream or downstream of the CH+SC (and thus they were essentially part of the same long-term block being selected for, but perhaps with slightly different haplotype configurations) (e.g. **Fig S23.6**). Alternatively this could have been an artifact of our peak calling criteria, where peaks needed to be 50kb (0.2cM) from the boundary of another more significant peak (something that happens on chrIV in particularly due to the large amount of that chromosome that is under selection and so CMH peaks are so tightly clustered together).

However, there are a set of 4 SNPs that show rapidly increasing allele frequencies in CH+SC and either low or moderately increasing allele frequencies in LB (chrIV:15,433,790, chrXI:3,860,620, chrXI:8,098,124, chrXI:12,033,114). ChrXI:12,033,114 provides the cleanest signal of a CH+SC peak that is evolving less rapidly in LB (**Fig S23.7**). In all four cases, there is some evidence that the CMH scores in these genomic regions in Loberg are greater than the background (and hence they all lie in the sensitive set of peaks), but the allele frequency changes since founding 30 years ago are not nearly as rapid as the frequency changes that have occurred in Cheney and Scout in 8 years and less.

**https://drive.google.com/file/d/1LnocIJfNans0dylZfbcgkFGDMiV7YS57/view?usp=sharing**
**Fig_S23.1**. GIF showing for each significant specific peak determined using physical distance the observed allele frequencies over time for CH+SC alongside best fit *s* and *p0* trajectories assuming *h*=0.5 (s_LLS, dashed purple line) and when estimating *h* (s_NLS, purple line). GIF made using https://imgflip.com/gif-maker?from=images

**https://drive.google.com/file/d/1TCA5C3ANtEgLHNTiP4OTxbgRh7oPU2F0/view?usp=sharing**
**Fig_S23.2**. GIF showing for each significant specific peak determined using genetic distance the observed allele frequencies over time for CH+SC alongside best fit *s* and *p0* trajectories assuming *h*=0.5 (s_LLS, dashed purple line) and when estimating *h* (s_NLS, purple line). GIF made using https://imgflip.com/gif-maker?from=images

**https://drive.google.com/file/d/11nGyGO5WbEKgFAb08LxQXQ8ky1RgLQM6/view?usp=sharing**
**Fig_S23.3**. GIF showing for each significant specific peak determined using physical distance the observed allele frequencies over time for CH+SC+LB alongside best fit *s* and *p0* trajectories assuming *h*=0.5 (s_LLS, dashed purple line) and when estimating *h* (s_NLS, purple line).GIF made using https://imgflip.com/gif-maker?from=images

**https://drive.google.com/file/d/1jV5VZ0JiUknaVNeTcg6gWM52Z1pOc3Ty/view?usp=sharing**
**Fig_S23.4**. GIF showing for each significant specific peak determined using genetic distance the observed allele frequencies over time for CH+SC+LB alongside best fit *s* and *p0* trajectories assuming *h*=0.5 (s_LLS, dashed purple line) and when estimating *h* (s_NLS, purple line).GIF made using https://imgflip.com/gif-maker?from=images

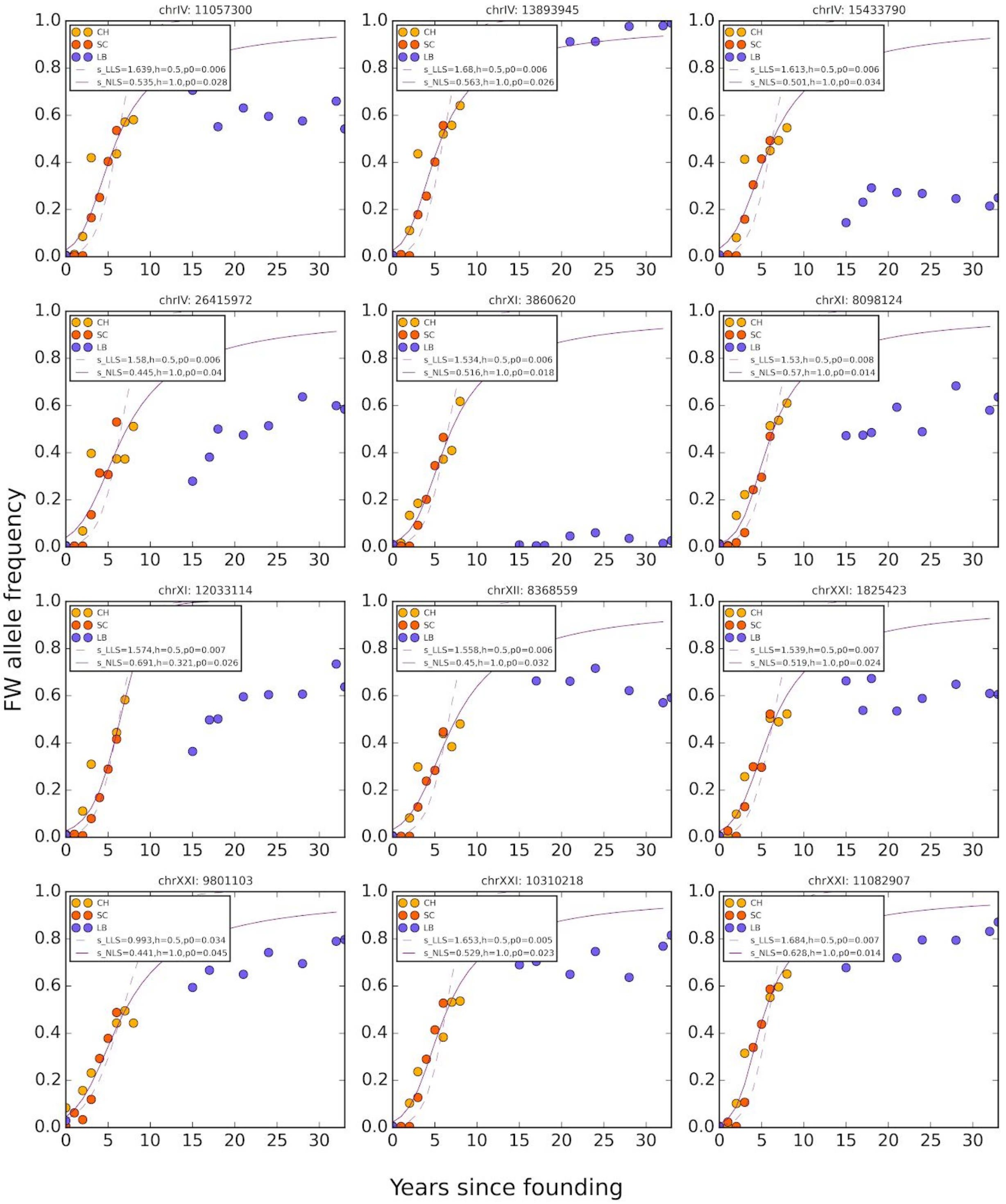


**Fig_S23.5** Allele frequency trajectories for best SNP in specific TempoPeaks for CH+SC defined by genetic distance that were absent in the corresponding analysis for LB.

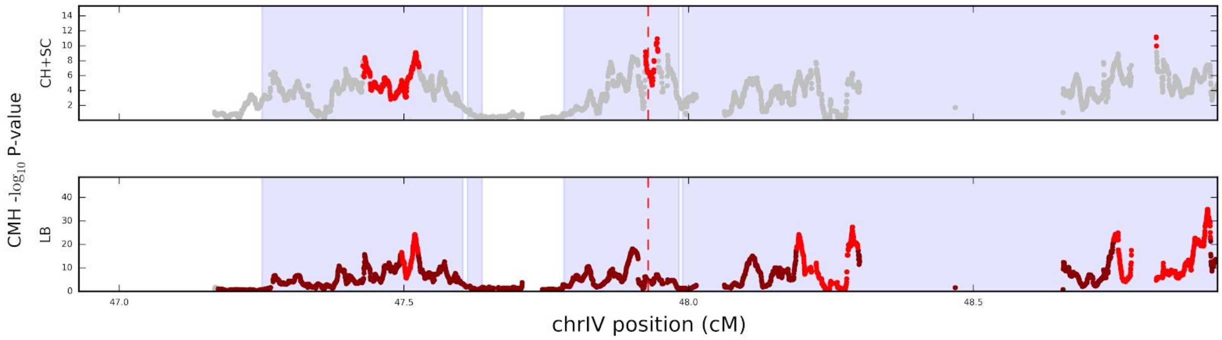


**Fig_S23.6:** CMH scores for local region of chrIV in genetic distance around SNP position 11,057,300bp (vertical red dashed line) for CH+SC (top) and LB (bottom), with regions in red and dark red found in specific and sensitive TempoPeaks respectively. Blue shading shows sensitive EcoPeak.

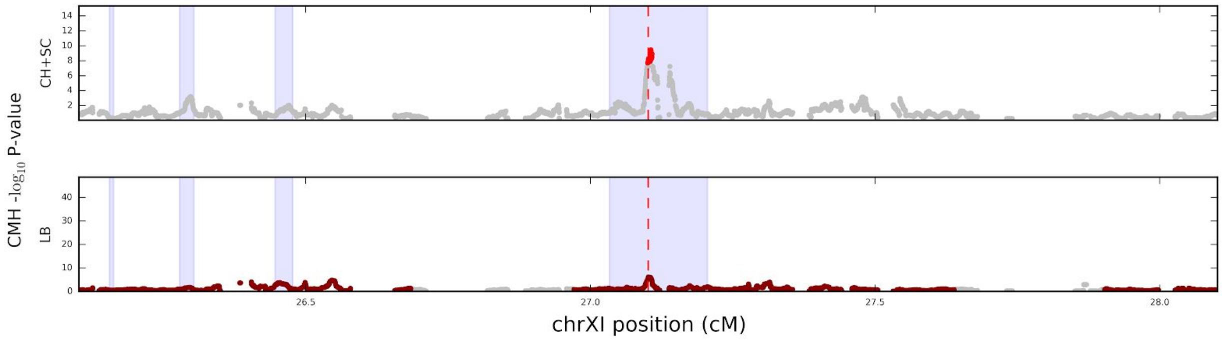


**Fig_S23.7:** CMH scores for local region of chrXI in genetic distance around SNP position 12,033,114bp (vertical red dashed line) for CH+SC (top) and LB (bottom), with regions in red and dark red found in specific peaks and sensitive peaks respectively. Blue shading shows sensitive EcoPeak.

#### 24. ESTIMATING TEMPOPEAK SELECTION COEFFICIENTS FOR CHRIV USING A DEEP NEURAL NETWORK APPROACH

Even under our specific p-value threshold and using genetic distance (our strictest criteria), there are several chromosomes with a large number of significant TempoPeaks undergoing rapid allele frequency change. In particular, for CH+SC+LB we identified 28 distinct TempoPeaks with estimated *s* values ranging from 0.08 to 0.64 and mean of 0.45 when treating each region independently. In addition, while in physical distance these TempoPeaks span 60% of the length of chrIV (**Fig S21.1**), when using genetic distance they are clustered into a much smaller region that represents only 23% of the genetic map (**Fig S21.2**). Thus, even though they are separated by recombination hotspots as defined by the local rate, the local rate is low and the TempoPeaks are very much linked. Therefore, the validity of treating SNPs in TempoPeaks in this region as independent when estimating *s* by both us (above) and others (*65*) is likely highly flawed. Multiple linked loci increasing rapidly in frequency due to positive selection will undoubtedly both interfere with each other (Hill-Robertson effects when on different initial haplotype backgrounds (*62*)) and increase the overall effect (linked selection). However, apart from simple scenarios such as two locus selection (*66*, *67*), estimating the effect of multiple linked alleles is a non-trivial task (*68*).

As a consequence, we have developed a new approach that leverages forward simulations and deep neural networks (DNN), an approach that is increasingly proving a valuable tool in population genetics for seemingly intractable problems (*69*–*71*). We are particularly motivated to examine our data within the context of the “transporter hypothesis” put forward by Schluter and Conte (*72*) that is currently the leading model for explaining how reperated adaptation in freshwater populations is maintained amongst threespine stickleback. In this model, standing genetic variants that are adaptive in a network of freshwater environments are continually recycled into a main anadromous population such that they are present at low frequencies and thus can reassemble into selected haplotypes in a new freshwater environment. Note that our goal in this analysis is not necessarily to accurately estimate $s$ values for individual loci, but rather to examine to what extent the transporter model can capture the main features of our contemporary data and the selection dynamics on the chromosome as a whole. In this regard, we estimate a distribution of selection coefficients across inferred peaks, i.e. a distribution of fitness effects (DFE).

There are effectively three main stages of this analysis: 1) simulating linked loci with positive selection parameterized with various randomly drawn DFEs within the context of the demographic and evolutionary model of the transporter hypothesis, 2) using the DNN framework to estimate the best DFE given the observed allele frequency trajectory data from our Pool-Seq experiments and 3) comparing the transporter hypothesis under the best fit DFE to various features of the observed genomic data.

*Simulations.*

We used `SLIM` (v3) (*73*) to conduct all forward simulations under a diploid Wright-Fisher framework with non-overlapping generations. Our demographic model representing the transporter hypothesis involved two contrasting habitats, with an initial set of 10 freshwater water populations each with $Ne_{fw}$=1,000 diploid individuals, and 1 anadromous population with $Ne_{an}$=10,000 diploid individuals. Bi-directional migration was allowed between each freshwater population and the anadromous population (but not between freshwater habitats). Migration per generation was set such that $m_{\mathrm{an>fw}}$=0.01 and $m_{\mathrm{fw>an}}$=0.001. In this way the total number of individuals from each habitat contained the same overall population sizes and received the same number of migrants from the opposing habitats, $nm$=100. This setup is conceptually similar to that implemented by Galloway et al. (*35*) in their representation of the transporter hypothesis.

At the beginning of the simulation, one freshwater population, $p1$, is chosen to be seeded with freshwater adaptive alleles. 200 chromosomes (i.e. 10%) are randomly selected to possess a haplotype containing all freshwater adaptive alleles. Note therefore we are not simulating the emergence of freshwater alleles and haplotypes (and no new mutations are allowed throughout the simulation). Instead we are assuming the alleles have already established in freshwater stickleback and arranged themselves to form full length haplotypes with optimum fitness. We seed $p1$ at 10% to ensure the freshwater alleles are not lost due to drift at the beginning of the simulation. Each freshwater allele is assigned some positive $s$ from a DFE. If this allele enters the anadromous population, it is assigned 1/fitness of that in the freshwater habitat, and we assume that the anadromous population is essentially at a constant optimum fitness except for the impact of any of these migrant freshwater alleles.

The population then evolves for 1,000 generations, which we found through experimentation to be sufficient time to ensure that freshwater alleles, when the DFE allows it, can establish across all 10 freshwater populations and reach an overall equilibrium across the whole system in terms of there being no allele frequency changes in each population. After 1,000 generations, new freshwater habitats with $Ne$=1,000 are established from copies of individuals from the anadromous population to represent Loberg ($t$=1,001), Cheney ($t$=1,022) and Scout ($t$=1024). Unlike the original 10 freshwater habitats, no migration is allowed to or from these populations. The simulation terminates after 1,030 generations.

Given the difficulty of fitting multiple linked selective loci simultaneously, we identified a new set of significant TempoPeaks on chrIV using the specific p-value threshold with windows defined by genetic distance as before, but increasing the required minimum distance between peak boundaries from 50kb to 100kb. This resulted in 19 TempoPeaks rather than the original 28, thus reducing the dimensionality of our inference somewhat. The characteristics of the 19 SNPs with the highest CMH scores in each of these peaks formed the basis of the genomic architecture of our simulations.

We simulated 19 biallelic SNPs with the same physical distance and recombination rate as the observed peak SNPs. To determine our DFE, for any given simulation we drew 19 random values from a beta distribution with two parameters ($\alpha$ and $\beta$). These values were assigned as the $s$ values for each of the simulated 19 freshwater-adaptive alleles, preserving the rank order of the estimated $s$ values when treating each SNP as independent (i.e. the largest of the 19 random values drawn from the beta distribution was assigned to the SNP with the highest estimated $s$). For the beta distribution, the $\beta$ parameter was drawn on a $\log_{10}$ scale from -1 to 5 (so from 0.1 to $1\times10^5$). The $\alpha$ was then drawn on a $\log_{10}$ scale from -1 to the previously chosen $\beta$ values. As the mean of a beta distribution is $\alpha/(\alpha + \beta)$, this restricts the mean $s$ value for any simulation to be a maximum of 0.5.

To assign dominance coefficients for each freshwater allele, we first drew a random integer between 0 and 19 from a uniform distribution to represent the number of SNPs with $h$=0 ($n_{h=0.0}$), followed by another random integer between 0 and ($n_{h=0.0}$-1) to represent the number of SNPs with $h$=0.5 ($n_{h=0.5}$). The remaining 19-($n_{h=0.0}$+$n_{h=0.5}$) SNPs were then assigned $h$=1. SNPs were assigned *h* values in the reverse rank order of the estimated $s$ values (i.e. the SNP with the highest $s$ was assigned the *h* with the lowest dominance. Our approach draws from a model that suggests alleles with greater effects on fitness will be associated with lower dominance coefficients (*74*, *75*). We note that this hypothesis is usually contextualized from the perspective of deleterious effects, which would be the case in the anadromous population.

The primary output of our simulations was an estimated allele frequency for each freshwater-adaptive allele from a sample of 100 individuals from each time point matching our contemporary time series. As in the CMH test, RS2009 was utilized as a proxy for the founding Loberg population. Overall there were 19 × (7+7+9) = 437 distinct allele frequency estimates generated per simulation, which are equivalent to 437 allele frequency estimates from our real Pool-Seq data. Note that we do not simulate an aspect of pooling and 2nd generation sequencing in our simulated data (and so additional overdispersion of the variance), while also not matching sample sizes exactly (3 out of our 23 Pool-Seq samples consisted of less than 100 samples). Instead, we make the assumption that due to our large sample sizes and coverage overall, that our observed Pool-Seq allele frequencies are a good estimator of the true sample allele frequencies.

We were able to conduct 676,703 simulations under this framework, with each simulation generated using randomly selected $\alpha$,$\beta$,$n_{h=0.0}$ and $n_{h=0.5}$ parameters and producing an output of 437 allele frequencies.

*DNN inference*

We utilized `Keras` (v2.3.1) running on top of `tensorflow` (v2.1.0) to use our simulations to train a DNN that aimed to predict our 4 input parameters (the output layer) given the 437 allele frequencies (the input layer). We experimented with various DNN parameterizations and found the following setup to produce the most reliable result (**Fig S24.1**): 3 fully connected hidden layers each with 1024 nodes, a batch size of 32, an elu activation function, a He normal kernel initializer, an ADAM optimizer and a Huber loss function (*76*). 10,000 simulations were first set aside for later validation analysis. From the remaining simulations, 80% were used for training and 20% for validation by the DNN. The 437 allele frequencies were normalized to have zero mean and unit variance before being used as input for the DNN. The model was allowed to run for a maximum of 100 epochs, with early stopping allowed if the training performance did not improve for 3 consecutive epochs. Once trained, the DNN was applied to the observed Pool-Seq allele frequency data to estimate $\alpha$,$\beta$,$n_{h=0.0}$ and $n_{h=0.5}$.

We found that a key aspect for successfully training our DNN was filtering our simulated data prior to training to remove regions of the parameter space with low information content. Given that we know that our true data includes SNPs with very high freshwater allele frequencies in many of our time points because of the high estimated $s$ values (with some for Loberg approaching fixation), we examined how many simulations contained at least one sampled time point with a freshwater allele frequency above some threshold $x$ as a function of $\alpha$ and $\beta$ on a $\log_{10}$ scale. Starting with an $x$ of 0.1 (i.e. 10%), we see that the highest density is for simulations with both $\alpha$ and $\beta$ close to 0.01 (**Fig S24.2**). There is generally a higher density of simulations that meet our threshold along a strip of parameter space where $\log_{10}(\beta)$ - $\log_{10}(\alpha)$ < 2. As we increase $x$ to 0.25 (**Fig S24.3**) and 0.5 (**Fig S24.4**) we see this strip becoming increasingly prominent, to the extent that is does not appear that any simulation where $\log_{10}(\beta)$ - $\log_{10}(\alpha)$ is greater than 2 can produce even a single time point amongst our three populations where at least one freshwater allele frequency is > 0.5. This area of the parameter space reflects a mean *s* < 0.01. Thus, before training our DNN, we filtered out all simulations where $\log_{10}(\beta)$ - $\log_{10}(\alpha)$ < 2, and thus we are restricting our mean *s* across all 19 loci to being between 0.01 and 0.5 (note that increasing the absolute values of $\log_{10}(\beta)$ and $\log_{10}(\alpha)$ but keeping the difference the same simply decreases the variance of the DFE), with our primary DNN training being conducted on 474,019 simulations.

In order to examine the effect of filtering on parameter estimates, we trained our model on simulated data both with and without filtering 100 times, each time varying the random seed to initiate the training. For each of the 100 DNNs we then predicted the four target parameters from the 10k simulations we had withheld before training. For each parameter we then performed a linear regression of the known true value for each simulation against the mean, 5th percentile and 95th percentile of the 100 DNN estimates. The predictions from the DNN trained with filtered data (**Fig S24.5**) clearly has improved performance compared to the unfiltered data (**Fig S24.6**) for all four parameters. For the unfiltered data there appears to be a large subset of simulations where the beta parameter estimates condense into a single predicted value despite the true values spanning the entire available range. There are also other regions of the parameter space that generally perform poorly and form clouds. These areas are almost certainly the result of the simulations in the parameter space where we found there to be little allele frequency information. The correlation coefficients between the true data and estimated means are much higher for the filtered data (0.80 to 0.91 versus 0.50 to 0.76) and the linear regression models are much closer to an $x = y$ relationship. In the case of the filtered data, the estimated values are fairly unbiased for lower $\log_{10}(\alpha)$ and $\log_{10}(\beta)$ values, starting to increase their variance and somewhat underestimate the true value systematically when $\log_{10}(\alpha)$ and $\log_{10}(\beta)$ are greater than 2 and 3 respectively. With regard to the dominance coefficients, though there is a large variance, the DNN does a better job of estimating $n_{h=0.0}$ than $n_{h=0.5}$ with the latter being systematically more underestimated.

We then used our 100 DNNs trained using filtered simulated data to estimate the four target parameters from our observed data. We used `fastkde` (v1.17.1) (*77*, *78*) to calculate a four dimensional kernel density across our 100 iterations and used the point with the highest density to get our final parameter estimates, as well as the mean and standard deviation of $s$ (**Table S24.1**). We note that our beta distribution parameter estimates are in the range where the DNN appears to perform particularly well on simulated data (**Fig S24.5**). Examination of the 2-dimensional density kernel for $\log_{10}(\alpha)$ and $\log_{10}(\beta)$ estimates shows that the two parameters are highly correlated, such that the estimated mean value of the DFE is fairly restricted (**Fig S24.7**), though there is variation in the variance from run to run (**Fig S24.8**). The distribution of the estimates of the relative number of recessive versus semi-dominant (and thus also dominant) SNPs are fairly constrained and support there being no recessive freshwater alleles and between 1 and 2 semi-dominant alleles (**Fig S24.9**).

*Validation versus real data*

In order to examine how well our DNN-based model was able to recreate patterns in our observed data, we first simulated our transporter model using the estimated parameters from **Table S24.1** 100 times. As $n_{h=0.0}$ *and* $n_{h=0.5}$ are fixed integers as input but are output as probabilistic floating point numbers by the DNN, we performed 100 simulations with $n_{h=0.5} = 1$ and $n_{h=0.5} = 2$ separately. In both cases, the range of allele frequencies outputted by the simulations for each lake and time point overlap well with the real data, though $n_{h=0.5} = 1$ seems to perform slightly better visually (**Fig S24.10** and **Fig S24.11**), and had a lower mean absolute error (0.157 vs 0.214). The main discrepancy between the observed versus simulated results, was for chrIV:16,366,165 where the simulations did not capture the lower frequencies for Loberg. This is not unexpected as this SNP is in one of the regions described previously as being significant only in CH+SC, but not LB. In addition, the first and last SNP (chrIV:8,359,882 and chrIV:28,894,879) have simulated allele frequencies that are generally slightly lower (but still captured by the range) than the observed data across time points.

We next simulated from our best parameter model with $n_{h=0.5} = 1$, but additionally introducing neutral SNPs into the simulations alongside our 19 selected sites in order to examine the pattern of CMH scores across the entire chromosome. To mimic realistic levels of genetic variation across the chromosome without having to simulate *de novo* mutations (which would require substantial computational burden with a significant burn in period), we first identify all SNPs from the long-term set that were polymorphic in our RS2009 data set. We then randomly select 400,000 of these SNPs and randomly place them in any of our 10 freshwater or 1 anadromous starting population at the same frequency we observe them in RS2019. They are given the same physical and genetic position as in the real data (i.e. we are applying the same recombination landscape). Following the end of the simulation, we then calculate individual CMH scores as for the observed data (though without the correction for overdispersion due to read sampling (*60*)), and then plot the distribution of p-values within 2500bp windows sliding every 50bp.

As this simulation process is computationally intensive, we performed this simulation 10 times. The p-value landscapes in physical and genetic distance for 5 of these iterations are shown **Fig S24.12**, **Fig S24.13** and **Fig S24.14** alongside the original observed data, while Fig 4D shows the result of average the p-value for each window across the 10 simulations. The simulations visually show much of the same topology in p-values across the genome (though with perhaps less peakedness), especially when utilizing genetic distance. Thus it appears that the model inferred from our DNN can capture the overall genomic architecture of linked neutral SNPs, even if they were not explicitly part of the model. One interesting observation is that while the 19 selected SNPs are almost always in the vicinity of a CMH p-value peak, they are not always the most significant p-value themselves, showing that closely linked neutral alleles can often hitchhike to produce allele frequency trajectories that are even more extreme than the actual allele under positive selection.

Finally, we examined to what extent our best fit DNN model under the transporter hypothesis could produce haplotype patterns observed in our SNP array data for Rabbit Slough (n=750 diploid individuals) as well as Loberg 1999 (n=25) and Loberg 2013 (n=25). We performed our simulations 1000 times, each time outputting haplotypes at our 19 loci matching the time points and sample sizes of our three array data set. In the case of RS2009, we sampled haplotypes from the anadromous population at the time point directly before initiating our freshwater habitat equivalent to Loberg Lake (so $t$=1,000). As expected, there was considerable variability from run to run. We show 3 representative simulations in **Fig S24.15** and **Fig S24.16**. In addition, we obtained an average haplotype behaviour by keeping all diploid individuals from across the 1000 simulations (so in the case of Rabbit Slough we generated 1000 *750 individuals), and then resampling down from these larger sets to match original samples sizes. Therefore, any haplotype structures that are prominent across many simulations would more likely be represented in this subset. These average haplotype plots are shown in Fig 5B, alongside the equivalent plots for the observed array data. Note we represent the haplotypes as diploid genotypes in individuals over the genome as in the array data, and thus we do not require any phasing of the observed data to make comparisons.

In general, our simulations correspond well to the observed data. For Rabbit Slough, our simulated anadromous populations capture the prominence of the majority of individuals being 100% anadromous in terms of genotypes across chrIV (all red rows), with a much smaller proportion containing regions of heterozygosity for freshwater adaptive alleles of varying tract lengths (yellow tracts). Similarly for Loberg 1999 and 2013, our simulations capture the transition from primarily heterozygote genotypes to homozygote freshwater tracts between these two points (from a mix of yellow and blue tracts to majority blue), as well as the asymmetry of the left hand side of chrIV seemingly evolving quicker than the right half in terms of individuals acquiring freshwater alleles.

**Table S24.1.** Best parameter estimates from our DNN analysis using multidimensional kernel density estimation

| Parameter | Estimate |
|---|---|
| $\log_{10}(\alpha)$ | 0.607 |
| $\log_{10}(\beta)$ | 1.781 |
| $n_{h=0.0}$ | 0.012 |
| $n_{h=0.5}$. | 1.664 |
| Mean and std of $s$ | 0.063, 0.030 |

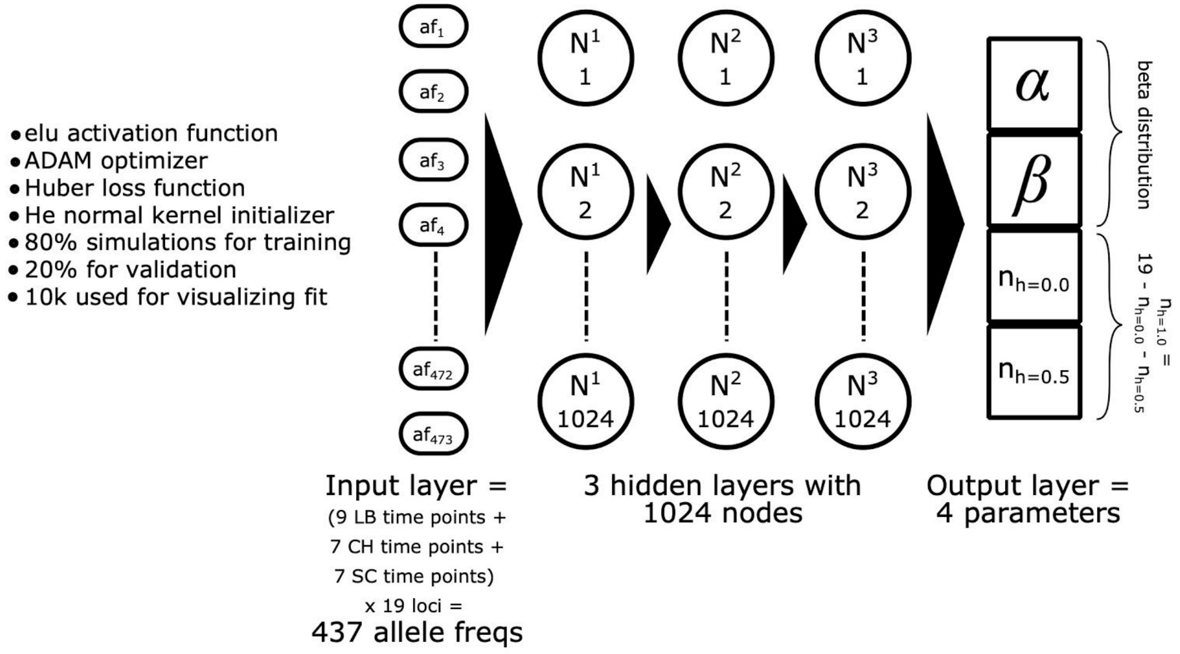


**Fig S24.1**: Figure showing the parameterization and structure of our DNN.

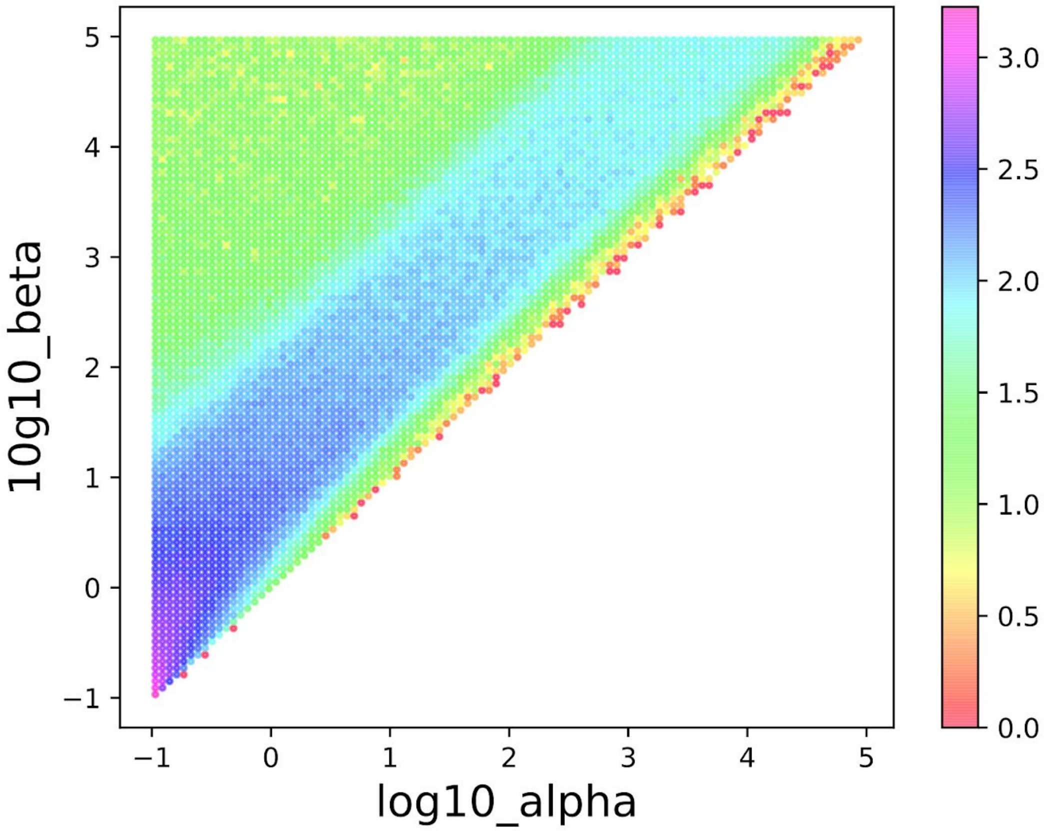


**Fig S24.2:** Two-dimensional histogram showing the number of simulations (in log10 color scale) where the freshwater allele frequency in at least one sampled time point was greater than 0.1, as a function of log10(alpha) and log10(beta) parameters of the beta distribution.

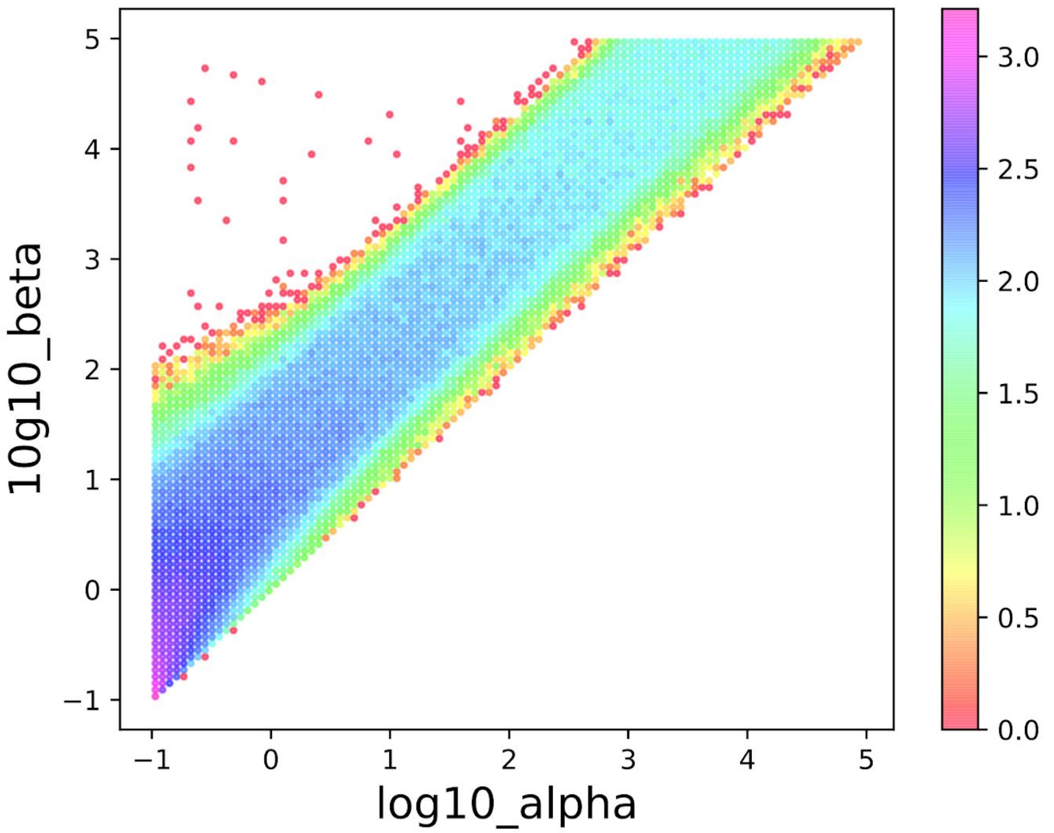


**Fig S24.3:.** Two-dimensional histogram showing the number of simulations (in log10 color scale) where the freshwater allele frequency in at least one sampled time point was greater than 0.25, as a function of log10(alpha) and log10(beta) parameters of the beta distribution.

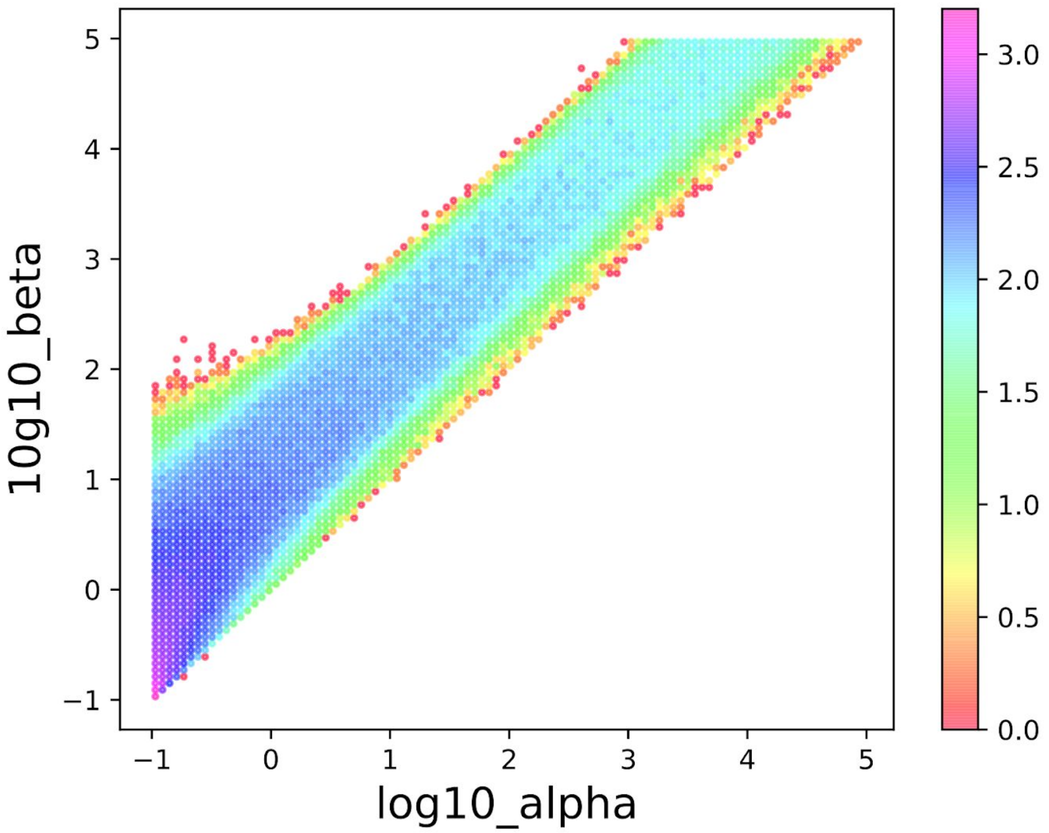


**Fig S24.4:.** Two-dimensional histogram showing the number of simulations (in log10 color scale) where the freshwater allele frequency in at least one sampled time point was greater than 0.5, as a function of log10(alpha) and log10(beta) parameters of the beta distribution.

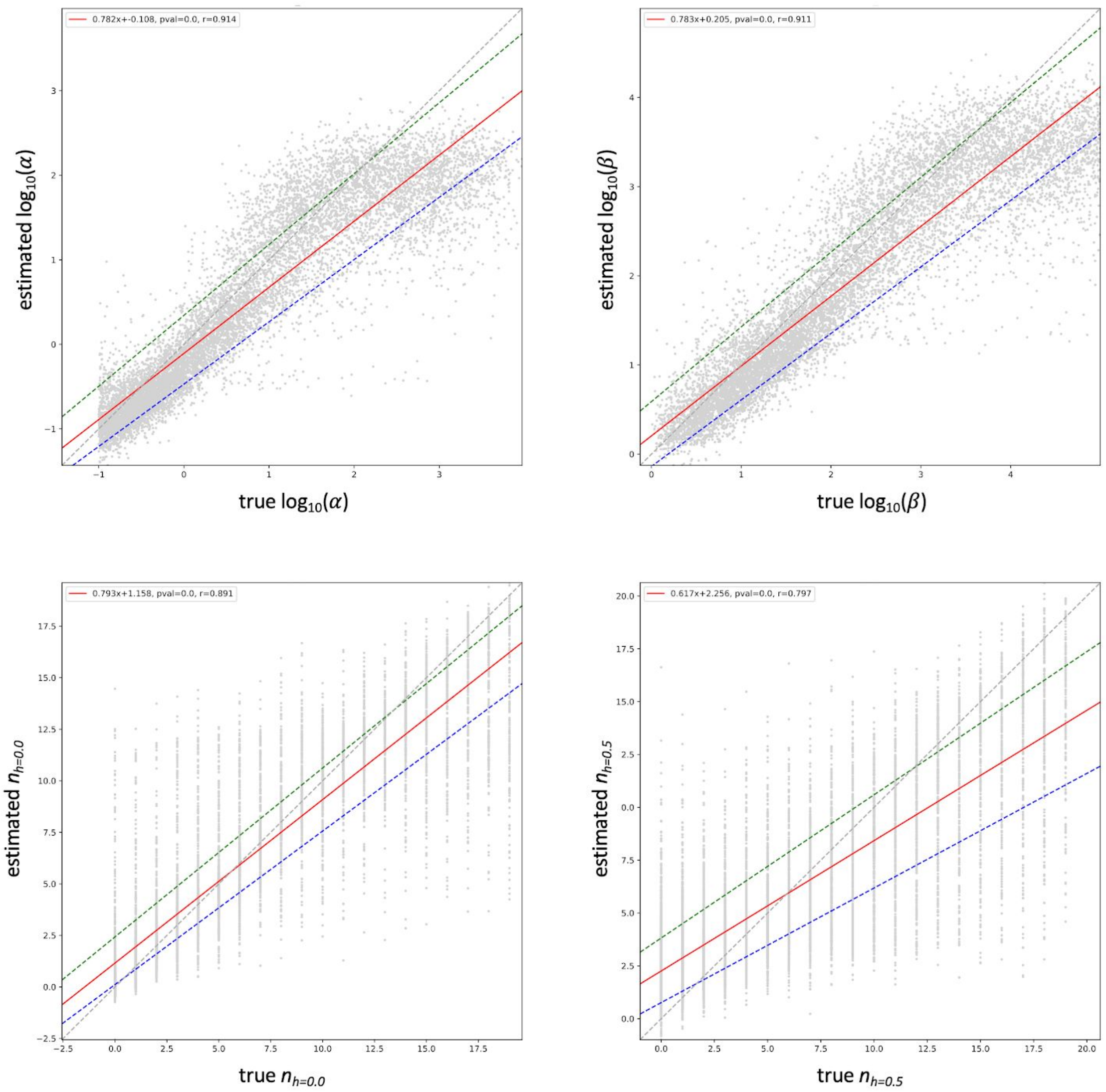


**Fig S24.5:** Comparison of true parameter values from 10k simulations after filtering versus mean of those predicted by DNN from 100 runs. Dashed grey line is the expected relationship, red, blue and green lines are linear regression models for the mean, 5th and 95th percentile of the 100 DNN estimates. Inferred regression equation for mean show in top right for each parameter.

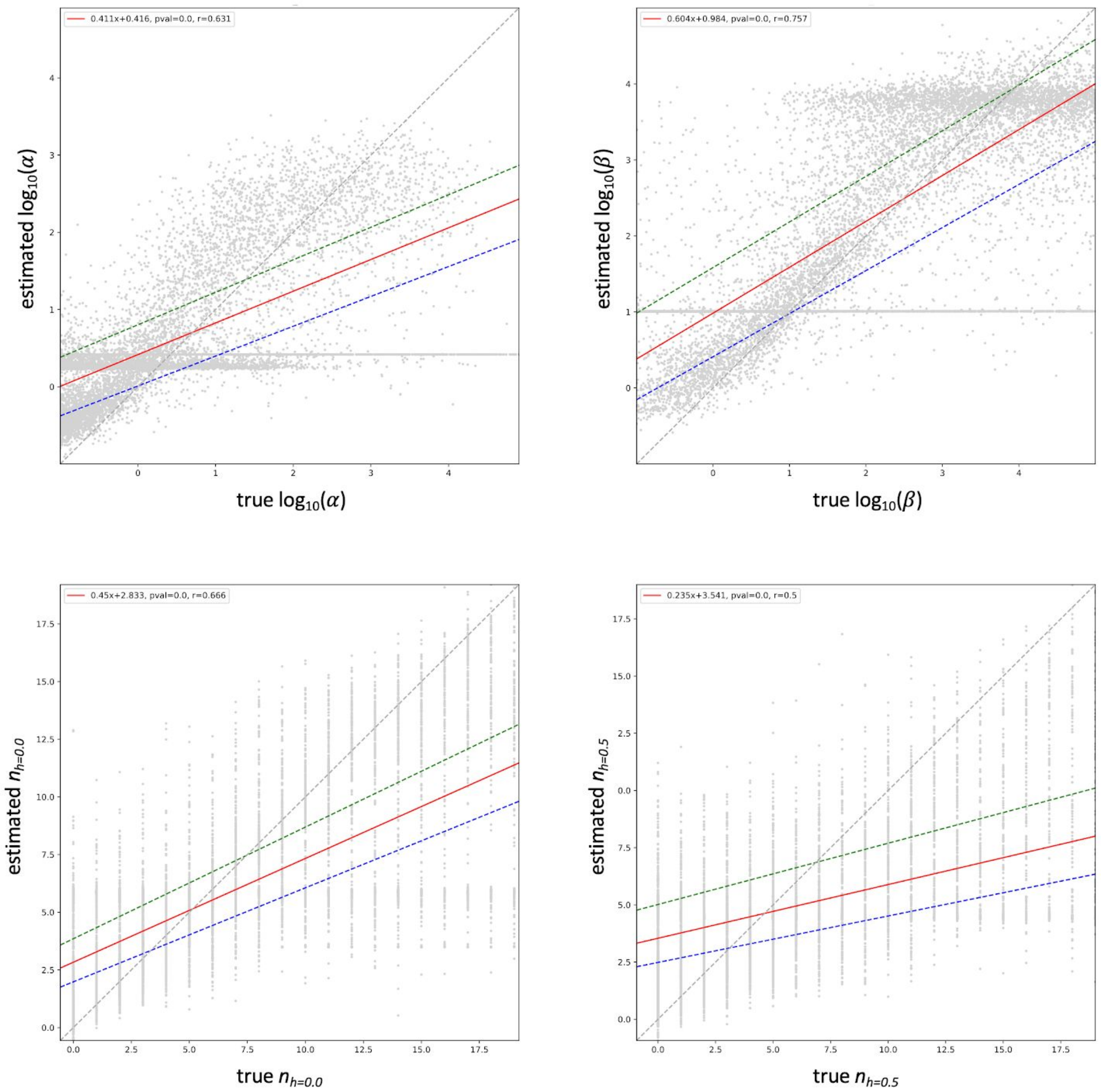


**Fig S24.6:** Comparison of true parameter values from 10k simulations without filtering versus mean of those predicted by DNN from 100 runs. Dashed grey line is the expected relationship, red, blue and green lines are linear regression models for the mean, 5th and 95th percentile of the 100 DNN estimates. Inferred regression equation for mean show in top right for each parameter.

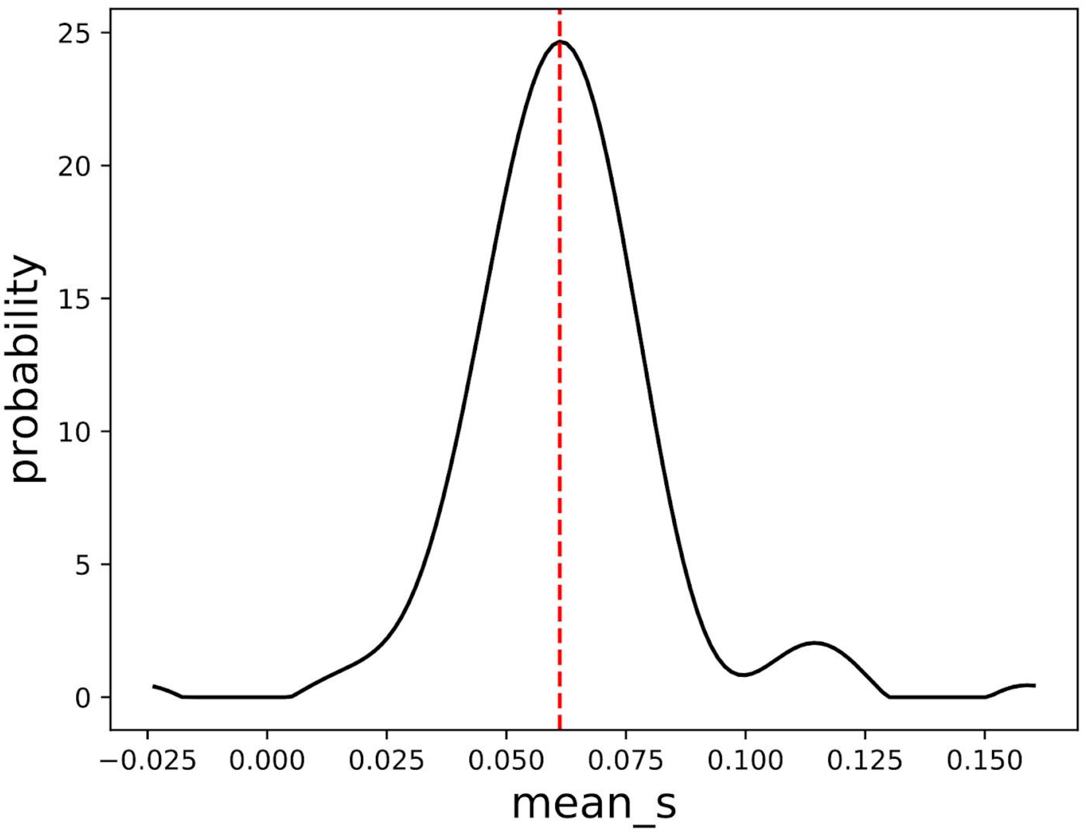


**Fig S24.7:.** Distribution of mean $s$ based on beta distribution parameter estimates across 100 DNN models. Red dashed line represents multidimensional kernel density estimate.

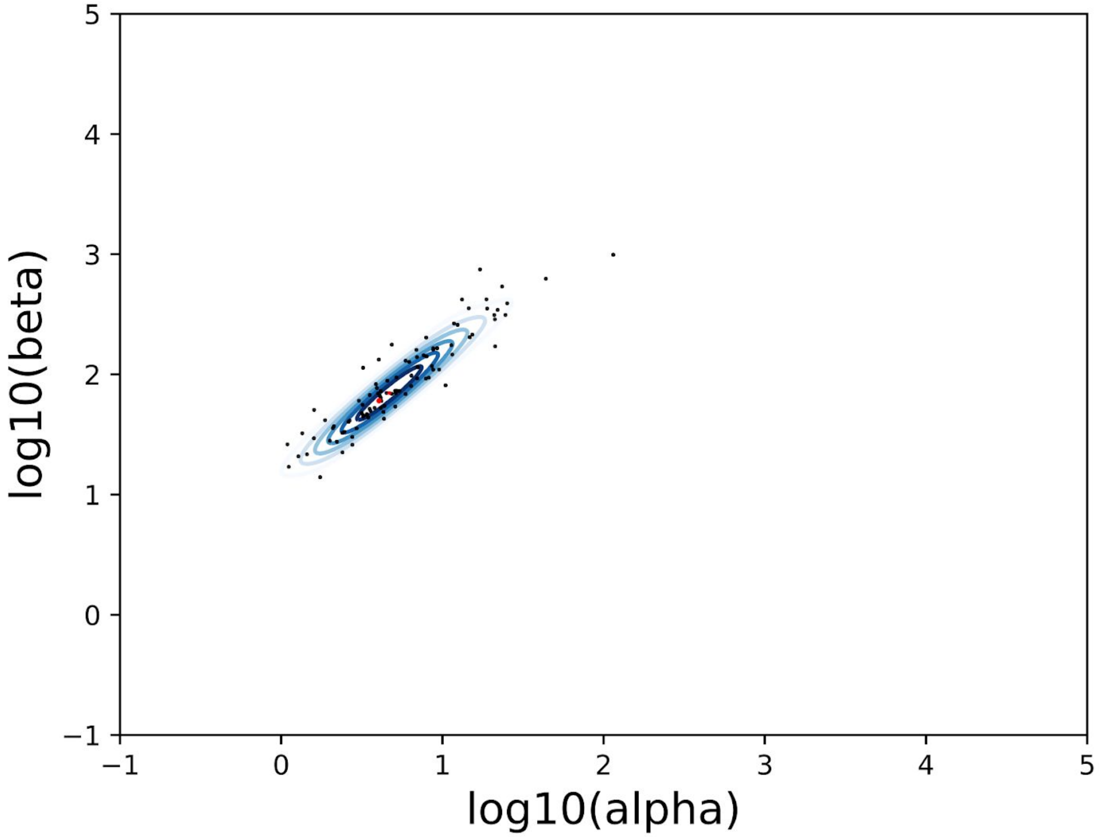


**Fig S24.8:** Contour plot of two-dimensional kernel density for log10(alpha) and log10(beta) parameters of the beta distribution. Red circle and star represent points with highest density when estimating a four-dimensional and two-dimensional kernel density respectively.

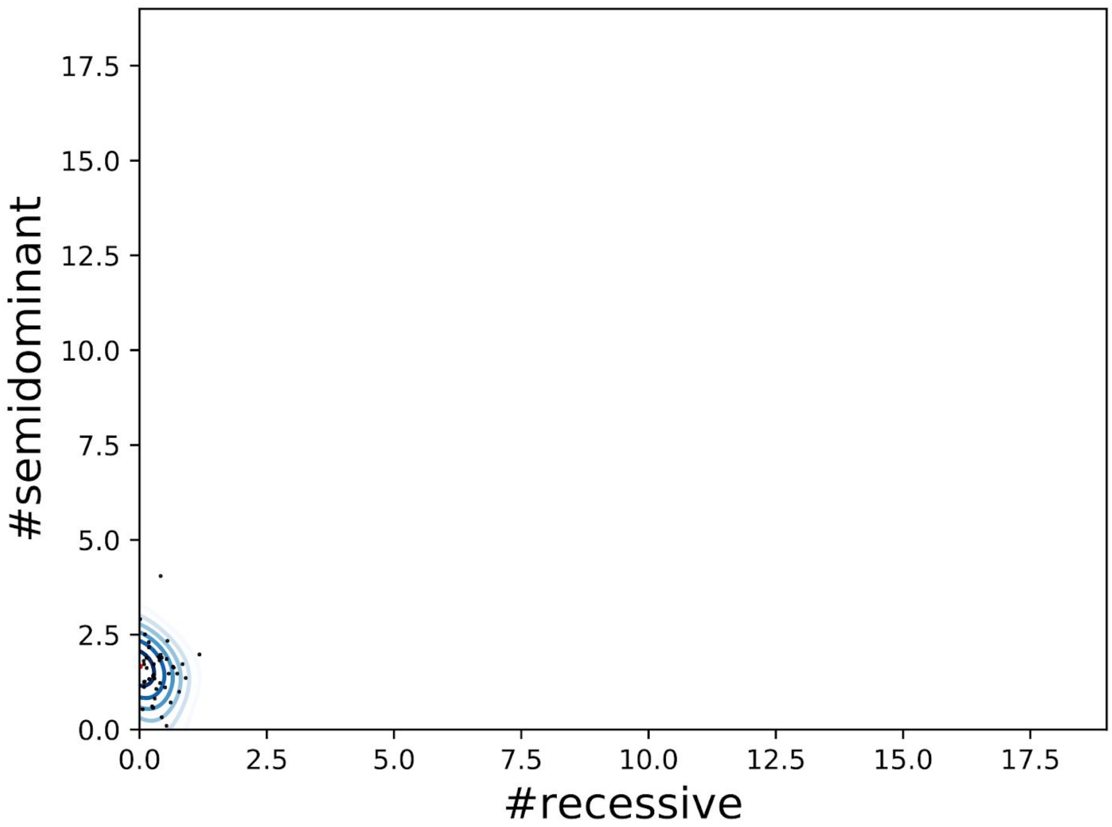


**Fig S24.9:** Contour plot of two-dimensional kernel density for n_h=0.0 and n_h=0.5 parameters. Red circle and star represent points with highest density when estimating a four-dimensional and two-dimensional kernel density respectively.

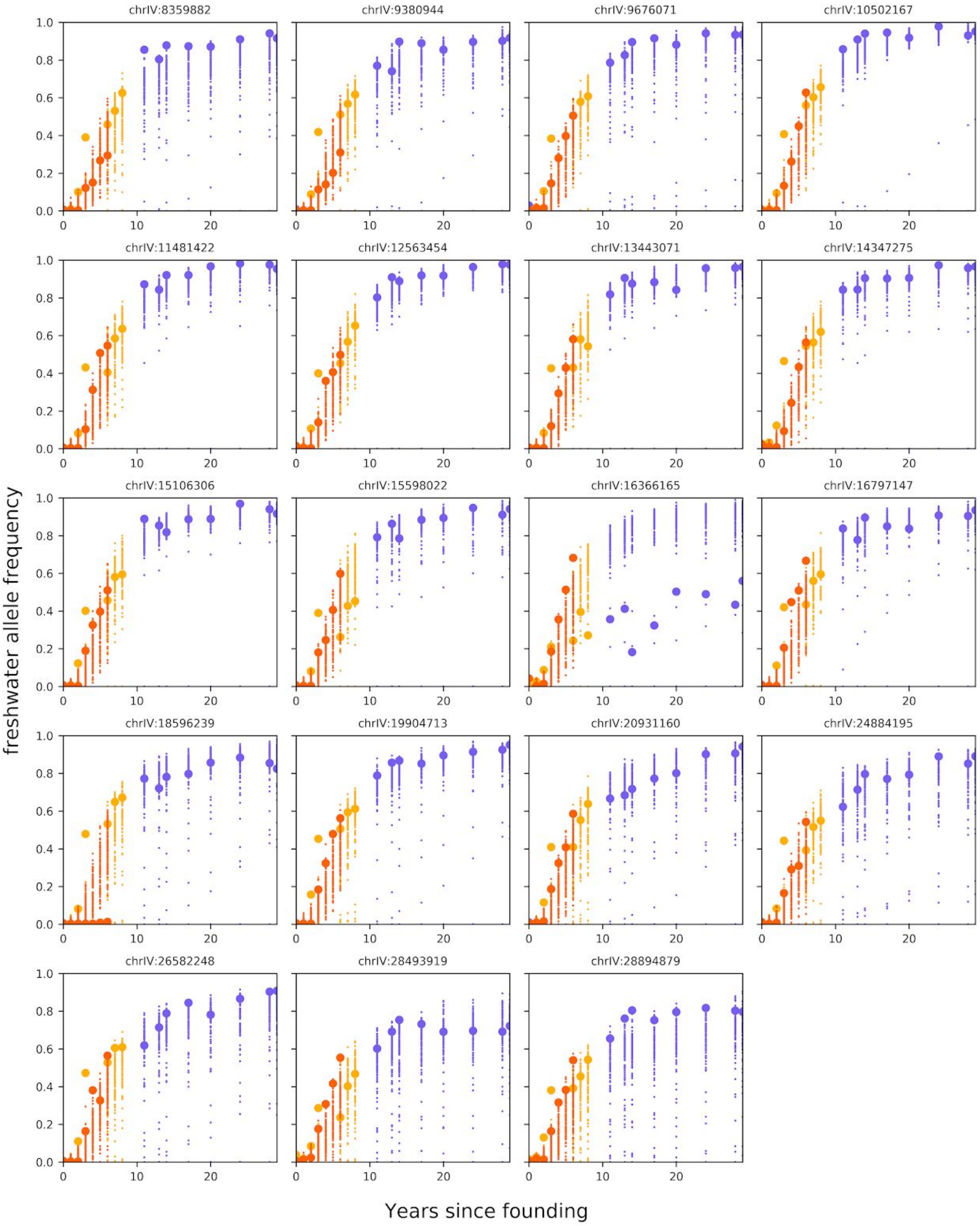


**Fig S24.10:** Observed allele frequencies from time-series Pool-Seq data for three contemporary lakes (large dots, same color scheme as Fig 1) at the 19 specific TempoPeak SNPs determined using genetic distance plotted against equivalent allele frequencies generated from 100 simulations under the best fit model from the DNN with n_h=0.5 set to 1.

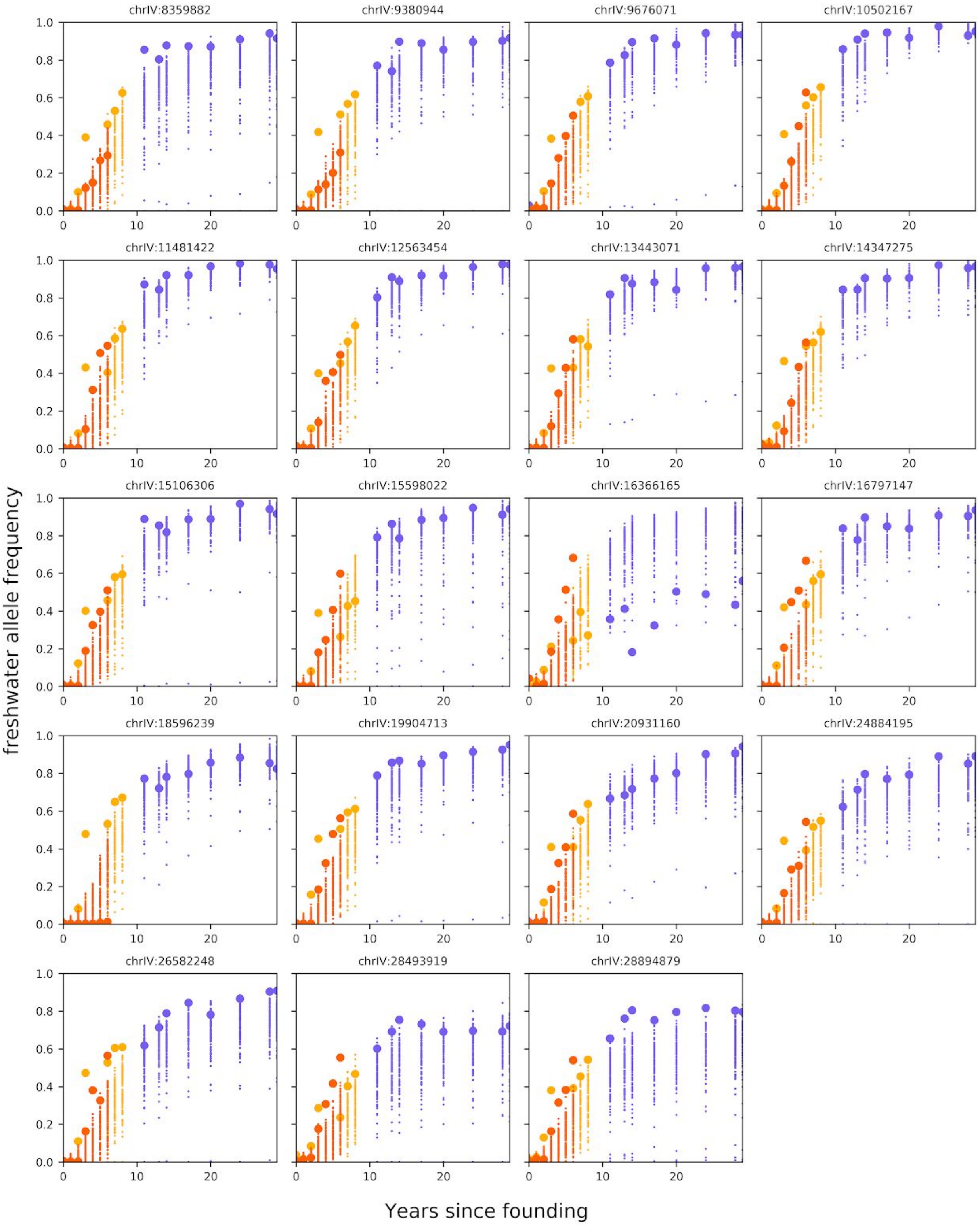


**Fig S24.11:** Observed allele frequencies from time-series Pool-Seq data for three contemporary lakes (large dots, same color scheme as Fig 1) at the 19 specific TempoPeak SNPs determined using genetic distance plotted against equivalent allele frequencies generated from 100 simulations under the best fit model from the DNN with n_h=0.5 set to 2.

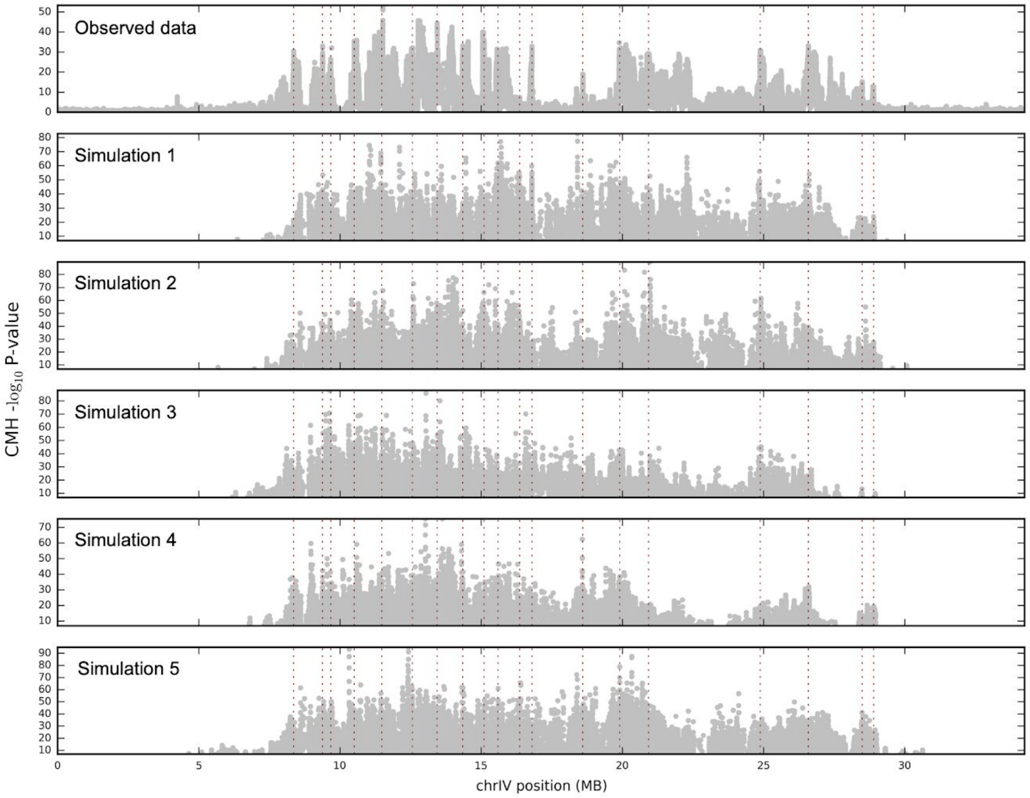


**Fig S24.12:** Genome-wide distribution of mean CMH -log10 p-values on chrIV in windows of 2500bp sliding every 50bp for the observed CH+SC+LB as well as five simulated data sets using the best fit parameters from our DNN model. Position of each selected SNP used to train the DNN are shown with the red dashed vertical line.

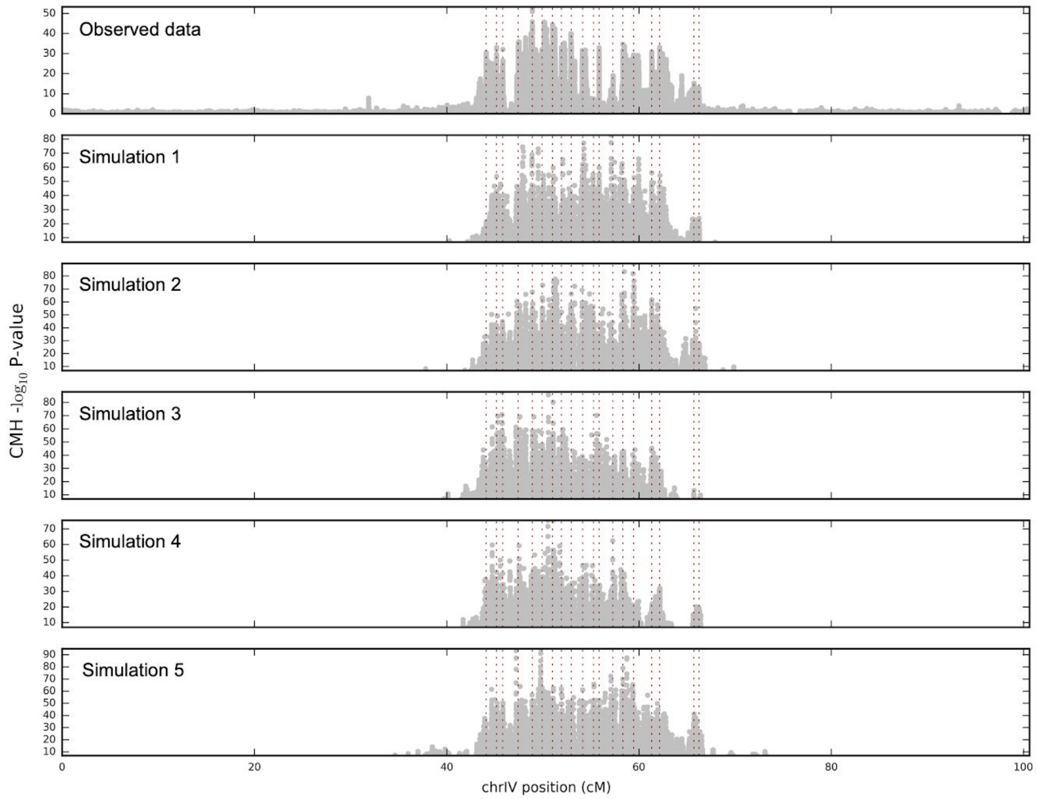


**Fig S24.13:** Genome-wide distribution of mean CMH -log10 p-values on chrIV in windows of 0.01cM sliding every 0.0002cM for the observed CH+SC+LB as well as five simulated data sets using the best fit parameters from our DNN model. Position of each selected SNP used to train the DNN are shown with the red dashed vertical line.

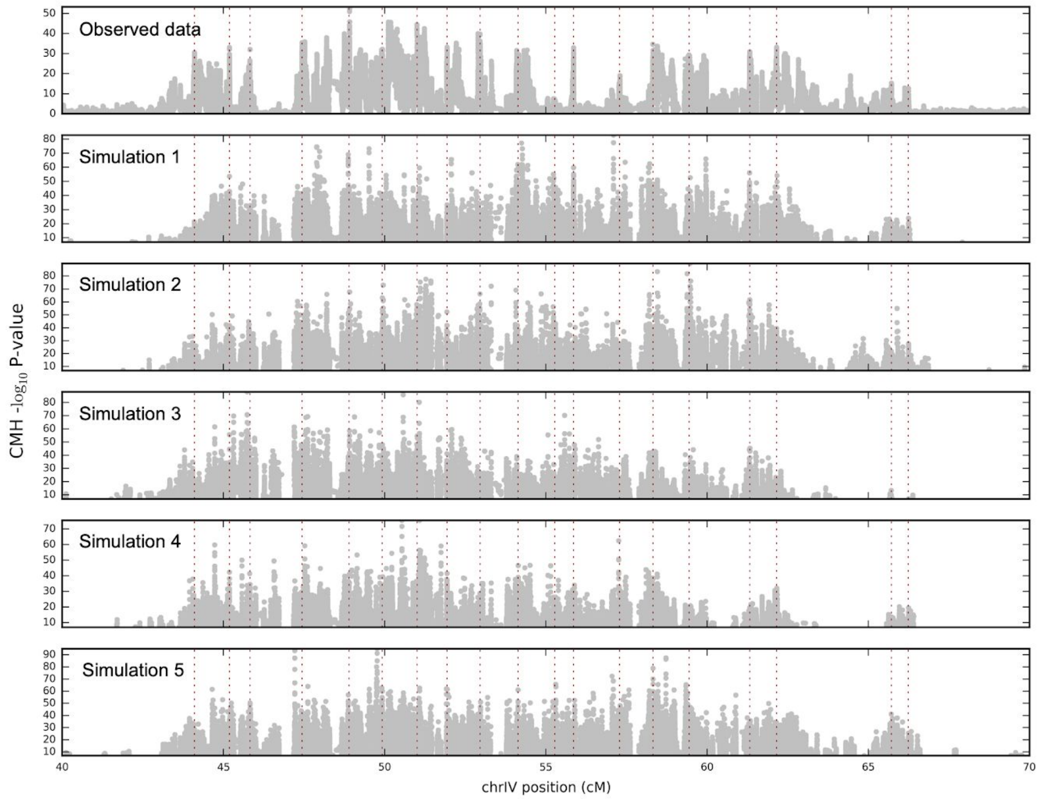


**Fig S24.14:** Distribution of mean CMH -log10 p-values on chrIV zooming in from 40 to 70 cM in windows of 0.01cM sliding every 0.0002cM for the observed CH+SC+LB as well as five simulated data sets using the best fit parameters from our DNN model. Position of each selected SNP used to train the DNN are shown with the red dashed vertical line.

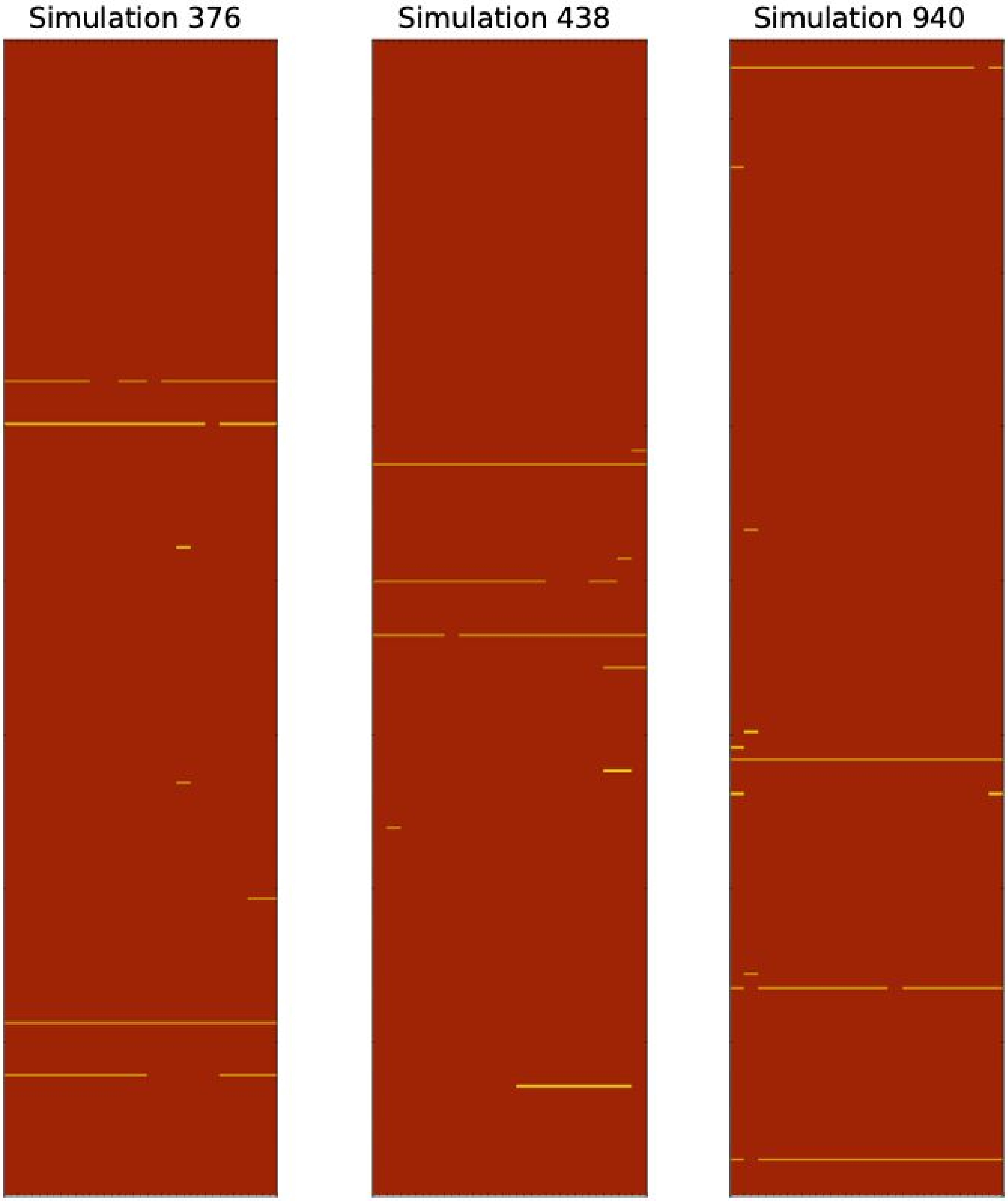


**Fig S24.15:** Sampled diploid genotype haplotypes across 19 SNPs associated with freshwater adaptation from anadromous population representing Rabbit Slough from 3 representative simulations under the best fit model estimated using our DNN. Red represents homozygote genotypes for the anadromous alleles, yellow represents heterozygote genotypes and blue represents homozygote genotypes for the freshwater alleles.

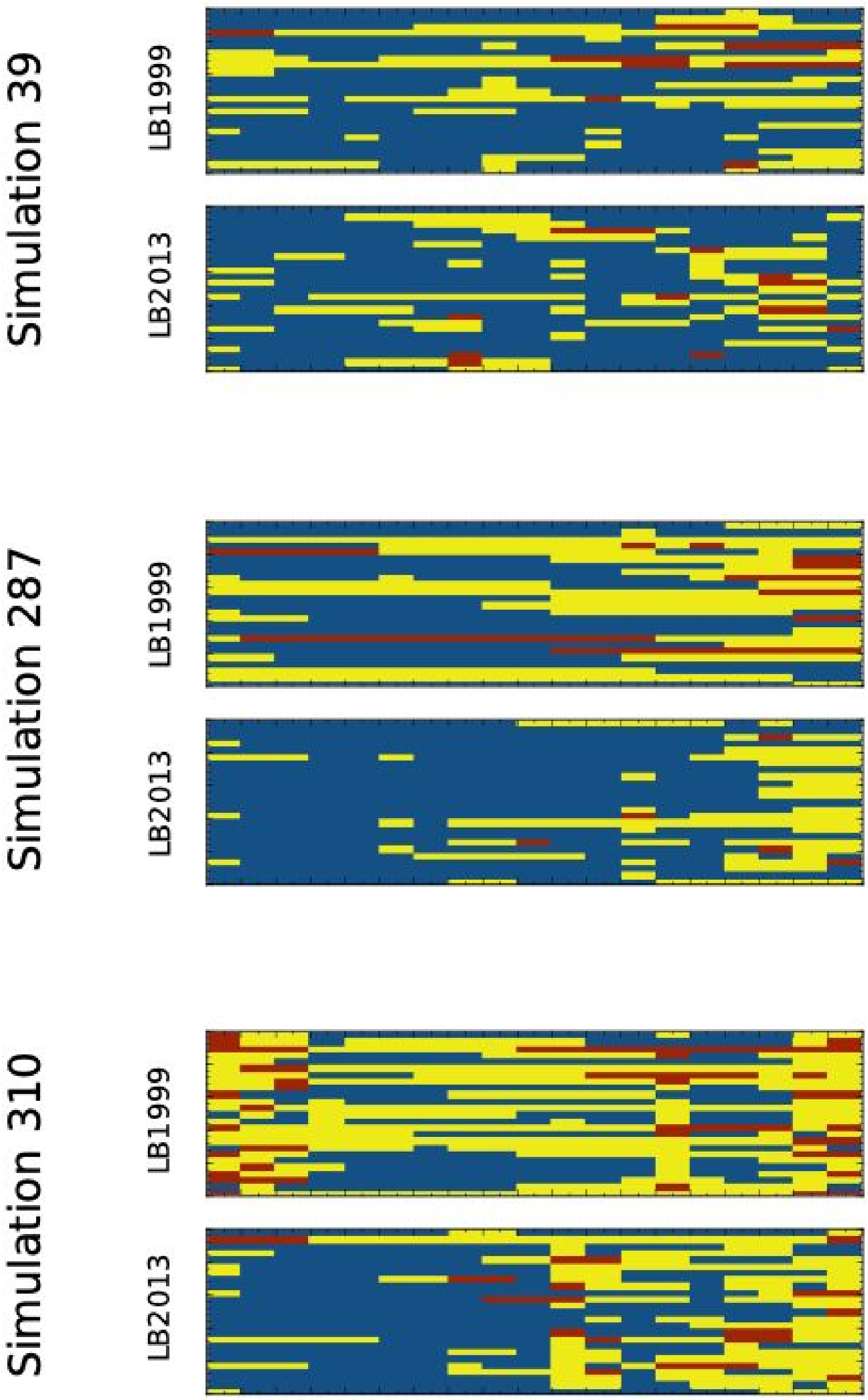


**Fig S24.16:** Sampled diploid genotype haplotypes across 19 SNPs associated with freshwater adaptation from anadromous population representing Loberg 1999 and 2013 from 3 representative simulations under the best fit model estimated using our DNN. Red represents homozygote genotypes for the anadromous alleles, yellow represents heterozygote genotypes and blue represents homozygote genotypes for the freshwater alleles.

#### 25. PREDICTING SPEED OF SELECTION IN STICKLEBACK

We collated data on the following for every specific CH+SC+LB and CH+SC TempoPeak: TempoPeak size (in both bp and cM); frequency of differentiation in the established geographic survey populations (best p-value within the TempoPeak in the EcoPeak analysis, by SNP- and window-based analyses, as well as z-score transforms thereof, in both the northeast Pacific and globally); speed of differentiation in the contemporary evolution experiments (best CMH p-value within the TempoPeak in all of CH, SC, LB, CH+SC, and CH+SC+LB); variants called (total, significant [$p < 10^{-6}$], and both previous per kb); genic overlap (gene count, gene count/kb, and a binary overlap indicator); overlap with major QTLs (both number overlapping the region and a binary overlap indicator) from Marques and Peichel 2017 (*20*), filtered for PVE > 20 and peak interval < 5 Mb); recombination rate (within the TempoPeak and while including 100kb on either side); distance to the nearest 20x recombination hotspot; absolute sequence divergence and estimated allele age (of the entire TempoPeak and of just the 1kb window containing the best ecotypic p-value); other nearby EcoPeaks and TempoPeaks (for both, distance to nearest [in both bp and cM], and number within 2 cM, 5 cM, 10 cM, and on the whole chromosome, for both specific and sensitive call sets); CpG island and TG arrays (binary indicators, counts, and counts/kb); non-exonic conserved sequences (phastCons, total and per kb); ka/ks (for the best gene in the TempoPeak, by actual ka/ks value, unadjusted and adjusted p(ka/ks > 1), defaulting to genome-wide median for intervals not overlapping genes); carrier frequency in the Rabbit Slough marine samples (of the variant with the best CMH p-value); geographic dispersal (both by binary overlap with global EcoPeaks and as maximum distance “as the fish swims” between populations where the haplotype is found); and a randomly generated number for comparison.

In addition to the analysis of these factors individually presented in Fig. 5A, we also asked whether these contained similar or different information from each by examining the speed explained by multiple factors through stepwise forward regression, as in Chaves et al. 2016 (*79*), until the marginal significance of the next best predictor was p > 0.01 (**Fig S25.1**, **Tab S25.1-S25.3**). The full model is able to explain over half the variation in selection speed between the different TempoPeaks. Without considering empirical recurrence, other predictors contribute more, suggesting that recurrence reflects an integration of many of these other predictors.

We further tested how the different contemporary evolution populations predict the speed of evolution in the other contemporary evolution populations and how they compare to empirical ecotypic recurrence across the northeast Pacific (**Fig S25.2**). We observed that ecotypic analysis performs approximately as well as the replicate young populations (and in some cases better). The remaining unexplained differences in speed of selection may therefore be due to lake-specific factors and stochasticity rather than differences between speed of selection and consistency of selection.

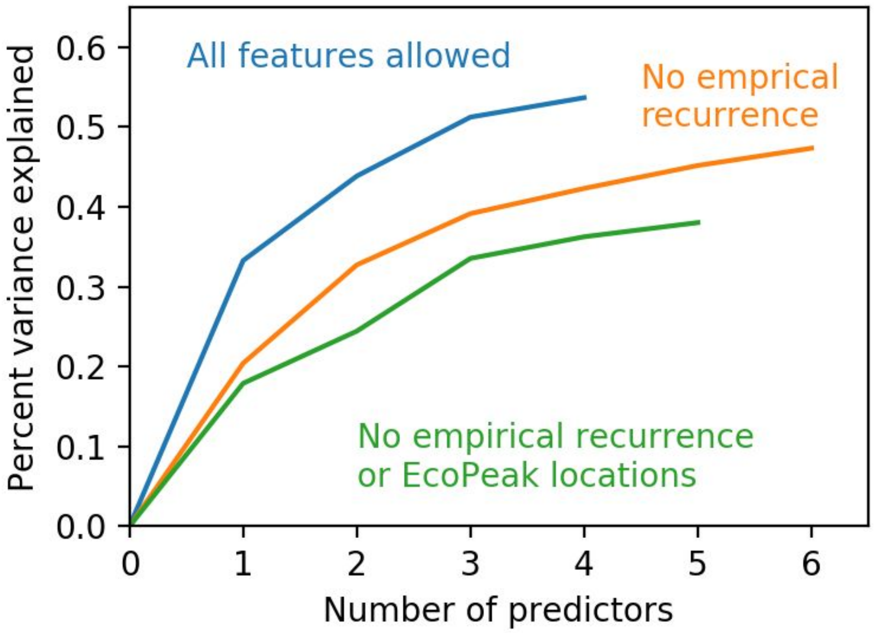


**Fig S25.1:** Stepwise forward regression of features predicting speed of selection in CH+SC+LB, with all predictors allowed(a), without ecotypic recurrence (b), and without ecotypic recurrence or neighboring EcoPeaks (c). The features added as predictors in each step are listed in **Tables S25.1-S25.3** below.

| Order | Predictor added | p sig. improvement | Adj. R squared (cumulative) |
|---|---|---|---|
| 0 | (constant) | - | - |
| 1 | Best SNP in EcoPeak analysis | 6.77e-32 | 0.3327 |
| 2 | Number of specific EcoPeaks within 10cM | 1.77e-14 | 0.4385 |
| 3 | Number of overlapping major QTLs | 4.66e-12 | 0.513 |
| 4 | Best z-score in global EcoPeak analysis | 3.18e-05 | 0.5366 |

**Table S25.1.** Stepwise forward regression analysis and performance of features predicting selection speed, with all features allowed.

| Order | Predictor added | p sig. improvement | Adj. R squared (cumulative) |
|---|---|---|---|
| 0 | (constant) | - | - |
| 1 | Number of specific EcoPeaks within 10cM | 1.04e-18 | 0.2041 |
| 2 | Number of overlapping major QTLs | 3.89e-14 | 0.3272 |
| 3 | TempoPeak size (physical) | 5.36e-09 | 0.3915 |
| 4 | Genic overlap (binary) | 2.19e-05 | 0.4230 |
| 5 | TempoPeak size (genetic) | 3.33e-05 | 0.4517 |
| 6 | Recombination rate | 0.00023 | 0.4733 |

**Table S25.2.** Stepwise forward regression analysis and performance of features predicting selection speed, without empirical recurrence.

| Order | Predictor added | p sig. improvement | Adj. R squared (cumulative) |
|---|---|---|---|
| 0 | (constant) | - | - |
| 1 | Number of overlapping major QTLs | 2.27e-16 | 0.1790 |
| 2 | TempoPeak size (physical) | 1.07e-07 | 0.2444 |
| 3 | TempoPeak size (genetic) | 4.06e-11 | 0.3354 |
| 4 | Recombination rate | 0.00017 | 0.3625 |
| 5 | Genic overlap (binary) | 0.00201 | 0.3802 |

**Table S25.3.** Stepwise forward regression analysis and performance of features predicting selection speed, without empirical recurrence or knowledge of locations of nearby EcoPeaks

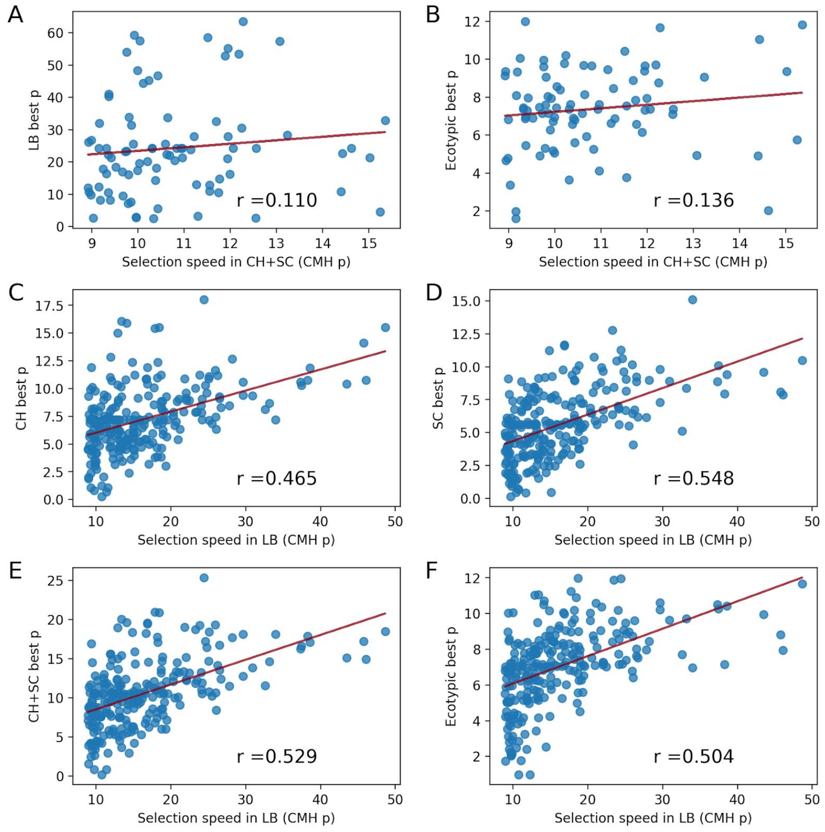


**Figure S25.2:** Comparison of speed predictions based on extant populations or other contemporary evolution populations. Speed of selection in CH+SC TempoPeaks explained by best p-value in region in LB (A) or extant population analysis (B), and speed of selection in CH+SC TempoPeaks explained by best p-value in region in CH (C), SC (D), CH+SC (E), or extant populations (F).

### 26. PREDICTING LOCI OF SELECTION IN STICKLEBACK, FINCHES AND CICHLIDS

We collated data on the following for 2,500 bp non-overlapping windows tiled across the genome for truth data, training labels, and comparative performance purposes: EcoPeak overlap (binary, for northeast Pacific specific EcoPeaks, as training data); significant differentiation in contemporary evolution populations (CH+SC+LB, binary, as truth data); frequency of ecotypic differentiation in established geographic survey populations (best score within the window in the northeast Pacific EcoPeak analysis, by SNP-based log p-value, window-based log p-value, and window-based z-score, as our “gold standard”); speed of differentiation in the contemporary evolution experiments (best CMH p-value within the TempoPeak in all of CH, SC, LB, CH+SC, and CH+SC+LB). We collated data on the following for use as predictors of general evolutionary features without reference to empirical recurrence: genic overlap (gene count and a binary overlap indicator); overlap with major QTLs (both number overlapping the region and a binary overlap indicator) from Marques and Peichel 2017 (*20*), filtered for PVE > 20 and peak interval < 5 Mb); recombination rate (within the window and while including 100kb on either side); absolute sequence divergence and estimated window age (on linear and logarithmic scales); CpG island and TG arrays (binary indicators and counts); non-exonic phastCons; median minor allele frequency in the RS marine samples; and a randomly generated number for comparison.

To incorporate the influence of genomic context without making use of empirical ecotypic recurrence, we created an additional context metric, based on the observation that selected regions typically have low recombination rates and high sequence divergence. This metric quantifies the proportion of nearby regions (within 10 cM) that have higher than median divergence and lower than median recombination.

To select which of the above to use in the combined model to predict loci under selection in CH+SC+LB, individual features were added through stepwise forward logistic regression using the Logit model from the statsmodels.api Python module with default settings. The final model (parameters in **Table S26.1**) was trained entirely on the observations of empirical ecotypic recurrence across the Pacific Northwest, but performance was comparable when trained on subsets of the CH+SC+LB data (**Fig S26.1**). Models were also built and tested with a variety of other model architectures (from sklearn), tested iteratively on each chromosome after training on the rest of the genome to prevent overfitting (**Fig S26.2**); however, we prefered logistic regression because the performance was similar or superior while offering insights into the weighting of the inputs.

For cichlids, we used data from a newly speciated “nyererei-like” and “pundamilia-like” species pair from Python Island and Makobe Island, Lake Victoria (*80*). For our truth sets, we use the previously reported regions of high differentiation (HDR) from Python Island, Makobe Island, and those shared between them (used in Figure 5C). Predictors (in 20kb windows) were recombination rate and absolute sequence divergence between sympatric species pairs. Predictors were then tested separately and in the joint stickleback-derived model (**Fig S26.3**).

For Darwin's finches, we used data from comparisons of 12 pairs species, with the reported islands of genomic divergence as our truth set (*81*). Predictors (in 50kb windows) were absolute sequence divergence (as reported (*81*)), recombination rate, and QTL overlap (as reported (*79*)). Recombination rates were taken from syntenic regions in zebra finch (diverged ~35 Mya), as no recombination maps exist in Darwin's finches but recombination rates are reported to vary little between bird species (*82*). QTLs were treated equally across all pairs of finches regardless of variation between them in the reported trait. Predictors were then tested separately and in the joint stickleback-derived model (**Fig S26.3**). We also ran the model without QTLs to verify that they weren't required to recover well known loci (**Figure S26.4**).

While the composite score does not always perform better than the best predictor in a given species, it will perform across species. Comparing cichlids and Darwin's finches, we note that the best individual feature for predicting selected loci in Darwin's finches is absolute sequence divergence, and that while recombination rates are lower in the islands of divergence, they are not genome-wide outliers like the absolute sequence divergence. In contrast, while cichlids have a few genome-wide absolute divergence outliers in selected regions, the majority of selected regions are outliers by recombination rate with little difference in sequence divergence. This difference may arise from the greatly differing time scales, with the cichlid radiation far younger. Thus, although they both show patterns in sequence divergence and recombination rates, no single individual feature is a reliable predictor in both (**Fig S26.3**).

The association of genomic regions utilized by evolution with low recombination rates and high absolute sequence divergence has been previously reported in a variety of systems, including cichlids (*80*) and Darwin's finches (*81*). However, to our knowledge these associations have always been considered descriptively, while we show here that they can also be used predictively across widely divergent species.

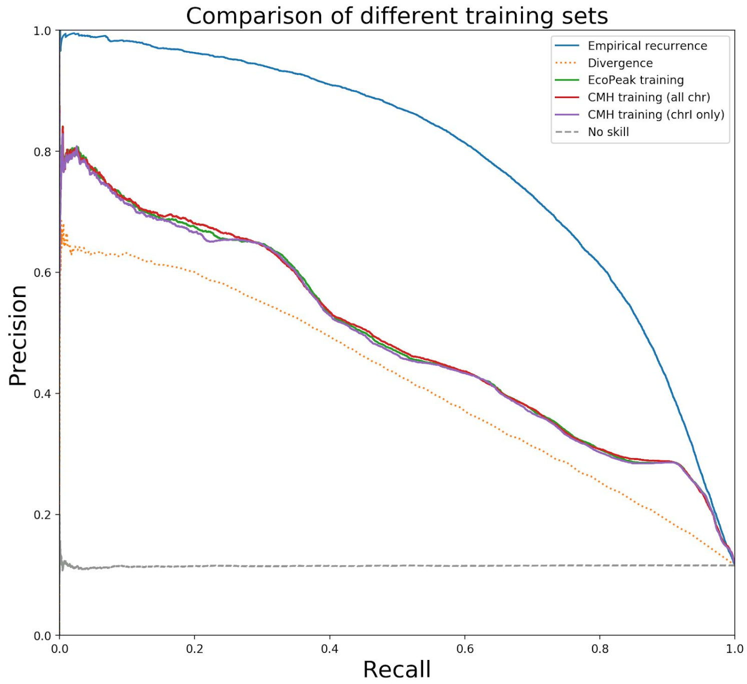


**Figure S26.1:** Results across multiple training sets, using QTL overlap, sequence divergence, recombination rate, and genomic context as input. The model is insensitive to the specific training data.

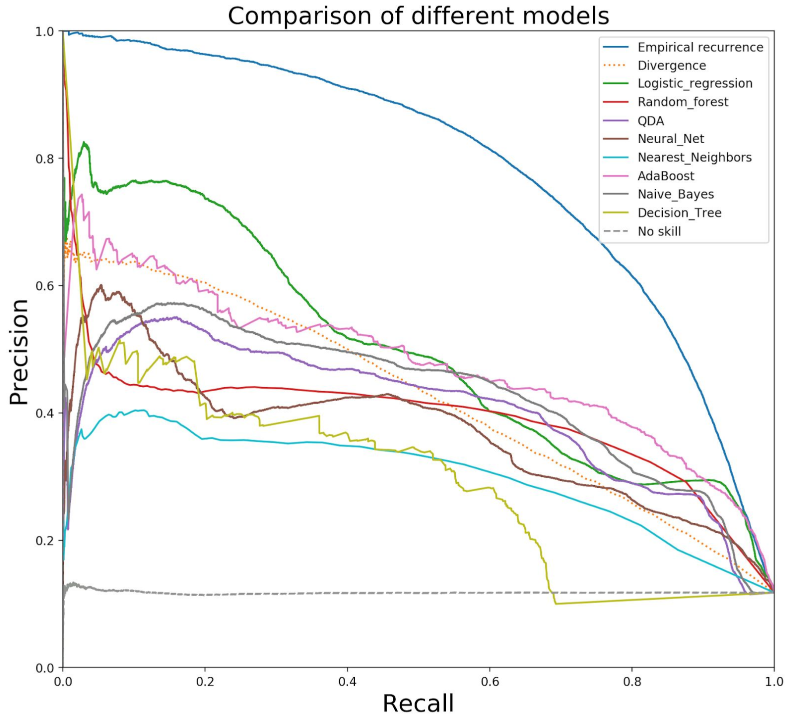


**Figure S26.2:** Results with multiple types of models, compared to empirical recurrence and sequence divergence.

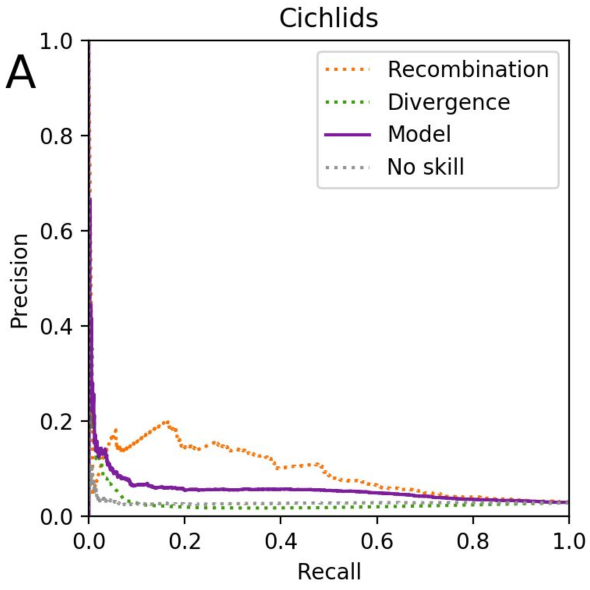


**Figure S26.3:** Results of individual parameters and combined model in cichlids and a representative pair of Darwin's finches (*G. magnirostris* (Genovesa) vs. *G. acutirostris* (Genovesa). Note how most of the precision in cichlids is derived from recombination rates, while most of the precision in finches is derived from sequence divergence.

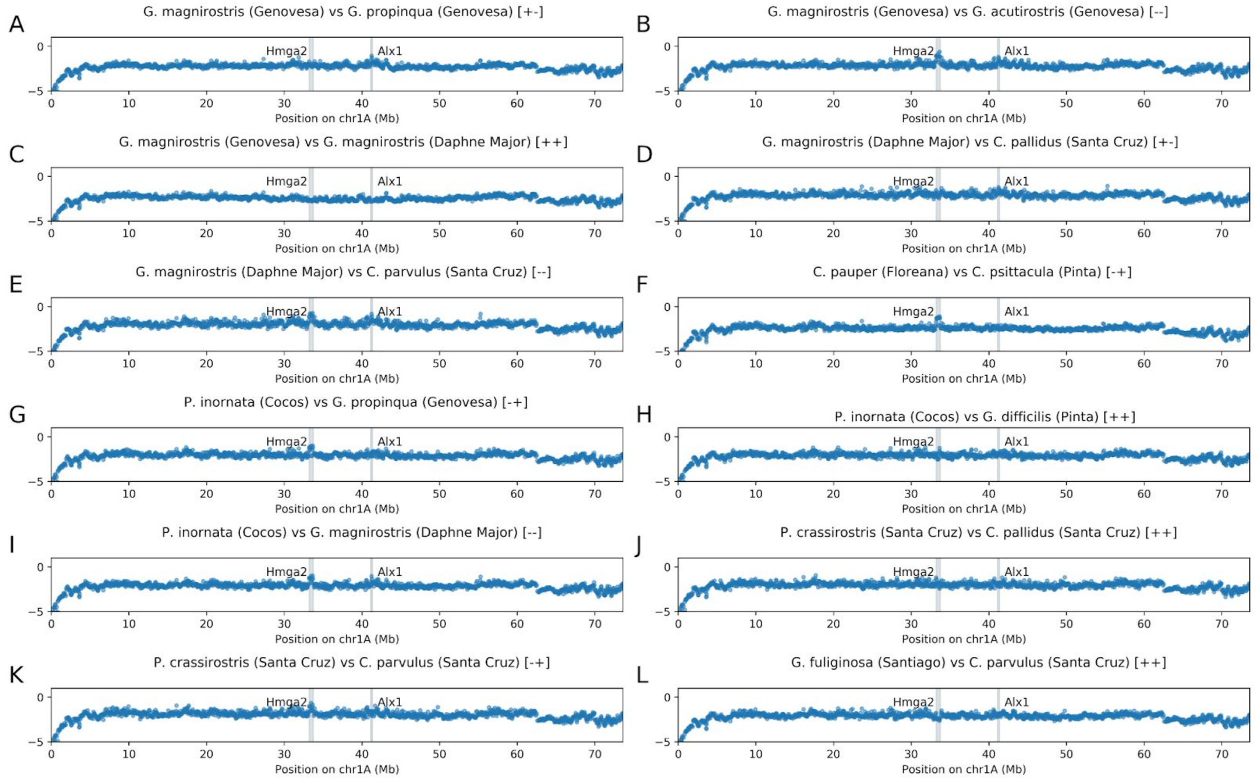


**Figure S26.4: Evolutionary predictions in Darwin's finches based on sequence divergence and recombination rate along chr1A, without QTL input.** The *Hmga2 (83)* and *Alx1 (84)* loci implicated in variation in beak morphology are indicated for reference and are among the top results in several comparisons. The name and island of each species is given, and it is noted whether each pair has the same (+) or different (-) haplogroup at *Hmga2* and *Alx1*, respectively. Species comparisons, sequence divergence, and haplogroup assignments are from Han et al. 2017 (*81*).

| Term | Coefficient (with all terms) | Coefficient (without context) | Coefficient (without QTLs or context) |
|---|---|---|---|
| Constant | -5.3285 | -3.1190 | -2.9112 |
| Divergence | 143.5057 | 174.9383 | 185.5900 |
| Recombination (cM/Mb) | -0.0225 | -0.2031 | -0.2090 |
| QTL overlap | 0.5045 | 0.9617 | - |
| Context (10 cM) | 4.6690 | - | - |

**Table S26.1:** Final model parameters (with and without QTLs and context score)
